# Neofunctionalized RGF pathways drive haustorial organogenesis in parasitic plants

**DOI:** 10.1101/2025.02.09.637280

**Authors:** Maxwell R. Fishman, Anne Greifenhagen, Takanori Wakatake, Anuphon Laohavisit, Ryoko Hiroyama, Sachiko Masuda, Arisa Shibata, Satoko Yoshida, Ken Shirasu

**Author notes:** Nara Institute of Science and Technology, Graduate School of Science and Technology, Ikoma, Nara, Japan (Takanori Wakatake). Institute of Transformative Bio-Molecules, Nagoya, Japan (Anuphon Laohavisit). Corresponding author: Ken Shirasu.

## Abstract

Parasitic plants initiate rapid *de novo* organogenesis of a specialized feeding structure called a haustorium upon contact with their hosts. Currently, little is known about the internal signals regulating haustorium development. Here, we identify root meristem growth factor (RGF) peptides in *Phtheirospermum japonicum* as endogenous inducers of prehaustorium formation. Treatment with specific RGF peptides in the absence of hosts triggered prehaustoria and induced expression of *PjYUC3*, a gene required for auxin biosynthesis and prehaustorium formation. CRISPR-mediated knockouts showed that PjRGFR1 and PjRGFR3, receptors activated by the haustorium specific RGF peptides PjRGF2 and PjRGF5, are essential for prehaustorium formation, revealing functional redundancy. Phylogenic analyses indicate that PjRGF2 is broadly conserved among Orobanchaceae, whereas PjRGF5 appears to have recently evolved through tandem multiplication and neofunctionalization. Our findings establish RGF peptides and their corresponding receptors as critical components of haustorium developmental signaling and provide insights into the evolutionary trajectories that shape plant parasitism.

**Teaser:** Plant peptide hormones regulate and induce the parasitic plant specialized organ for connecting to and feeding from the host.

## Introduction

Parasitism has independently evolved 12 to 13 times from autotrophic angiosperms and has resulted in around 1% of angiosperms being either root or shoot parasites (*1, 2*). Parasitic plants occur as obligate non-photosynthetic holoparasites or obligate photosynthetic hemiparasites that both require a host to survive or as facultative parasites that can survive without a host (*3*). In order to parasitize hosts, parasitic plants form a direct connection with host plants through a specialized organ called a haustorium (*2*). Haustoria arise *de novo* upon host contact and progress through specific developmental stages, starting with the prehaustorium, followed by the formation of intrusive cells that penetrate the host tissue, and culminating in the establishment of a vascular connection (*2*). This intimate linkage enables parasitic plants to secure water, minerals, and nutrients from their hosts, a process essential for survival of obligate parasites (*1–3*).

The family Orobanchaceae consists predominantly of root parasites (*4*), encompassing agriculturally devastating obligate parasites in the genera *Orobanche, Phelipanche, Alectra*, and *Striga*, as well as comparatively benign facultative parasites in *Pedicularis* and *Phtheirospermum* (*5, 6*). Most members of this family initiate prehaustoria formation upon the perception of lignin- and phenolic-based host-derived haustorium-inducing factors (HIFs), even in the absence of a host (*7, 8*). The first identified HIF was 2,6-dimethoxy-*p-*benzoquinone (DMBQ) (*9*). DMBQ is a lignin-derived quinone that induces prehaustoria in *Striga* and most other hemiparasitic plants in Orobanchaceae (*10*). The obligate holoparasites *Phelipanche* and *Orobanche* develop prehaustoria in response to cytokinins, sphaeropsidones, and very high concentrations of DMBQ (*11, 12*). DMBQ has become a central tool to dissect the early signaling pathway underlying haustorium development, particularly in the facultative parasites *Phtheirospermum japonicum* and *Triphysaria versicolor,* and in the obligate hemiparasite *Striga hermonthica* (*13–17*). DMBQ, and other similar quinone-based HIFs, are first sensed by CARD1-LIKE (CADL) receptor kinases (*14*), triggering Ca^2+^ signaling in roots and activating a MAPK cascade. These early signals result in the differential regulation of hundreds of genes, leading to the accumulation of reactive oxygen species (ROS) and the *de novo* biosynthesis of auxin through the up-regulation of YUCCA3 MONOOXYGENASE (YUC3) specifically at the site of haustorium initiation (*13, 15, 18*). Both ROS and auxin are essential for haustorium development; however, the signaling pathways linking the initial CADL signaling cascade to the downstream factors such as YUC3 are largely unknown.

ROOT GROWTH MERISTEM FACTOR (RGF) peptides (also referred to as GLV or CLEL peptides) were initially identified for their role in maintaining the root apical meristem (RAM) in *Arabidopsis thaliana* (from here on referred to as Arabidopsis) but also regulate lateral root elongation and spacing, as well as root gravitropism (*19–22*). They consist of a family of sulfated plant peptide hormones present in all land plants outside of hornworts and mosses (*23*). They serve as ligands for the RGF RECEPTOR (RGFR)/RGF INSENSITIVE (RGI) leucine-rich repeat receptor-like kinases where the conserved tyrosine, proline, and histidine/asparagine in the mature peptide are critical for interacting with RGFRs or their co-receptors (*24–26*). The regulatory role of RGF peptides in maintaining the RAM is partially accomplished by stabilizing the AP2-type transcription factor PLETHORA2 (PLT2) through the production of superoxide radical (O_2_^-^) (*27*). Moreover, RGF peptides selectively regulate a subset of auxin-responsive genes and mobilize PIN proteins to specify auxin localization (*22, 28*). Given the importance of auxin and ROS during haustorium development in parasitic plants, RGF peptides are plausible regulators of haustorium signaling.

In this current study we identified three RGF peptides in *P. japonicum* (PjRGFs) that are up-regulated following DMBQ treatment. Treatment of these DMBQ-induced PjRGF peptides led to an up-regulation of *PjYUC3* and induced prehaustoria in *P. japonicum*. Two of these PjRGF peptides are specifically expressed in the prehaustorium, reflecting the localization patterns of a *P. japonicum* PLT2 homologue (PjPLT) and O_2_^-^. Further analysis of RGF peptides in several Orobanchaceae species uncovered a single conserved haustorium-specific RGF peptide that also induces prehaustoria in *Striga hermonthica*. Genetic analysis further revealed that knockouts of two PjRGFRs reduced the number of DMBQ-induced prehaustoria in *P. japonicum* hairy roots and one PjRGF peptide regulates cell division at the apex of the prehaustorium. Taken together, we established RGF peptides in Orobanchaceae as endogenous prehaustorium inducers, placing RGF signaling downstream of DMBQ and upstream of *YUC3* in the haustorium development pathway. These data suggest that parasitic plants neofunctionalize the RGF signaling pathway for *de novo* organogenesis, contributing to the evolution of plant parasitism.

## Results

### RGF peptides are up-regulated during haustorium induction

To identify signaling components activated by DMBQ perception, we analyzed RNA-seq data from *P. japonicum* seedlings. Since subtilase encoding-genes, which code for apoplastic proteases, are important for haustorium development (*29*), we searched for apoplastically-cleavable peptide hormone-encoding genes that were up-regulated in response to DMBQ treatment. From this approach, we identified a gene encoding an RGF peptide, from herein called *PjRGF1*, which was induced following DMBQ treatment (Fig. S1A, B; Supplementary Table S1). Although only four RGF peptide-encoding genes were initially annotated in the *P. japonicum* genome (*30*), additional mapping and annotation revealed another 13 putative *RGF* genes, 9 of which were expressed (Fig. 1A; Fig. S2A-C; Supplemental Table S2).

**Fig. 1.**
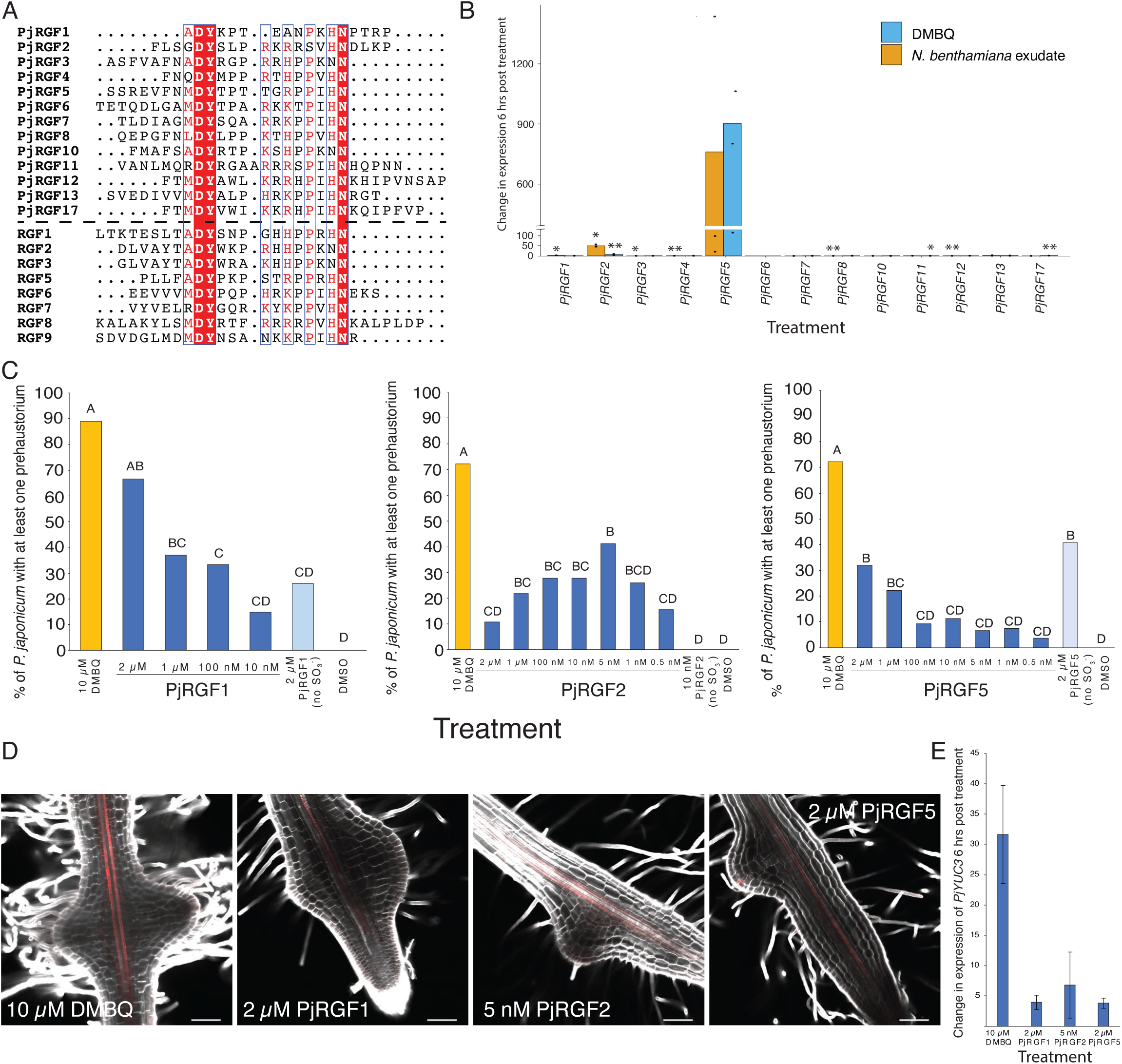
PjRGF peptides induce prehaustoria. A) Multiple sequence alignment of the C-terminus of the identified RGF peptides in *P. japonicum* and eight of the eleven RGF peptides in Arabidopsis. B) Change in expression of the identified PjRGF peptides six hours post 10 µM DMBQ treatment or *N. benthamiana* treatment as compared to treatment with DMSO. Each dot represents the fold-change per biological replicate. *PjRGF5* was consistently up-regulated however the high-variability between replicates caused the p-value to be > 0.1. Student’s T-test was used to determine statistical significance. * represents p-value < 0.05 **represents p-value < 0.1. C) Percent of *P. japonicum* forming at least one prehaustorium two day post treatment with DMBQ, PjRGF1, PjRGF2, or PjRGF5. One-way ANOVA followed by Tukey HSD was used for identifying statistical significance. D) Confocal images of *P. japonicum* two days after treatment with 10 µM DMBQ, 2 µM PjRGF1, 5 nM PjRGF2, or 2 µM PjRGF5 stained with calcofluor white (white) and basic fuschin (red). Scale bars = 75 µm. E) Change in expression of *PjYUC3* six hours post treatment with 10 µM DMBQ, 2 µM PjRGF1, 5 nM PjRGF2, or 2 µM PjRGF5.

To determine which of these expressed *PjRGFs* were HIF responsive, RT-qPCR was performed on cDNA from *P. japonicum* roots treated for six hours with either DMBQ or *Nicotiana benthamiana* exudate. These data showed that *PjRGF1, PjRGF2*, and *PjRGF5* were up-regulated following DMBQ treatment, whereas only *PjRGF2* and *PjRGF5* were induced by *N. benthamiana* exudate (Fig. 1B). Notably, in *S. hermonthica,* the *RGF2* homolog (*ShRGF2*) was also up-regulated six hours post DMBQ treatment (Fig. S3A, B), suggesting that RGF peptides may play a conserved role in the haustorium developmental signaling pathway across multiple parasitic species within Orobanchaceae.

### RGF peptides induce prehaustoria in parasitic plants

RGF peptides induce a variety of root-related phenotypes in autotrophic plants (*19, 28*). Here, surprisingly, application of PjRGF1, PjRGF2, or PjRGF5 to *P. japonicum* induced prehaustoria (Fig. 1C, D). Unsulfated PjRGF1 and PjRGF2 peptides showed only a weak effect but unsulfated PjRGF5 possessed prehaustorium induction activity (Fig. 1C). Similarly, ShRGF2 treatment of *S. hermonthica* also induced prehaustoria (Fig. S3C, D). These induced structures were morphologically similar to DMBQ-induced prehaustoria in both *P. japonicum* and *S. hermonthica*, and their induction rate was dose dependent in *P. japonicum* (Fig. 1C, D; Fig. S3C. D; S4A). The overall efficacy of the PjRGF peptide treatments was lower than that of DMBQ and varied among individual peptides. Both PjRGF1 and PjRGF5 reached maximal induction at 2 µM, while PjRGF2 was most effective at 5 nM (Fig. 1C). Higher concentrations of PjRGF2 sometimes resulted in nonspecific cell division in the epidermis and cortex (Fig. S4B). Prehaustorium formation begins with controlled cell division in the transition zone (*7*), so nonspecific cell division likely reduced the number of prehaustoria formed and suggests that *P. japonicum* is particularly sensitive to PjRGF2. Overall, these findings demonstrate that PjRGF1, PjRGF2, PjRGF5, and ShRGF2 act as ligands capable of inducing prehaustoria.

*PjYUC3* up-regulation was previously found to be downstream of HIF treatment, functioning as an important component during prehaustorium development (*13*). Consistent with this observation, treatment of *P. japonicum* roots with PjRGF1, PjRGF2, or PjRGF5 induced *PjYUC3* expression six hours after treatment (Fig. 1E), though to a lesser extent than DMBQ. In keeping with the previous findings that auxin accumulates at the apex of the *P. japonicum* prehaustorium, where *PjYUC3* is expressed (*13*), a DR5 reporter revealed auxin localized in the epidermis at the apex of the prehaustorium upon treatment with PjRGF2 or DMBQ, or upon association with host roots (Fig. 2A). These data suggest that the prehaustorium development pathway triggered by PjRGF peptides converges on PjYUC3 mediated auxin biosynthesis, echoing the pathway induced by classic HIFs.

**Fig. 2.**
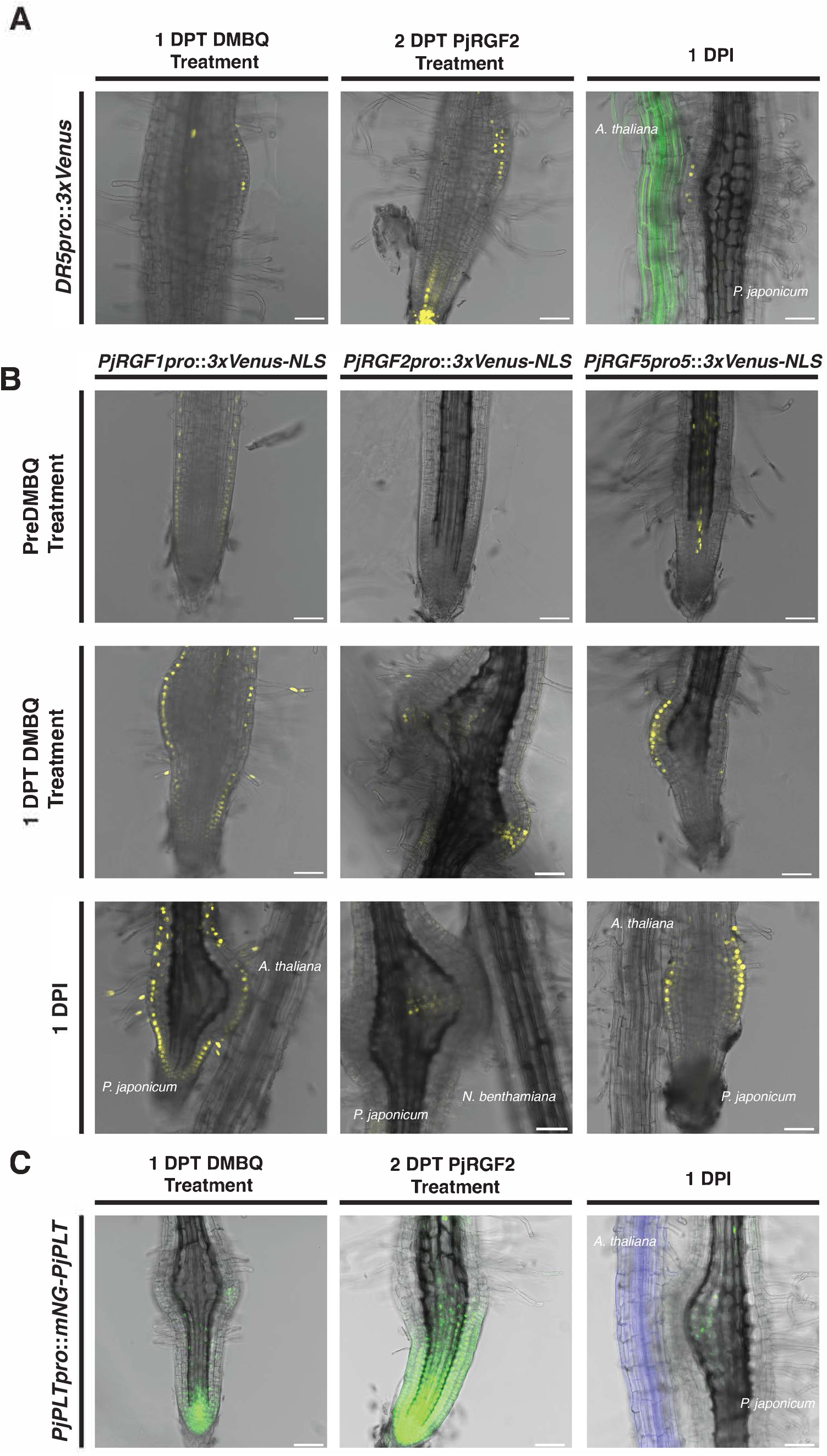
Hausotrial expression dynamics of *PjRGF* peptides and downstream signaling events. A) Expression of *3x-Venus-NLS* driven by the *DR5* promoter following 1 day post treatment (DPT) 10 µM DMBQ, 2 DPT 5 nM PjRGF2, and at 1 one day post infection (DPI). *DR5* expression, which correlates with auxin localization, occurs at the epidermal cells at the apex of the prehaustorium in all treatments. B) Representative confocal images of transcriptional reporters for *PjRGF1* (*PjRGF1pro*), *PjRGF2* (*PjRGF2pro*), and *PjRGF5-5* (*PjRGF5pro*5) prior to 10 µM treatment of DMBQ, 1 DPT 10 µM DMBQ, and 1 DPI. Both *PjRGF2pro* and *PjRGF5pro* are localized to the prehaustorium while *PjRGF1pro* is not. C) Localization of mNG-PjPLT after 1 DPT 10 µM DMBQ, 2 DPT 5 nM PjRGF2, and at 1 DPI. mNG-PjPLT localizes within the prehaustorium of *P. japonicum* upon all treatments but the localization of mNG-PjPLT slightly differs. Scale bars are 75 µm. Fluorescent Arabidopsis is expressing *3SS::LT6-GFP*.

### *PjRGF2* and *PjRGF5* expression localizes to the prehaustorium

Because RGF peptides typically act close to their site of expression (*31*), we investigated whether *PjRGF1, PjRGF2*, or *PjRGF5* were specifically expressed in the prehaustorium following DMBQ treatment. We generated transcriptional reporter lines expressing *3x-Venus* tagged with a nuclear localization sequence (NLS) under the control of each *PjRGF* promoter (*PjRGF1/2/5pro*). At one day post DMBQ treatment and one day post inoculation with a host root, *PjRGF2pro* and *PjRGF5pro* were clearly expressed in the prehaustorium, while *PjRGF1pro* was expressed in the epidermis throughout the root, regardless of prehaustorium formation (Fig. 2B). Prior to prehaustorium formation, *PjRGF2pro* showed no expression, while *PjRGF5pro* was observed in the cortex and stele. Given that *PjRGF2* and *PjRGF5* were up-regulated at six hours post DMBQ treatment (Fig. 1B), we hypothesized that their expression might localize at this early time point to the transition zone where prehaustoria form. Indeed, at six hours post DMBQ treatment, *PjRGF2pro* expression was observed around the transition zone, and *PjRGF5pro* showed increased expression relative to untreated roots (Fig. S5; Fig. S6). Collectively, these data indicate that PjRGF2 and PjRGF5 are prehaustorium-specific RGF peptides likely contributing to haustorium developmental signaling, while other PjRGF peptides may not play a major role.

### PjPLT localizes to the prehaustorium

If RGF peptides indeed influence haustorium development, a recognizable marker of their activity should be detectable in the prehaustorium. In Arabidopsis, RGF peptides induce O_2_^-^ production that then stabilizes PLT2 (*27*). Likewise, O_2_^-^ was previously shown to accumulate in the prehaustoria of *S. hermonthica* (*18*). Consistent with these findings, nitro blue tetrazolium staining showed that O_2_^-^ also accumulated in the prehaustorium of *P. japonicum* (Fig. S7). These observations suggest a PLT transcription factor might be involved in prehaustorium development. Among six predicted PLT homologs in *P. japonicum*, only four contain two AP2 domains characteristic of PLTs (Fig. S8A). The closest homolog to Arabidopsis PLT2 is Pjv1_00021464, which was named *PjPLT* (Fig. S8B). Under its native promoter, mNeonGreen-tagged PjPLT (mNG-PjPLT) localized to the RAM prior to DMBQ treatment or host inoculation (Fig. S9A). At one day post-DMBQ treatment, mNG-PjPLT was observed in the RAM, the transition zone, and the outer cell layers at the apex of the prehaustorium. mNG-PjPLT also localized within the prehaustorium at 1 DPI, shifting to the center of the prehaustorium (Fig. 2C). This pattern appears to be regulated post-translationally, as a transcriptional reporter for *PjPLT* was expressed broadly in the meristem, transition, and elongation zones as well as the developing prehaustorium, both at 1 DPT DMBQ and at 1 DPI (Fig. S9B). Given that PjRGF peptides induce prehaustoria and RGF peptides stabilize PLT2 in *Arabidopisis*, we anticipated that upon exogenous PjRGF treatment PjPLT would localize similarly to what is observed under DMBQ treatment. Indeed, mNG-PjPLT was present in the prehaustorium following PjRGF2 treatment (Fig. 2C). However, mNG-PjPLT was also found throughout an elongated meristemic zone and in the interior of the prehaustorium (Fig. 2C), diverging slightly from its localization under DMBQ treatment or at 1 DPI. Nevertheless, both DMBQ and PjRGF2 guide mNG-PjPLT into the prehaustorium, indicating that small differences in localization still result in successful prehaustorium formation.

### PjRGF5 is a novel RGF peptide that underwent a series of tandem duplication events in *P. japonicum*

RGF peptides commonly function redundantly, so it is unsurprising that *PjRGF2* and *PjRGF5* are both expressed in the prehaustorium. Analysis of a *P. japonicum* genome assembly produced from PacBio Continuous Long Reads revealed a series of tandem duplications of *PjRGF5* across five contigs (Fig. 3A). At least five nearly identical copies of *PjRGF5* and six *PjRGF5* pseudogenes reside in these duplications, all of which are associated with LTR, TIR, and Helitron transposable elements (TE), with *PjRGF5-7, PjRGF5-8,* and *PjRGF5-9* having a TIR TE inserted in the intron. These TEs may have driven the duplication events. In addition, subcloning of *PjRGF5* cDNA uncovered yet another *PjRGF5* duplicate (Fig. S10). Several of the contigs carrying *PjRGF5* duplicates are quite small, indicating a poorly assembled region (Fig. 3A), and suggest further copies might remain unassembled. The sequences of the *PjRGF5* replicates are highly conserved, typically differing by only one or two single nucleotide polymorphisms (SNPs) in the sequences encoding the non-conserved region of the prepropeptide (Fig. S11A,B). For example, two copies, designated PjRGF5-2a and PjRGF5-2b, are identical. Multiple sequence alignment revealed that three annotated *PjRGF5* promoters (*PjRGF5-1pro, PjRGF5-2apro*, and *PjRGF5-2bpro*) are practically identical, while the other promoter, labelled PjRGF5-5pro, shares a high degree of similarity up to 1555 bp upstream of the start codon (Fig. S12). In agreement with these results, spatial expression patterns driven by the *PjRGF5-1pro, PjRGF5-2bpro* (Fig. S13), and *PjRGF5-5pro* (Fig. 2B) visualized using 3x Venus-NLS were observed in the same epidermal region of the prehaustorium. These data suggest that multiple duplicated genes together drive the production of bioactive PjRGF5 in the prehaustorium.

**Fig. 3.**
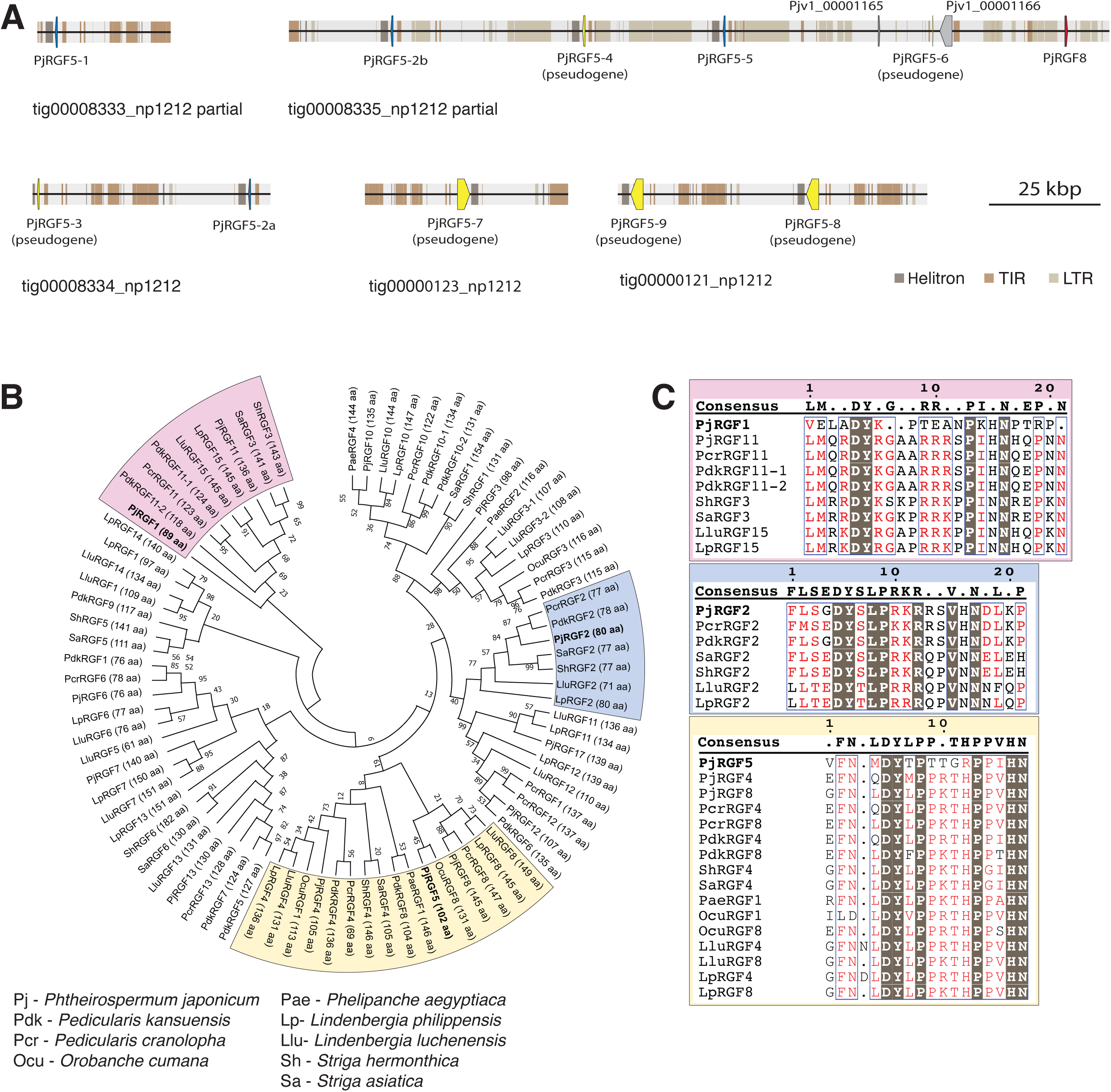
Genomic landscape of haustorium specific RGF peptides. A) Illustration of the contigs where *PjRGF5* duplications occur. Tandem duplications of *PjRGF5* occur on several contigs. Only small sections of tig00008335_np1212 and tig00008333_np1212 are shown in the illustration while the entirety of the other contigs are represented in the illustration. Blue *PjRGF5* duplications encode viable genes and yellow *PjRGF5* duplications encode likely pseudogenes. *PjRGF8* is shown in red. The short length of tig00008334_np1212, tig00000123_np1212, and tig00000121_np1212 suggest a potentially complicated area of the genome to assemble. Colored segments of each contig represent areas where Helitrons, TIRs, and LTRs occur. *PjRGF5-7, PjRGF5-8*, and *PjRGF5-9* all have a TIR in the intron. B) Maximum-likelihood phylogenetic tree with a Bootstrap of 100 made using identified and annotated RGF prepropeptides encoded in the genomes of several species found in Orobanchaceae. Clades that include PjRGF1, PjRGF2, and PjRGF5 are colored in pink, blue, and yellow, respectively. PjRGF1/2/5 are in bold. C) Multiple sequence alignment of the C-terminus of the RGF prepropeptides found in clades with PjRGF1/2/5. Only the PjRGF2 mature peptide sequence is conserved.

The extensive duplication of *PjRGF5* points to a potential importance for haustorium development in *P. japonicum* and possibly other parasitic Orobanchaceae species. A phylogenetic analysis of RGF prepropeptides from *P. japonicum,* six parasitic Orobanchaceae species and two non-parasitic Orobanchaceae species showed that PjRGF1, PjRGF2, and PjRGF5 each occupy distinct clades (Fig. 3B, Supplemental Table S2) (*6, 32–35*). Surprisingly, the mature peptide for PjRGF1 and PjRGF5 did not align with the other RGF peptides within their clades, while the mature peptide of PjRGF2 was conserved across *Lindenbergia, Pedicularis*, and *Striga* (Fig. 3C, Supplemental Table S2). The only exceptions were the obligate holoparasites *Orobanche cumana* and *Phelipanche aegyptiaca,* which lack a PjRGF2 homolog. These data suggest that PjRGF1 and PjRGF5 may be recent acquisitions in *P. japonicum*, while PjRGF2 appears to be a more ancient, conserved haustorium-related RGF peptide in parasitic Orobanchaceae.

### Several *PjRGFRs* are expressed in the prehaustorium and PjRGFR1, PjRGFR3, PjRGFR4 are activated by PjRGF2 and PjRGF5

The receptors for RGF peptides, known as RGFRs/RGIs, were first identified in Arabidopsis (*24–26*). Similar to RGF peptides, RGFRs are conserved throughout land plants. A phylogenetic analysis of RGFRs from 360 species reveal three main clades and two sub-clades (Fig. S14) (*6, 34, 35*). *P. japonicum* encodes six RGFRs, more than most sequenced parasitic Orobanchaceae species (Supplemental Table S3). RT-qPCR analysis of cDNA extracted from *P. japonicum* root tips six hours post DMBQ or *N. benthamiana* exudate treatments showed that *PjRGFR3* expression decreased significantly only in response to DMBQ treatment and it is possible that *PjRGFR3* downregulation was a response to PjRGF2/5 signaling downstream of HIF perception (Fig. S15). However, spatial expression analysis using *PjRGFR promoter*-*3x Venus-NLS* reporter lines (*PjRGFRpro*) showed that *PjRGFR3pro* was not expressed in the prehaustorium (Fig. 4A). Instead, *PjRGFR1pro, PjRGFR2pro, PjRGFR4pro* and *PjRGFR5pro* each displayed some haustorium specific expression pattern, particularly when a host root was present, implying that multiple *P. japonicum* RGFRs likely coordinate prehaustorium development.

**Fig. 4.**
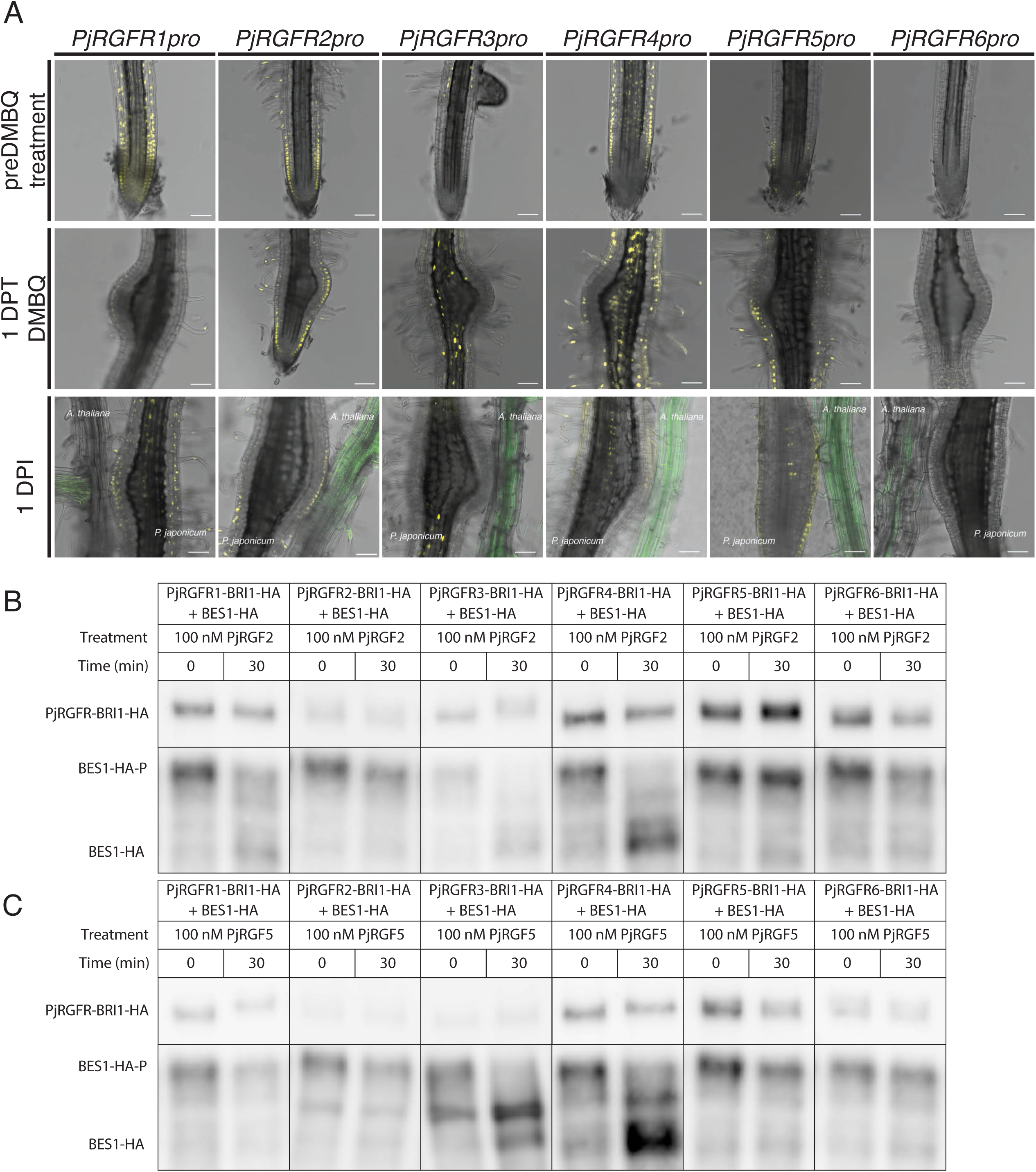
PjRGFR expression dynamics and their interactions with haustorium specific PjRGFs. A) Representative confocal images of transcriptional reporters for *PjRGFR1-6* (*PjRGFRpro*) before treatment with 10 µM DMBQ, 1 DPT 10 µM DMBQ treatment, and at 1 DPI. The spatial expression pattern of *PjRGFR1/2/4/5* promoters at 1 DPI localize to the prehaustorium. These expression patterns are less clear in roots at 1 DPT 10 µM DMBQ save for the spatial expression pattern of *PjRGFR2pro*. At 1 DPT 10 µM DMBQ and 1 DPI, *PjRGFR2pro* is expressed in the epidermal cells of the prehaustorium. Scale bars = 75 µm. Arabidopsis is expressing *35S::LT6-GFP*. B,C) *PjRGFR-BRI1-HA* chimera and *BES1-HA* were expressed in *N. benthamiana*. Treatment with B) 100 nM PjRGF2 resulted in dephosphorylation of BES1-HA when PjRGFR1/3/4-BRI1-HA were expressed and with C) 100 nM PjRGF5 resulted in dephosphorylation of BES1-HA when PjRGFR3/4-BRI-HA were expressed.

Given the difference between RT-qPCR data and promoter driven expression patterns, we next assessed which PjRGFRs directly bind to PjRGF2 and PjRGF5. As chimeric LRR-RLK receptors have previously aided in probing ligand-receptor interactions (*36*), we generated PjRGFR-BRI1 chimeras to evaluate their activation by PjRGF2 or PjRGF5 (Fig. S16). A shift in the BES1 transcription factor from its phosphorylated to dephosphorylated state served as a readout of receptor activation. In *N. benthamiana* leaves co-expressing the *PjRGFR-BRI1-HA* variants and *BES1-HA*, treatment with 100 nM PjRGF2 for 30 min activated PjRGFR1-BRI1-HA, PjRGFR3-BRI1-HA, and PjRGFR4-BRI1-HA (Fig 4B), while 100 nM PjRGF5 for 30 min activated PjRGFR3-BRI1-HA, and PjRGFR4-BRI1-HA (Fig 4C). These results suggest that PjRGFR1, PjRGFR3, and PjRGFR4 function as primary receptors in the prehaustorium signaling cascade of *P. japonicum*.

### PjRGFR1 and PjRGFR3 are positive regulators of prehaustorium formation

To determine the role of PjRGF2 and the PjRGFRs responsive to PjRGF2 and PjRGF5, we used the hairy root CRISPR system, a method employed in *P. japonicum* and other plant species with success (*37, 38*). CRISPR-zCas9i constructs targeting *PjRGF2, PjRGFR1, PjRGFR3*, and *PjRGFR4*, were generated with a RUBY reporter for transformed roots (Fig. 5A, Fig. S17). The frequency of knockout mutations ranged from 19% to 45% depending on the target gene (Supplemental Tables S4, S5). Transgenic roots expressing RUBY were treated with 10 µM DMBQ for two days, and the presence of prehaustoria was evaluated. Roots were classified as lacking prehaustorium (including minor cell division), displaying a normal prehaustorium, showing aberrant cell division in the prehaustorium, or exhibiting an aborted prehaustorium (Fig. 5A). Only *PjRGFR1* and *PjRGFR3* knockouts significantly reduced the proportion of roots forming prehaustoria compared to the control (Fig. 5B). In contrast, *PjRGFR4* knockouts had no significant effect on DMBQ-induced prehaustorium formation. Interestingly, although *PjRGF2* knockouts did not reduce the number of prehaustoria, they did develop an unusual prehaustorium phenotype more frequently than the control. These abnormal prehaustoria were pointed at the tip rather than rounded like a normal prehaustorium in a way that was reminiscent of an ectopic root (Fig. 5A). However, confocal microscopic images showed that this pointed structure did not resemble a root tip. Rather, it was capped by two large cells that were divided directly at the tip of the abnormal structure (Fig. S18), suggesting *PjRGF2* regulates cell division in the haustorium. Taken together, these results indicate that two PjRGFRs are crucial for prehaustorium formation and suggest that multiple PjRGF peptides likely function redundantly, given that a PjRGF2 single knockout did not alter number of prehaustoria.

**Fig. 5.**
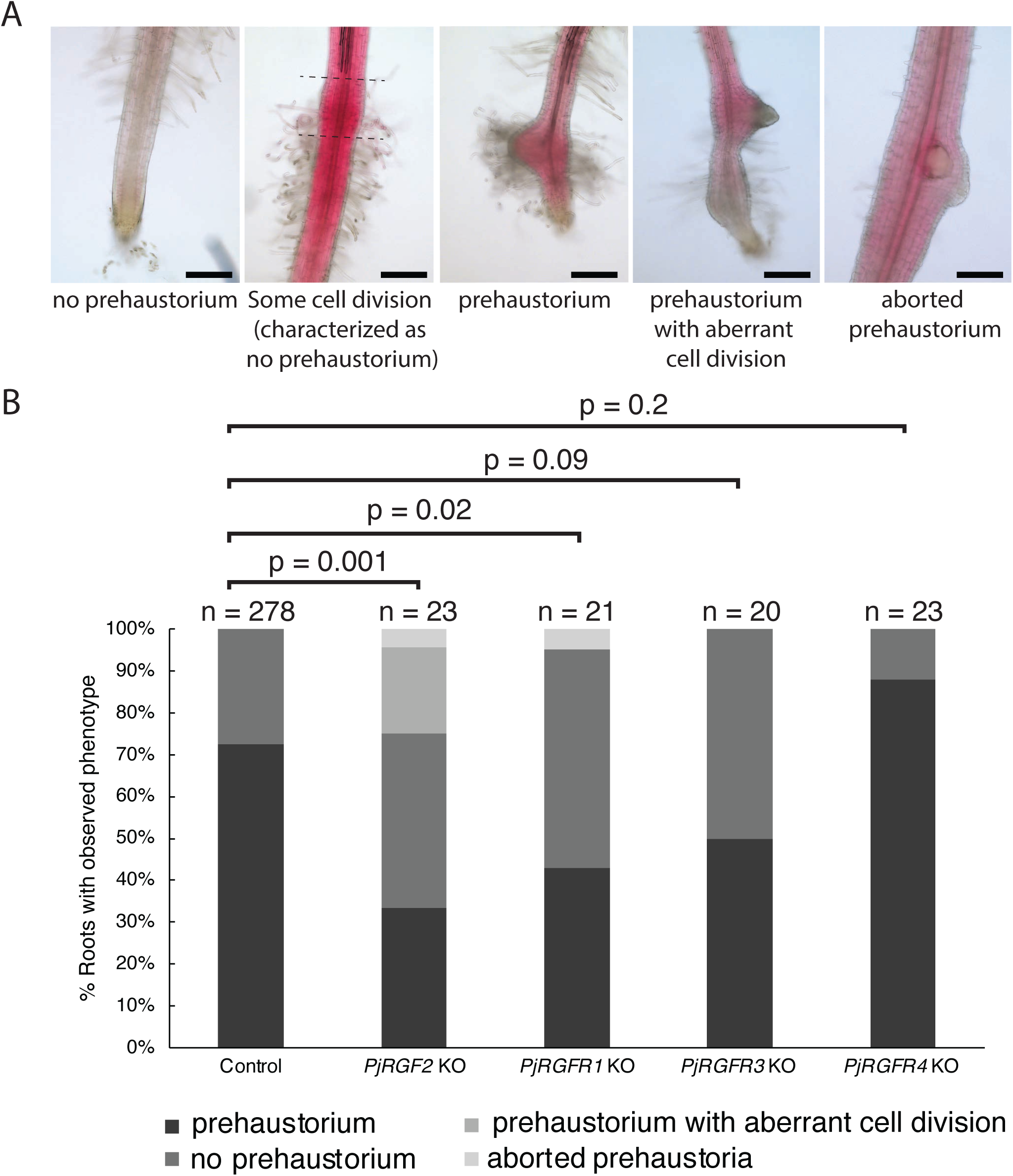
PjRGF signaling pathway is necessary for prehaustorium induction. A) Microscopic images of *P. japonicum* hairy roots overexpressing RUBY exhibited four distinct phenotypes following 10 µM DMBQ treatment. These phenotypes were characterized as no prehaustorium, some cell division occurred (scored as no prehaustorium), prehaustorium, prehaustorium with aberrant cell division, and aborted prehaustorium. Roots that exhibited some cell division were characterized as having no prehaustorium. Cell division can be seen to be occurring in the area in between the dotted lines in the representative photo for roots with some cell division. Scale bar = 200 µm B) Graph showing the percent of *P. japonicum* hairy roots exhibiting the previously described phenotypes. *PjRGF2* KO, *PjRGFR1* KO, and *PjRGFR3* KO hairy roots showed a significant difference in normal prehaustorium development as compared to control hairy roots while *PjRGFR4* KO hairy roots did not show a significant difference in prehaustorium development. p-values were calculated using a Mann-Whitney U test.

## Discussion

Previous insights into the internal signals driving haustorium development beyond the initial HIFs were lacking. Here, we show that RGF peptides function as endogenous signals that positively regulate haustorium formation following DMBQ treatment and are also capable of inducing prehaustoria in *P. japonicum* and *S. hermonthica* in the absence of HIFs. Another recently described endogenous peptide in *P. japonicum*, PjCLE1, can also induce prehaustoria (*39*); however, PjCLE1 is not expressed in the prehaustorium phase and instead shows up later, after xylem bridge formation. In contrast, as expected of peptide hormones involved in prehaustorium development, *PjRGF2* and *PjRGF5* are expressed at early timepoints following HIF perception (Fig. 1B). Peptide hormones often act redundantly, as demonstrated by RGF, CLE, and TOLS peptides, which all negatively regulate lateral root development in Arabidopsis (*40–42*). It is therefore possible that multiple peptide hormones regulate haustorium development in parasitic plants and each can independently induce prehaustoria in *P. japonicum*. Further work is needed to establish whether additional peptide hormones participate in the *P. japonicum* prehaustorium induction pathway.

Earlier work has shown that *de novo* biosynthesis of auxin through PjYUC3 is essential for haustorium formation (*13*). Consistent with this role, treatment of *P. japonicum* with PjRGF1, PjRGF2, and PjRGF5, like DMBQ, up-regulated *PjYUC3* in root tips (Fig. 1D). This differs from RGF peptides in Arabidopsis as they are linked to auxin signaling through PIN protein mobilization rather than local auxin biosynthesis (*21, 22, 28*). This does not rule out the possibility that PjRGF peptides also influence PjPIN distribution in the haustorium, as there is overlap with the spatial expression pattern of *PjRGF2pro* and *PjRGF5pro* with PjPIN1 and PjPIN2 (Fig. 2B) (*43*). However, PIN inhibition in *P. japonicum* does not impact prehaustorium formation (*43*). Therefore, it is likely that cell-specific up-regulation of *PjYUC3* is the main mechanism through which PjRGFs induce prehaustoria. The RGF peptide pathway in *P. japonicum*, while operating under the distinct context of haustorium formation, likely diverges from Arabidopsis beyond PLT2 stabilization given the fact that PjRGF2 extends the gradient of PjPLT in the RAM in a way reminiscent of RGF peptides in Arabidopsis (Fig. 2C) (*19, 27*). A similar divergence was observed between CARD1-LIKE-based DMBQ signaling in *P. japonicum* and CARD1-based DMBQ signaling in Arabidopsis (*14*). Therefore, RGF peptide signaling represents another evolutionary innovation in parasitic plants. Further characterization of PjRGF peptides should illuminate additional roles these signals play and the genes they regulate during haustorium development.

The conservation of RGF peptides in plants, coupled with their unique usage in *P. japonicum* and *S. hermonthica* highlights the need to explore how genes evolve in connection with parasitism. Although our current understanding of plant parasitism is limited, the multiplication and neofunctionalization of KAI2d/ShHTL strigolactone receptors from KAI2 in Orobanchaceae serve as a notable precedent (*44, 45*). Our findings indicate that PjRGF2 and PjRGF5 represent another set of genes co-opted specifically for parasitic function. Phylogenetic analyses revealed that these two peptides acquired haustorial functionality through distinct evolutionary trajectories: PjRGF2 aligns with other Orobanchaceae homologues, while PjRGF5 clusters with non-homologous RGF peptides (Fig. 3B). This implies that PjRGF5 is likely a more recent innovation in *P. japonicum*, arising through a series of tandem duplications and neofunctionalization (Fig. 3A). This may have begun with a duplication of PjRGF8, given the similarity of the PjRGF8 and PjRGF5 prepropeptides and the presence of LTR TEs around *PjRGF8* and *PjRGF5* copies (Fig. 3A,B). Tandem duplication followed by neofunctionalization is not unusual in RGF peptides and is seen in Arabidopsis with RGF7, GLV8, and an uncharacterized RGF peptide AT3g02245 (*46*). PjRGF2, in contrast, is preserved as a single-copy gene in both parasitic and non-parasitic Orobanchaceae species (Fig. 3B,C), suggesting that its parasitism function may have emerged via functional divergence soon after parasitism originated in the family. Investigating the regulatory mechanisms that control *PjRGF2* expression in non-parasitic and parasitic lineages will further unravel the evolution of plant parasitism.

Haustorium development is integral to plant parasitism and RGF signaling is now shown to be essential for prehaustorium formation in *P. japonicum*. Specifically, knockouts of *PjRGFR1* and *PjRGFR3* significantly reduced DMBQ-induced prehaustorium formation (Fig. 5B). In Arabidopsis, two RGFRs similarly regulate lateral root elongation (*20*), reflecting the fact that two RGFRs also govern prehaustorium initiation in *P. japonicum*. Based on our PjRGFR-BRI1 chimera assay, PjRGFR1 is solely activated by PjRGF2, whereas PjRGFR3 is activated by both PjRGF2 and PjRGF5 (Fig 4B,C), implying that PjRGF2 and PjRGF5 act redundantly. As such, a double mutant for *PjRGF2* and *PjRGF5* may be necessary to observe the phenotype seen in *pjrgfr3* and *pjrgfr1* knock-outs. Although ectodomain-BRI1 chimera assays were originally used validate known ligand-receptor pairs (*36*), our results show that they can help pinpoint receptor candidates for endogenous signaling peptides. Thus, although the precise role of PjRGFR4 during haustorium development remains unclear, our results suggest that PjRGFR4 likely responds to both PjRGF2 and PjRGF5 *in planta* (Fig 4B,C). Notably, *PjRGFR4pro* and *PjRGF2pro* expression localize to the base of the haustorium at 1 DPI (Fig. 2B, Fig. 4A), hinting that PjRGFR4 may modulate haustorium development after the initial prehaustorium stage. Therefore, further investigation into the *P. japonicum* RGF signaling pathway could provide enlightenment on haustorium development beyond the initial prehaustorium phase. Collectively, our work establishes endogenous RGF peptides and their cognate receptors as critical components of the haustorium developmental signaling pathway in parasitic plants, paving the way for future studies on how these pathways have evolved and diversified.

## Methods

### Plant material and growth conditions

*P. japonicum* seeds were surface-sterilized with 50% bleach, sown on ½ MS + 1% sucrose, and stratified at 4°C for one day in the dark. Plates were placed either flat or upright in the growth chamber at 25°C in the dark. After three days, plants were moved to 16-h day / 8-h night cycle (long day (LD) condition). Six-day-old *P. japonicum* seedlings were used for hairy root transformation. Seedlings not used for hairy root transformation were transferred to 0.7% water agar + 4.5 mM MES pH 6.4 after eight days of growth. Eleven-day-old seedlings were then transferred to 2 mL water + 4.5 mM MES pH 6.4 for HIF assays. *A. thaliana* Col-0 seeds were surface-sterilized as previously described, sown on ½ MS + 1% sucrose (*43*), stratified at 4°C for two days, and grown under 22°C LD conditions. Plates were placed upright in the growth chamber for parasitism assays. *N. benthamiana* seeds were surface-sterilized with 50% bleach, sown on ½ MS + 1% sucrose, and placed at 25°C LD conditions. Seven-day-old *N. benthamiana* seedlings were transferred to 2 mL of water + 4.5 mM MES pH 6.4. *S. hermonthica* seeds were sterilized as previously described and kept in water at 25°C (*8*). After 1 to 2 weeks, seeds were treated with 10 nM strigol to induce germination.

### *P. japonicum* hairy root transformation

Hairy root transformation was performed as previously described with slight modifications (*47*). Briefly, *Agrobacterium rhizogenes* AR1193 strains were streaked from glycerol stocks on LB supplemented with the appropriate antibiotics four days prior to transformation and grown at 28°C. Two days before transformation, streaked *A. rhizogenes* AR1193 was used to seed a 3 mL LB liquid culture that was then grown overnight at 28°C. After 10-14 hours of growth, 1 mL of the *A. rhizogenes* LB culture was added to 5 mL of AB:MES salts and grown overnight at 28°C. The next day the bacteria was pelleted and resuspended in 40 mL of 10 mM MgCl_2_ + 10 mM MES pH 5.5 + 200 µM acetosyringone + 0.01% Silwet. Six-day old *P. japonicum* seedlings were immersed in this solution as previously described and then transferred to B5 + 1% sucrose + 10 mM MES pH 5.5 + 450 µM acetosyringone agar plates and kept in the dark for two days at 22°C. Seedlings were then transferred to B5 + 1% sucrose + 300 mg/L cefotaxime and kept at 25°C LD conditions. Hairy roots typically emerged in 3 to 4 weeks. Plants with hairy roots were transferred to 0.7% water agar + 0.01% glucose + 100 mg/L Timentin (GoldBio) and were grown for at least four days at 25°C LD conditions before further use.

### RT-qPCR and RT-PCR experiments

Twelve-day-old *P. japonicum* that had been equilibrated in water + 4.5 mM MES for one day were treated with DMSO, 10 µM DMBQ, or *N. benthamiana* exudate for six hours. Root tips were collected and flash frozen in liquid nitrogen. Total RNA was extracted using the Promega max100 RNA isolation machine (Promega). First-strand cDNA was synthesized using ReverTra Ace qPCR RT kit (TOYOBO). RT-qPCR was performed using Thunderbird SYBR qPCR Mix (TOYOBO) on a Stratagene Mx3000p (Agilent). *PjUBC* was used as the internal control (*13*). Primers used in this study can be found in Supplemental Table S6.

For amplifying and sequencing *PjRGF* CDSs, total RNA was separately extracted from *P. japonicum* roots and *P. japonicum* leaves as described above. First-strand cDNA was synthesized as described above. The coding domain sequence (CDS) for *PjRGFs* was amplified from cDNA using KOD ONE (TOYOBO) and run on a 2% agarose gel preloaded with GelGreen (biotium). Bands of the correct size were gel extracted with FastGene Gel/PCR Purification kit (Nippon Genetics) and sent for Sanger sequencing (Eurofins). *PjRGF5* CDSs were subcloned into pENTR/SD/D-TOPO (ThermoFisher) and sent for Sanger sequencing.

### Plasmid construction

Promoters (roughly 2.0-2.5 kbp) used for this study were cloned from *P. japonicum* gDNA using KOD ONE, except for *PjRGF5-2b* and *PjRGF5-5*, which were synthesized (Genscript). All promoters were inserted into a binary vector upstream of a 3x-Venus-NLS and Arabidopsis HSP terminator via Infusion (Takara) or HiFi (NEB) assembly. *PjPLT* was cloned from *P. japonicum* gDNA using KOD ONE and *mNeonGreen* (*mNG*) was cloned from pICSL50015 (*48*) using KOD ONE. The *PjPLT* promoter, *mNG*, and *PjPLT* were inserted upstream of an HSP terminator via InFusion assembly. The ecto domain-coding regions for each RGFR were cloned from *P. japonicum* gDNA using KOD ONE and inserted into an ePiGreen vector already containing the coding sequences for *EFR* signal peptide and *BRI1* transmembrane and kinase domains via InFusion (*36*). RUBY was synthesized (Genscript) and then assembled into pGWB4 that was precut with Bsp1407I (ThermoFisher) using InFusion. RUBY was amplified from this vector and transferred to pAGM1310 to make it compatible with golden gate cloning (*49*). zCas9i was originally acquired from pAGM47523 and the bipartite nuclear localization signals were cloned from pFASTRK (*50, 51*). Primers, plasmids, and strains used in this study can be found in Supplementary Tables S6 and S7.

### *Agrobacterium* transformation

*A. rhizogenes* AR1193 and *A. tumafaciens* AGL1 were transformed via electroporation. Following electroporation, cells were recovered in LB medium at 28°C for two hours before being plated in LB agar that contained the appropriate antibiotics and kept at 28°C. Transformed colonies were picked two days after electroporation.

### *N. benthamiana* transient expression and PjRGFR-BRI1 band shift assay

*A. tumafaciens* AGL1 strains carrying binary vectors for constitutive expression of *PjRGFR-BRI1-3xHA, EFR-BRI1-3xHA*, or *BES1-3xHA* were grown overnight in LB media supplemented with the appropriate antibiotics for selection. The following day, the bacteria was pelleted and resuspended in infiltration buffer (10 mM MgCl_2_, 10 mM MES pH 5.5, 200 µM acetosyringone). The bacterial concentration was then adjusted to an OD600 of 0.5 for the LRR-BRI1-3xHA constructs and 0.25 for the BES1-3xHA constructs before being co-infiltrated into *N. benthamiana* leaves. 100 nM of PjRGF2 or PjRGF5 were infiltrated three days post *A. tumafaciens* infiltration. Three four mm leaf discs were collected at 0 and 30 minutes after PjRGF2/5 infiltration. Western blots were performed as described in Ngou et al. (*36*).

### Confocal microscopy

*P. japonicum* hairy roots were placed on glass bottom dishes, covered with 0.7% water agar, and allowed to acclimate for one day at 25°C LD conditions. Transformed hairy roots were then observed on a Leica SP8 WLL FALCON confocal microscope with a white light laser (WLL). Roots were treated with 10 µM DMBQ, 10 nM PjRGF2, or placed next to a host root and observed one day and two days after treatment. Additionally, *P. japonicum* roots fixed with 4% paraformaldehyde and cleared for four days in ClearSeeAlpha were stained with calcofluor white and basic fuschin (*52, 53*). These roots were mounted on SlowFade Diamond antifade mountant (ThermoFisher) and observed on a Leica SP8 WLL FALCON confocal microscope using 405 nm diode laser and WLL illumination.

### Nitro blue tetrazolium staining

*P. japonicum* seedlings were treated with either 10 µM DMBQ or DMSO. Seedlings were then taken at one- or two-days post treatment and stained with a 0.1% nitro blue tetrazolium salt (NBT; Fujifilm Wako) solution for 30-40 minutes in the dark. Roots that had a prehaustorium were excised and mounted on a slide. NBT staining was observed on an Olympus BX52 microscope (Olympus).

### *P. japonicum* genome sequencing

Genomic DNA was extracted using hexadecyltrimethylammonium bromide (CTAB) method (*30*). Healthy leaves from about one-month-old *P. japonicum* (ecotype Okayama) were flash frozen under liquid nitrogen, ground, and added into 3× volume of 2×CTAB buffer [consisting of 2% CTAB, 100 mM Tris-HCl (pH 8.0), 20 mM ethylenediaminetetraacetic acid (EDTA), 1.4 M NaCl, 2% polyvinylpyrrolidone (PVP) and 0.1% β-mercaptoethanol] on a stirring hot plate. The solution was transferred to 50 mL falcon tube and shaken at 60°C for 40 minutes before adding 1× volume of chloroform and gently mixed in a rotator at room temperature for 10 min. After centrifuging at 3500 rpm for 20 min at room temperature, aqueous phase was transferred to a new tube, 0.1× volume of 10% CTAB (consisting of 10% CTAB and 0.7 M NaCl) and 1× volume of chloroform was added. The solution was rotated gently for 10 min before being centrifuged at 3,500 rpm for 20 min. The aqueous phase was transferred to a new tube, 1× volume of isopropanol was then added and mixed gently. The solution was centrifuged at 10,000 rpm for 30 min, supernatant was discarded, and the pellet was washed with 70% ethanol. Centrifugation at 10000 rpm was repeated for 10 min. Supernatant was discarded, pellet was dissolved in 500 µL TE solution and followed by QIAGEN Genomic-tip 100/G following manufacturer’s instructions. Long read sequencing libraries were prepared using the SMRTbell Express Template Prep Kit (PacBio). Sequencing was performed on a PacBio Sequel. Long read sequences (x 48 coverage) were assembled using canu (ver1.8) with default setting (*54*). Genes were annotated using Exonerate (ver2.2) and transposable elements were predicted using HiTE (ver3.2.0) (*55, 56*).

### Alignments and phylogeny

Sequences for studied peptides and proteins were aligned using MUSCLE (ver3.8.1551) (*57*). Alignments were rendered with ESPRIT 3.0 (*58*). The maximum-likelihood phylogenetic tree for RGF prepropeptides was constructed using MEGA11 with a bootstrap value of 100, while for AP2 transcription factors phylogenetic tree Ugene (ver48.1) was used with a bootstrap value of 100 (*59, 60*). RGFR sequences from 355 species were taken from Ngou et al (*61*). The RGFRs from five additional Orobanchaceae species were added to this group to make a group of RGFRs from 360 species. Sequences were aligned and phylogenetic trees were constructed as described by Ngou et al (*61*). EFR and its homologues were used as an outgroup.

### Identification of unannotated RGF peptides in *P. japonicum* genome

Unannotated RGF peptides in *P. japonicum* were identified through tBLASTn using a list of the mature sequence of RGF peptides from 21 species as well as reciprocal tBLASTn of the mature sequence of PjRGFs. The 5’ and 3’ ends of the CDS for the RGF prepropeptide were mapped using RNA-seq data and then the sequence for the prepropeptide was amplified from *P. japonicum* cDNA. Amplified sequences were sent for sequencing. Unannotated RGF peptides from *P. cranolopha, P. kansuensis, L. lindenbergia*, and *L. phillipensis* were identified as described above. A putative prepropeptide CDS was determined using the homologous *P. japonicum* prepropeptide as a guide.

### P. japonicum and S. hermonthica HIF assays

Twelve-day-old *P. japonicum* seedling or hairy roots, equilibrated in ddH_2_O + 4.5 mM MES pH 6.4 + 100 µM Timentin (for hairy roots) + 0.01% glucose (for hairy roots) were treated with 10 µM DMBQ or synthesized PjRGF peptides (Peptide Institute and Genscript) at varying concentrations or DMSO. Haustorium formation was scored two days after treatment. One-day-old *S. hermonthica* was treated with 10 µM DMBQ, 5 nM synthesized ShRGF peptides, DMSO, or H_2_O. Haustorium formation was scored at one and two days after treatment.

### Guide RNA design

Guide RNAs were designed using CHOPCHOP, selected for high predicted efficiency and specificity to each target gene (*62*).

### CRISPR hairy root sequencing

Hairy roots from *P. japonicum* expressing gRNAs and zCAS9i were individually frozen and then macerated. Genomic DNA (gDNA) extraction buffer (200 mM Tris-HCl pH 7.5, 250 mM NaCl, 25 mM EDTA, 0.5% (w/v) SDS) was added, followed by 99% ethanol (2:1 ratio) to precipitate the gDNA, which was then resuspended in TE pH 8.0. The target region along with a small flanking region on either side was amplified from the extracted gDNA and subcloned into pUC19. Three subcloned colonies were sequenced to verify editing events.

### Statistical analysis

Statistical analysis was performed on R (ver4.3.2).

### RNA-seq analysis

*P. japonicum* seedlings were germinated and grown in 1/2 MS liquid medium supplemented with 0.25% sucrose under long-day conditions at 25 °C. After 2 weeks 5 µM DMBQ and DMSO as a mock control were supplemented to the liquid media, then whole seedlings were collected and snap-frozen at six time points: 0 h, 10 min, 30 min, 1 h, 6 h, and 24 h. Total RNA was extracted using RNeasy Plant mini kit (Qiagen). RNA-seq libraries were prepared with the KAPA mRNA HyperPrep Kit (Roche) according to the manufacturer’s directions. NEBNext Multiplex Oligos for Illumina (NEB) was used for the adaptor ligation step. Sequencing was performed on a NextSeq500 platform (single-end, 75 bp). Reads were quality controlled using fastp (ver0.24.0) (*63*), then mapped to the *P. japonicum* genome (*30*) using STAR aligner (ver2.7.11) (*64*). Read counts per gene were obtained using featureCounts in Subread (ver2.0.8) (*65*). Differentially expressed genes (DEGs) in DMBQ treatment samples were calculated using edgeR (ver4.0.16) with glmQLFTest function (*66*), following filtering out low expression genes using filterByExpr function with the default parameters and subsequent trimmed mean of M-values normalization. DEG detection result was provided in Supplemental table S1.

## Acknowledgements

We would like to thank Ms. Yoko Nagai for her technical support. The authors would also like to thank Bruno Ngou and Yasu Kadota for their helpful conversations and suggestions in regard to the PjRGFR-BRI1 chimera assays and for Bruno’s help in compiling the RGFRs from 355 species. Computations were partially performed on the NIG supercomputer at ROIS National Institute of Genetics.

## Author contributions

M.R.F. and K.S conceptualized the study; methodology, M.F., A.G., T.W., A.L., S.M., A.S., and S.Y.; data acquisition, M.R.F, A.G., T.W., A.L., R.H., S.M., A.S., and S.Y.; formal analysis, M.F., A.G., T.W., and S.Y.; Data curation, M.F., A.G., T.W., and S.Y.; writing, M.F., A.G., T.W., S.Y., and K.S.; Supervision, K.S.; project administration, K.S.; funding acquisition, K.S., S.Y., M.F., and A.G.

