## Supplemental Material for "Neofunctionalized RGF pathways drive haustorial organogenesis in parasitic plants"

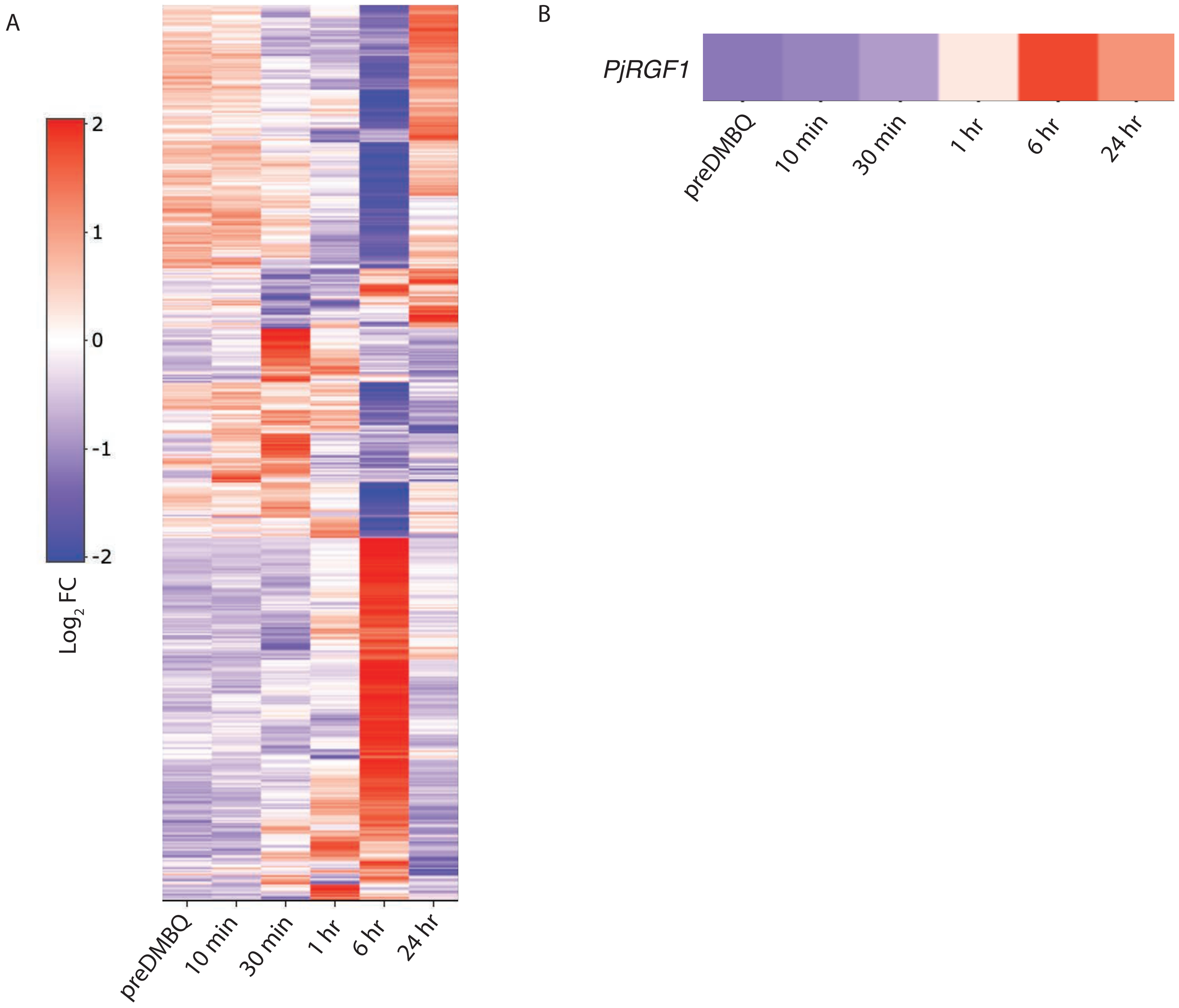


**Fig. S1. Transcriptional changes following DMBQ treatment** A) Heatmap of *P. japonicum* genes from whole *P. japonicum* seedlings that showed statistically significant changes in expression at 10 minutes, 30 minutes, one hour, 6 hours, or at 24 hours post 10 µM DMBQ treatment as compared to a preDMBQ treatment. *PjRGF1* (Pjv1_00004597) was among the genes that showed highest up-regulation at 6 hours post DMBQ treatment. B) The expression level of *PjRGF1* overtime following DMBQ treatment.

A

- **PjRGF1 - DYKPTEANPKHN**
- **PjRGF2 - DYSLPRKRRSVHN**
- **PjRGF3 - DYRGPRRHPPKNN**
- **PjRGF4 - DYMPPRTHPPVHN**
- **PjRGF5 - DYTPTTGRPPIHN**
- **PjRGF6 - DYTPARKKTPIHN**
- **PjRGF7 - DYSQARRKPPIHN**
- **PjRGF8 - DYLPPKTHPPVHN**
- **PjRGF9 - DYRMPKSHPPKNN**
- **PjRGF10 - DYRTPKSHPPKNN**
- **PjRGF11 - DYRGAARRRSPIHN**
- **PjRGF12 - DYAWLKRRHPIHN**
- **PjRGF13 - DYALPHRKPPIHN**
- **PjRGF14 - DYQPPHRKSPIHN**
- **PjRGF15 - DYAQPKPHLPIHN**
- **PjRGF16 - DYSQPKPHSPTHN**
- **PjRGF17 - DYVWIKKRHPIHN**

B

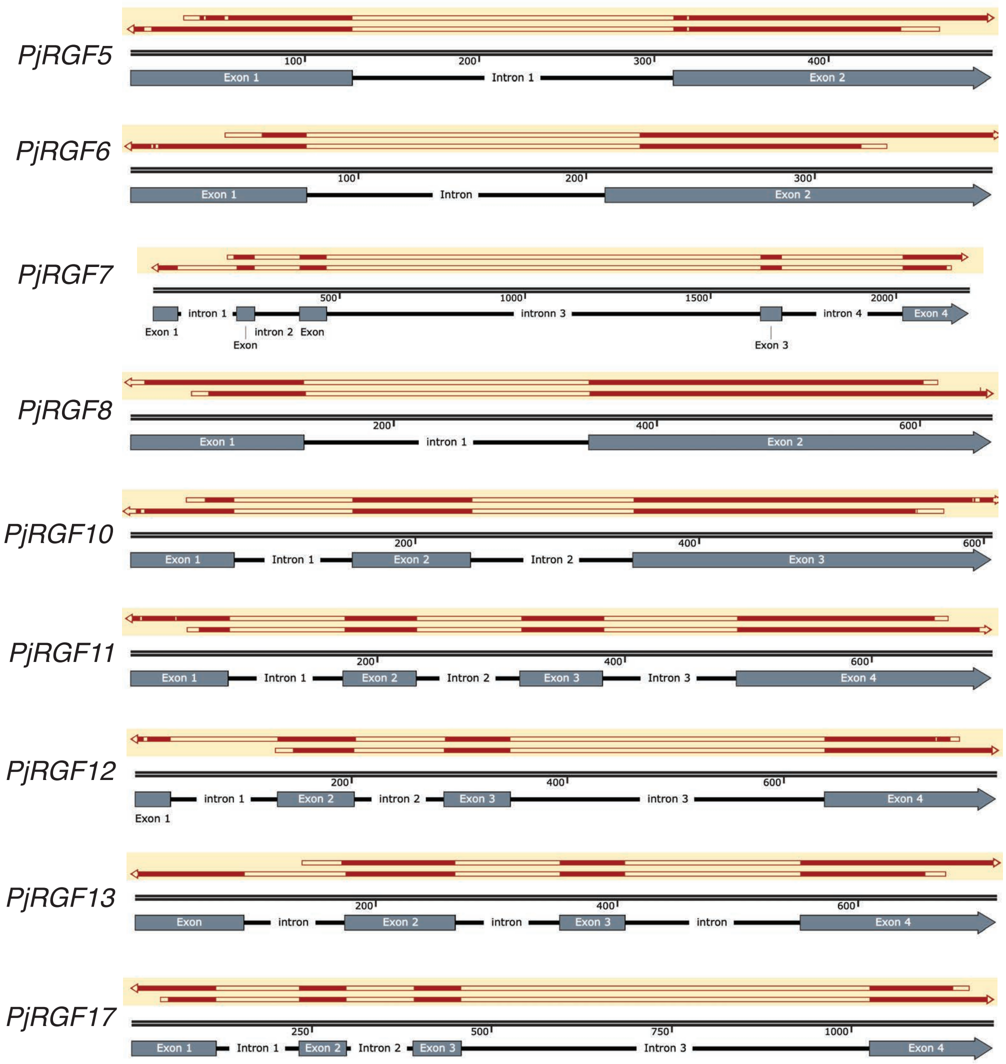


C

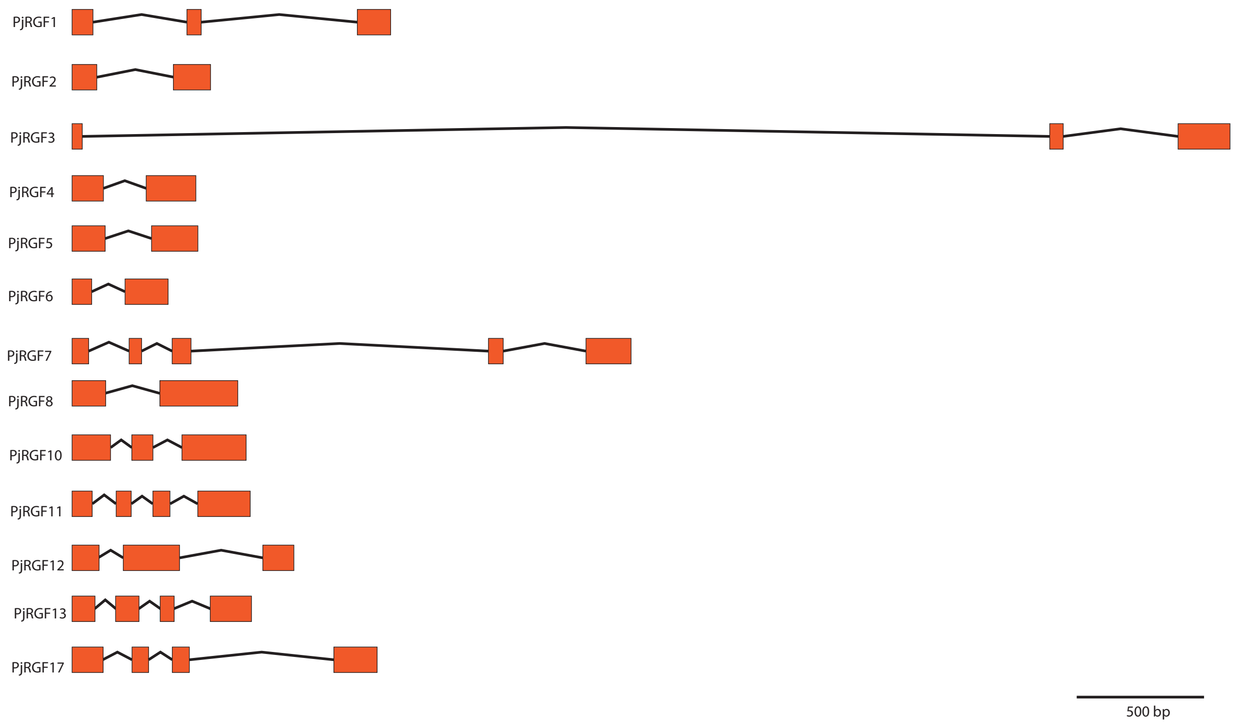


**Fig. S2. RGF peptides in *P. japonicum*** A) List of the bioactive peptides for putative RGF peptides in *P. japonicum*. Genes encoding the RGF peptides highlighted in yellow were determined to be transcribed by *P. japonicum*. B) Sanger sequencing alignments of newly identified *PjRGFs* with exons and introns of genes underneath. Length of the gene in bp is marked on the sequencet. C) Illustration of the genes encoding the 13 RGF peptides identified in *P. japonicum*.

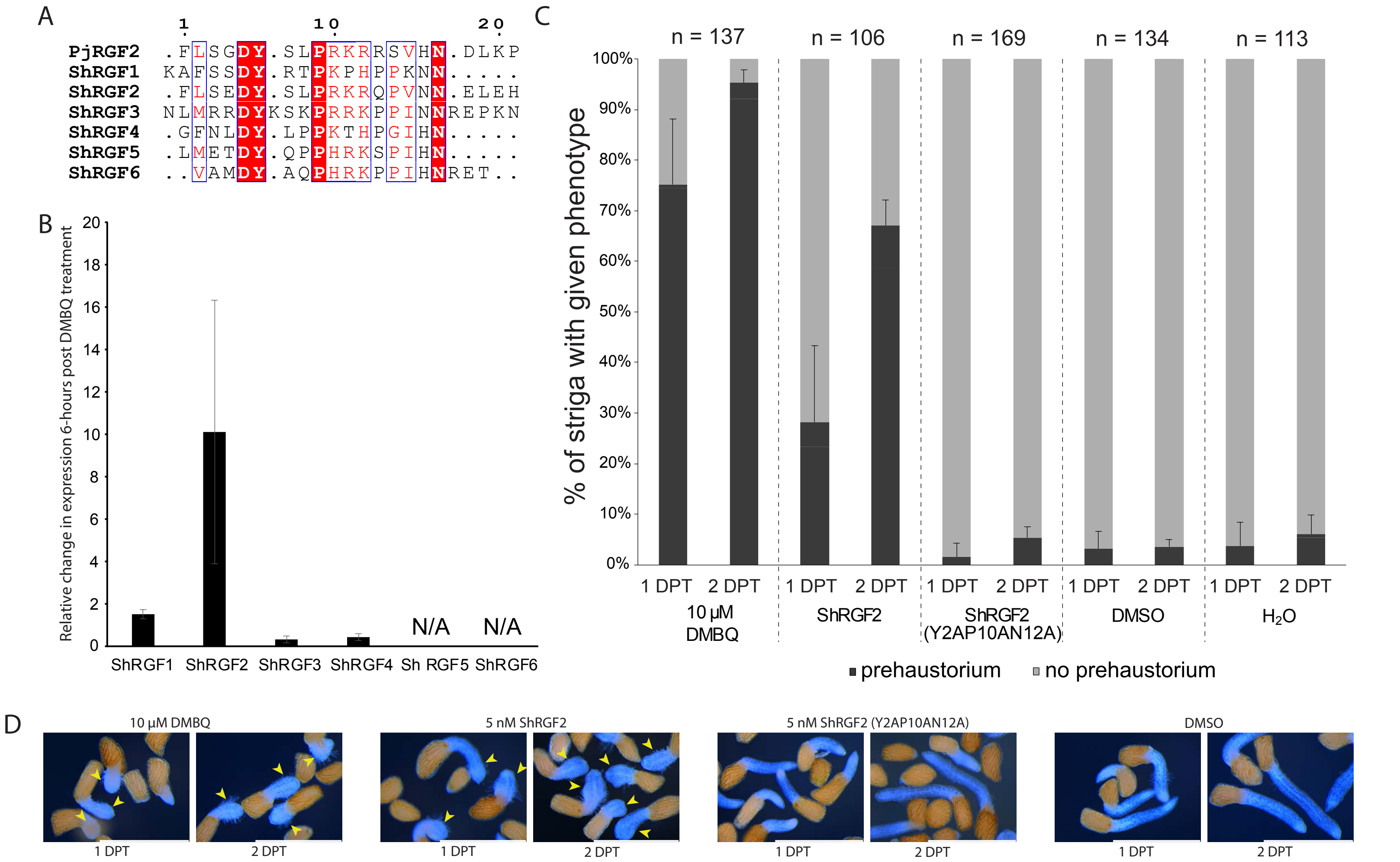


**Fig. S3.** ***S. hermonthica*** **RGF peptides induce prehaustoria** A) Multiple sequence alignment of the C-terminus of the six iannotated RGF peptides in *S. hermonthica* and PjRGF2. ShRGF2 shows high sequence similarity to PjRGF2. B) Relative change in gene expression of the genes encoding the identified ShRGF peptides six hours post 10 µM DMBQ treatment as compared to DMSO treatment. Only *ShRGF2* is highly upregulated following 10 µM DMBQ treatment. C) Percent of *S. hermonthica* that formed prehaustoria following the indicated treatments at one and two DPT. Treatment with 5 nM ShRGF2 induces prehaustorium formation in *S. hermonthica* but at a lower rate than 10 µM DMBQ. C) Photos of *S. hermonthica* seedlings at 1 DPT and 2 DPT of 10 µM DMBQ, 5 nM ShRGF2, 5 nM ShRGF2 (Y2AP10AN12A) or DMSO. Scale bars = 1 mm

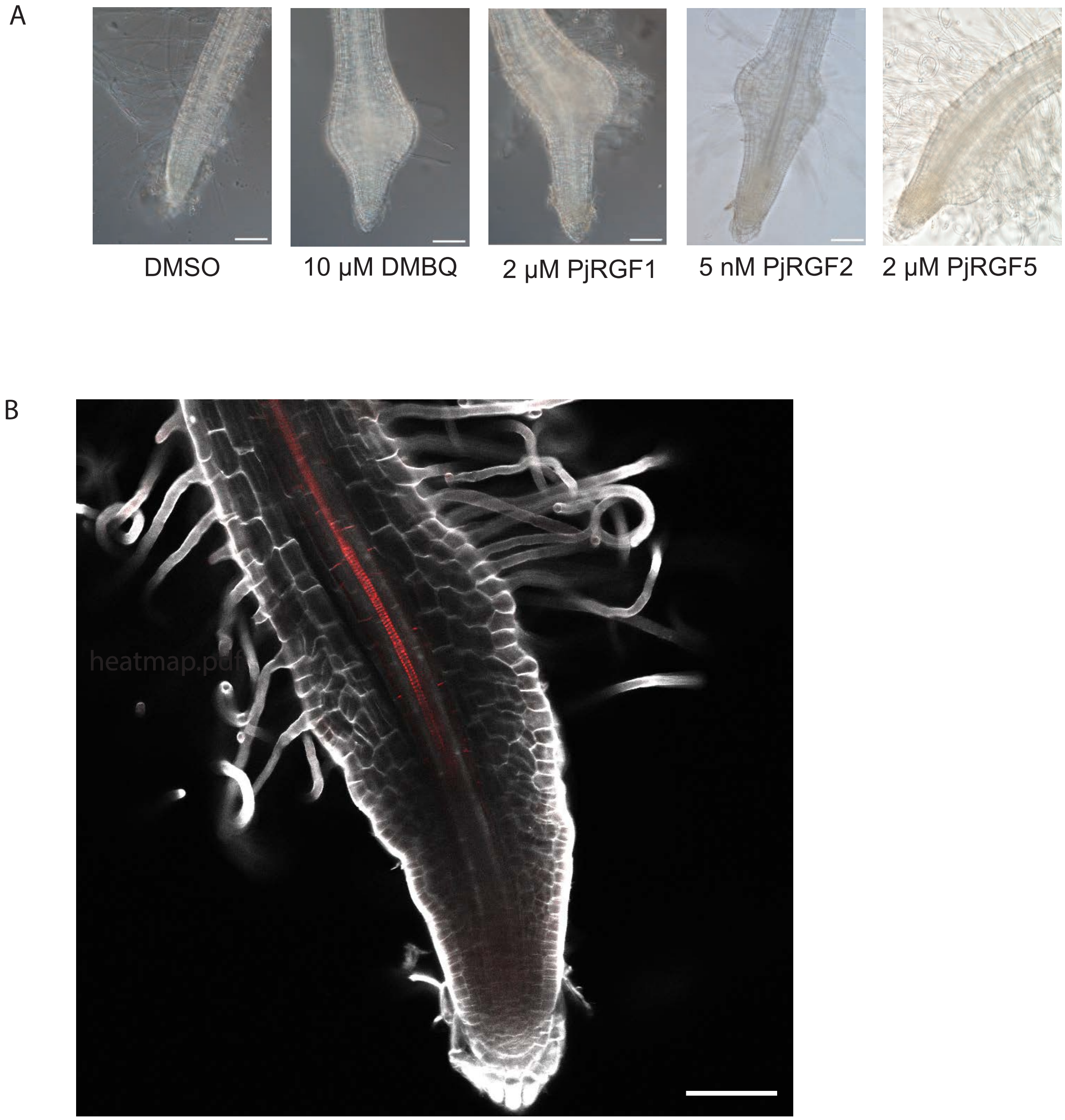


**Fig. S4. PjRGF peptide treatment phenotypes** A) Representative images of prehaustorium induced two days after treatment by 10 µM DMBQ, 2 µM PjRGF1, 5 nM PjRGF2, or 2 µM PjRGF5 and a root treated with DMSO. Scale bars equal 100 µm. B) Confocal image of a *P. japonicum* root tip two days post treatment with 2 µM PjRGF2. Abnormal expansion and division of cortex cells accompanied by anticlinal cell division of the epidermal cells results in a wavy pattern on the root. Pericycle cells do not appear to have undergone any extra anticlinal cell division. White – calcofluor white Red – basic fuschin. Scale bar is 75 µm.

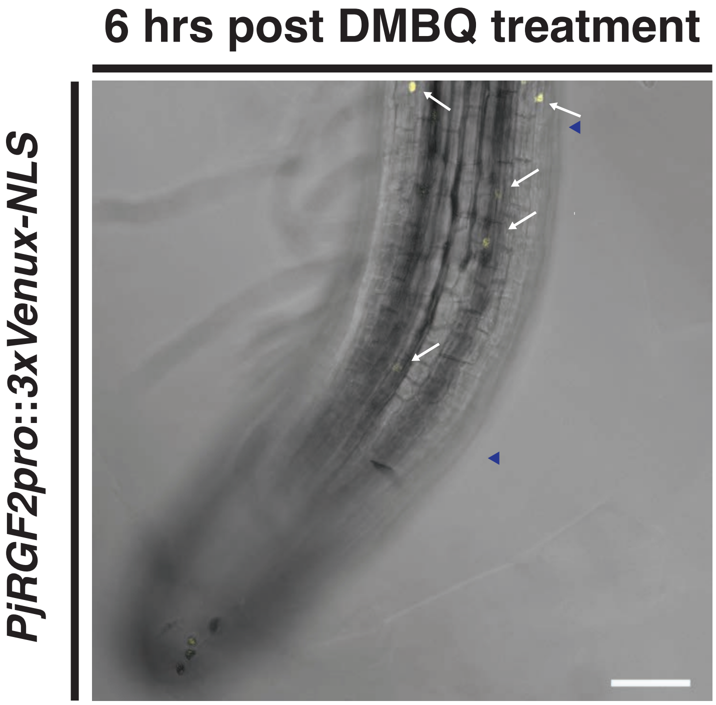


**Fig. S5. The PjRGF2 promoter is expressed at early timepoints following DMBQ treatment.** Confocal image of a *P. japonicum* hairy root expressing a *PjRGF2* transcriptional reporter (*PjRGF2pro*) at six hours post 10 µM DMBQ treatment (6 HPT). At 6 HPT expression of nuclear-localized 3x-Venus can begin to be observed in the transition zone and beginning of the elongation zone of the *P. japonicum* root. At 6 HPT *P. japonicum* roots will commonly bend and some of the root will move outside the focal plane. Scale bar is 75 µm. White arrows point to cells expressing *3x-Venus-NLS*. Blue triangles the start and end of the transition zone.

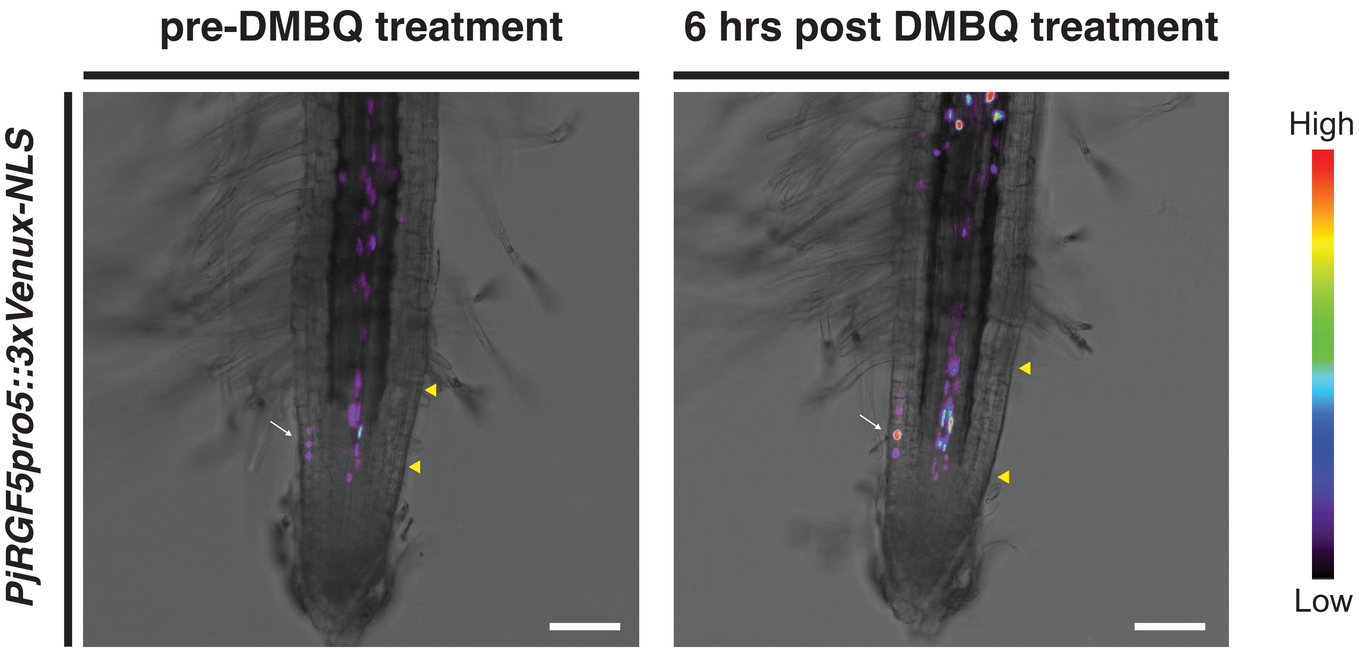


**Fig. S6. The *PjRGF5 promoter* shows an increase following DMBQ treatment.** Confocal images of *P. japonicum* hairy roots expressing a transcriptional reporter for *PjRGF5-5* (*PjRGF5-5pro*) pre-DMBQ treatment and at six hours post 10 μM DMBQ treatment (6 HPT). Spectrum artificial coloring was used to visualize the intensity of the 3x-Venus-NLS reporter. There is increased expression of the *PjRGF5-5pro* at 6 HPT can be seen to increase in the transition zone and also in the elongation zone. Scale bars are 75 μm. White arrows are pointing to epidermal expression of *PjRGF5-5pro* one the epidermis of the transition zone. Yellow triangles denote the begining and end of the transition zone.

A

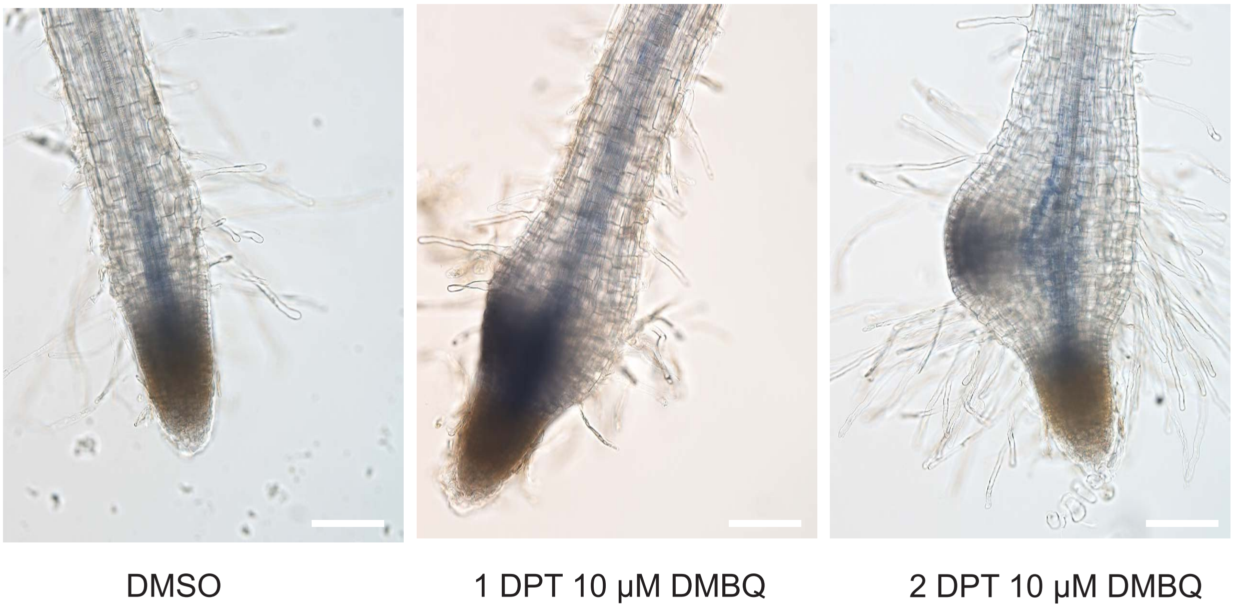


**Fig. S7. $O_2^-$ localization in *P. japonicum* roots** A) Representative images of P. japonicum roots stained with nitroblue tetrazolium (NBT) one day post treatment (1 DPT) or two days post treatment (2 DPT) with 10 μM DMBQ or following treatment with DMSO.Following DMBQ treatment, NBT stained areas within the prehaustorium. This suggests that $O_2^-$ is asymmetrically produced in the developing prehaustorium. Scale bars = 100 μm.

A

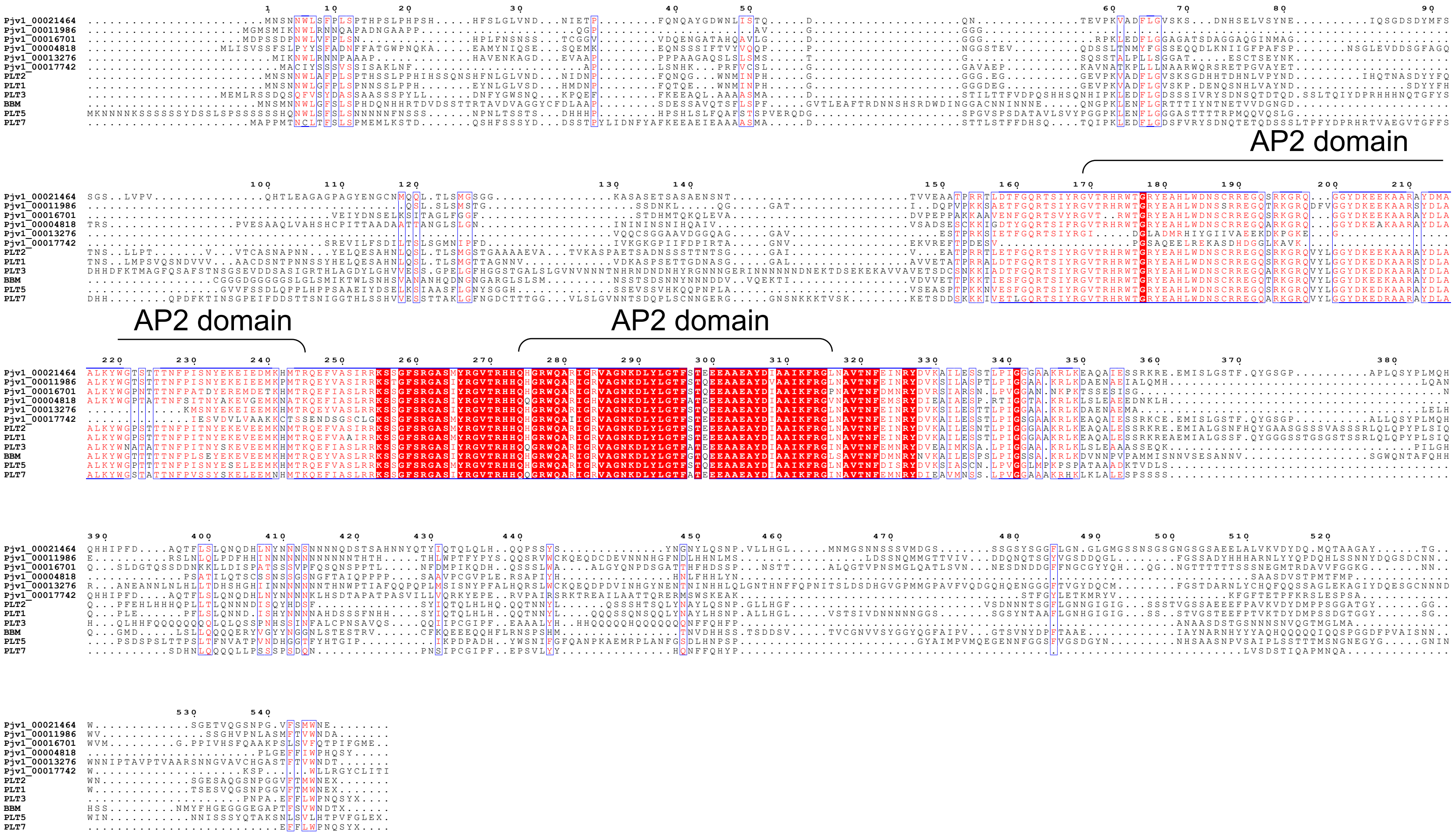


B

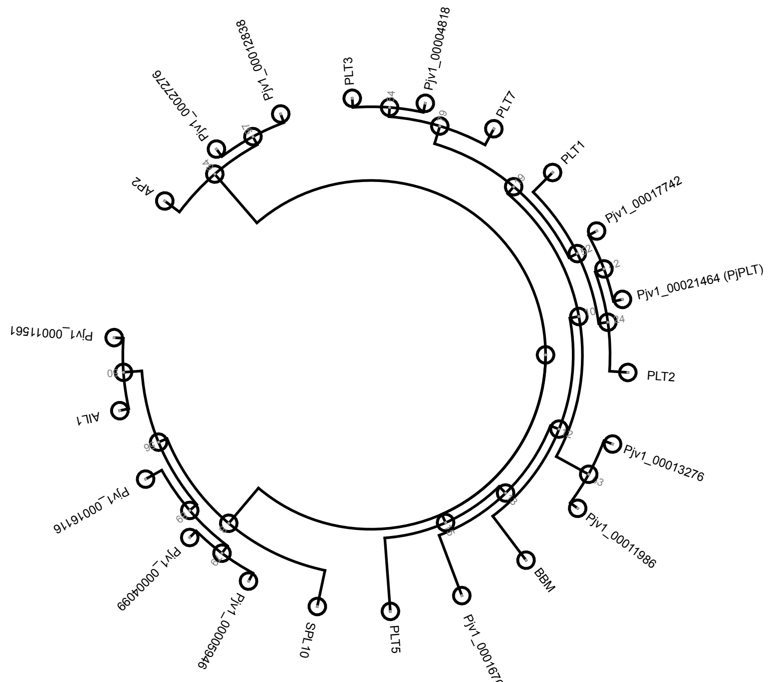


**Fig. S8. Identification of PLT homologues in *P. japonicum*** A) Multiple sequence alignment of the PLT transcription factors in Arabidopsis and homologues in *P. japonicum*. The first of two AP2 domains typically found in PLT transcription factors is missing in Pjv1_00013276 and Pjv1_0001774 B) Phylogenetic tree produced from an alignment of AP2 transcription factors in Arabidopsis and homologous genes in *P. japonicum*. Arabidopsis PLT transcription factors form a single clade within the tree and there are six *P. japonicum* homologous transcription factors within this clade. PjPLT is one of two putative homologues to PLT2 in *P. japonicum*. The other PLT2 homologue does not include two AP2 domains so it was not considered a likely homologue of PLT2. Bootstrap values are indicated at the tree nodes.

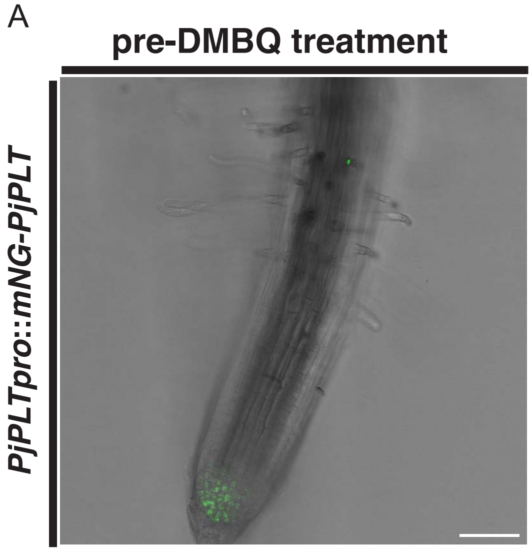


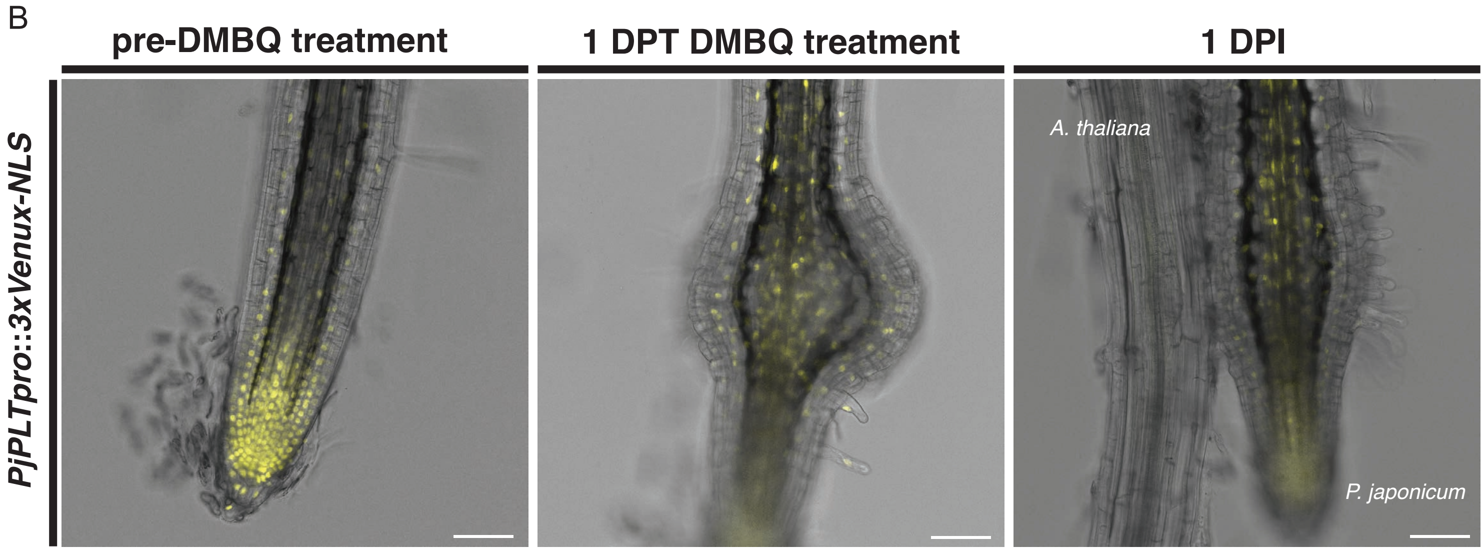


**Fig. S9. Localization and spatial expression pattern of PjPLT.** A) Confocal image of *P.japonicum* hairy root expressing mNG-PLT under its native promoter. mNG-PLT localizes to the RAM. B) Confocal images of *P. japonicum* hairy roots expressing a *PjPLT* transcriptional reporter (*PjPLTpro*). Images were taken pre-DMBQ treatment, at one day post 10 µM DMBQ treatment (1 DPT, and at 1 DPI). The *PjPLTpro* shows a gradient of expression along the untreated *P. japonicum* root. At 1 DPT and 1 DPI the expression of the *PjPLTpro* is expressed throughout the prehaustorium. Scale bars are 75 µm.

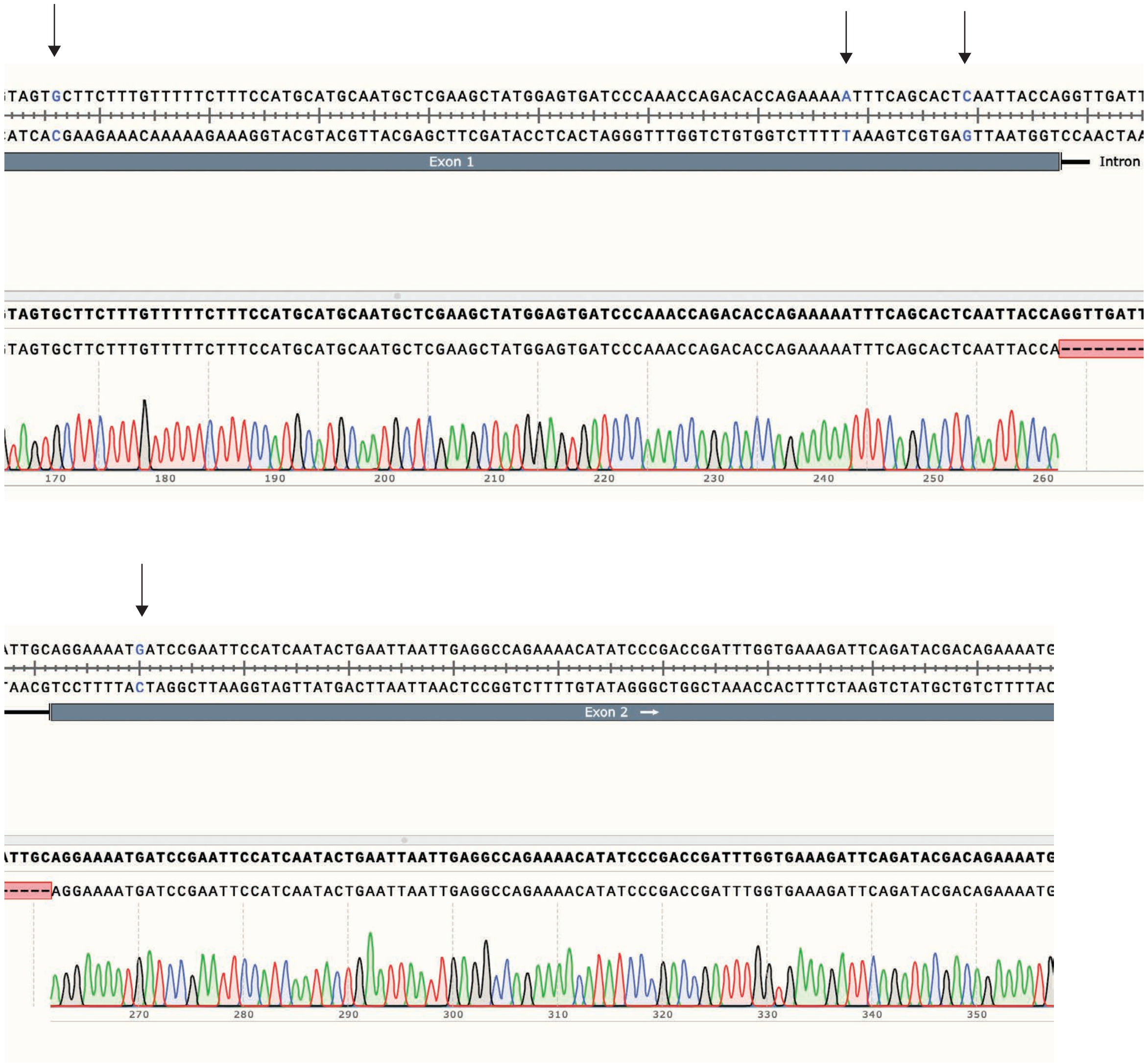


**Fig. S10. Sequencing data of unannotated *PjRGF5* variant.** Snapshots of the chromatagram from Sanger sequencing following subcloning of *PjRGF5* CDS that showed a unique, unannotated *PjRGF5* variant. The SNPs that make it unique are in blue and labelled with a black arrow.

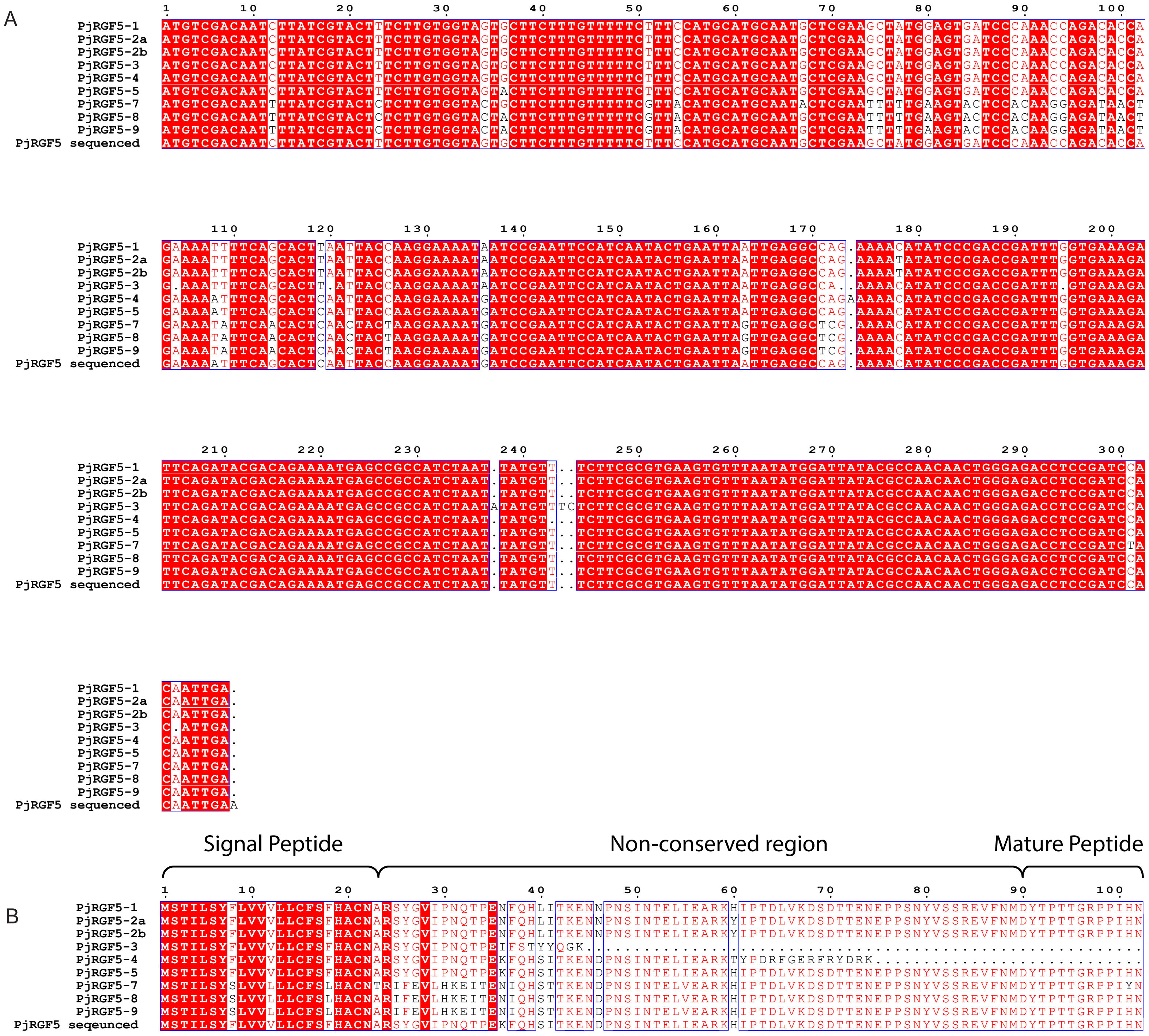


**Fig. S11. Alignments of PjRGF5 duplications** A) Multiple sequence alignment of the nine *PjRGF5* CDSs (including the ones that would be made by the pseudogenes) found in the current *P. japonicum* genome assembly and a *PjRGF5* CDS that was identified when sequencing *PjRGF5* transcript from cDNA (PjRGF5-sequenced). This *PjRGF5* sequenced CDS is not found in the current iteration of the *P. japonicum* genome. The *PjRGF5* sequences that encode viable prepropeptides only differ by SNPs at nucleotide 36, 136, or 178. B) Multiple sequence alignment of the prepropeptides encoded by the various PjRGF5 duplications in the *P. japonicum* genome. SNPs that occurred in the CDS for*PjRGF5-1/2a/2b/5* or the sequenced*PjRGF5* CDS result in changes in amino acid residues in the non-conserved residues of the prepropeptide.

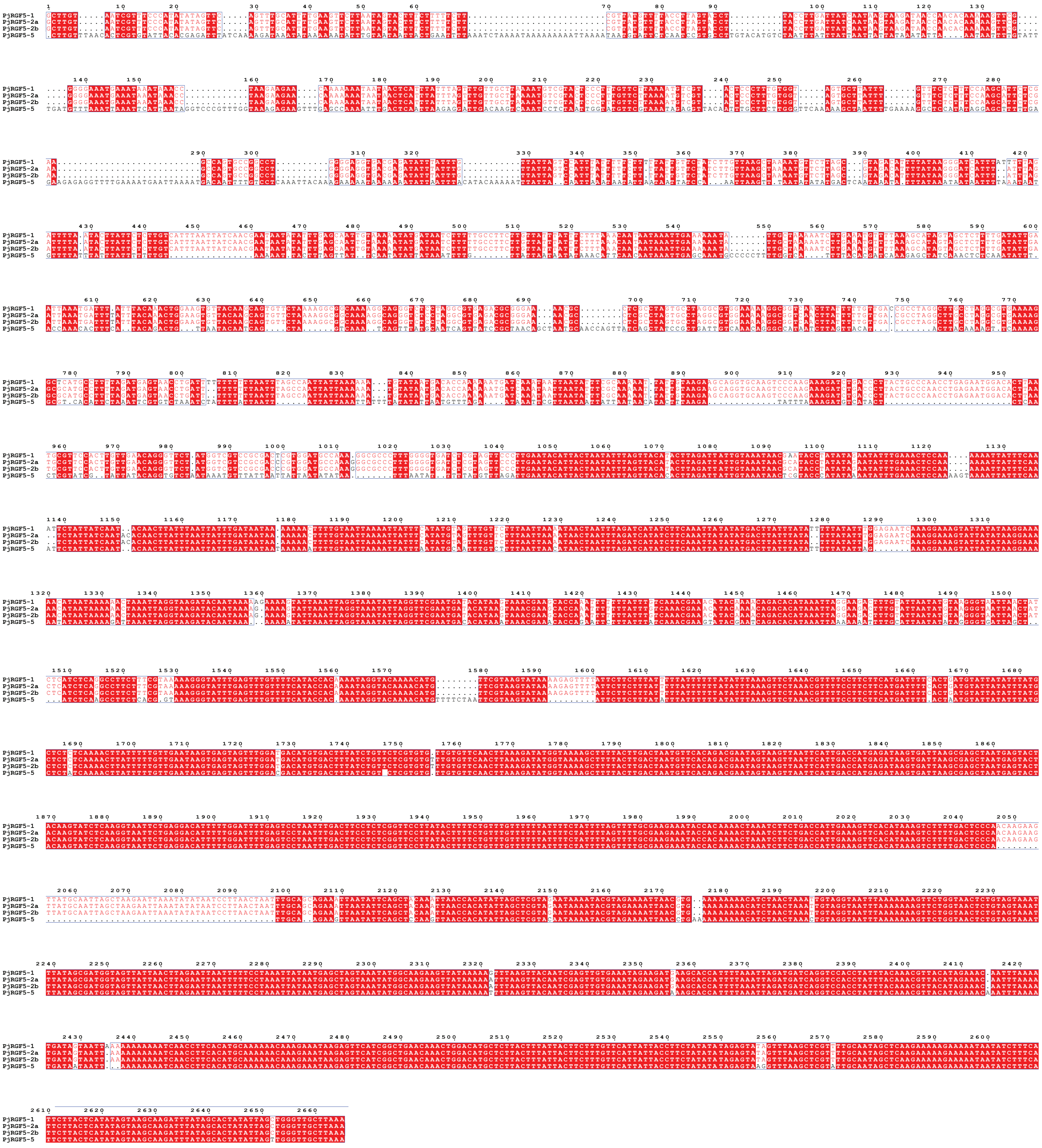

**Fig. S12. Alignment of the promoters for *PjRGF5* duplications.** Multiple sequence alignment of the sequence around 2.5 kbp upstream of the genes encoding *PjRGF5-1/2a/2b/5*. The entirety of this region shows high conserbvation for *PjRGF5-1* and *PjRGF5-2a/b* but not *PjRGF5-5*. All four sequences show greatest similartiy from nucleotide 1601-2665

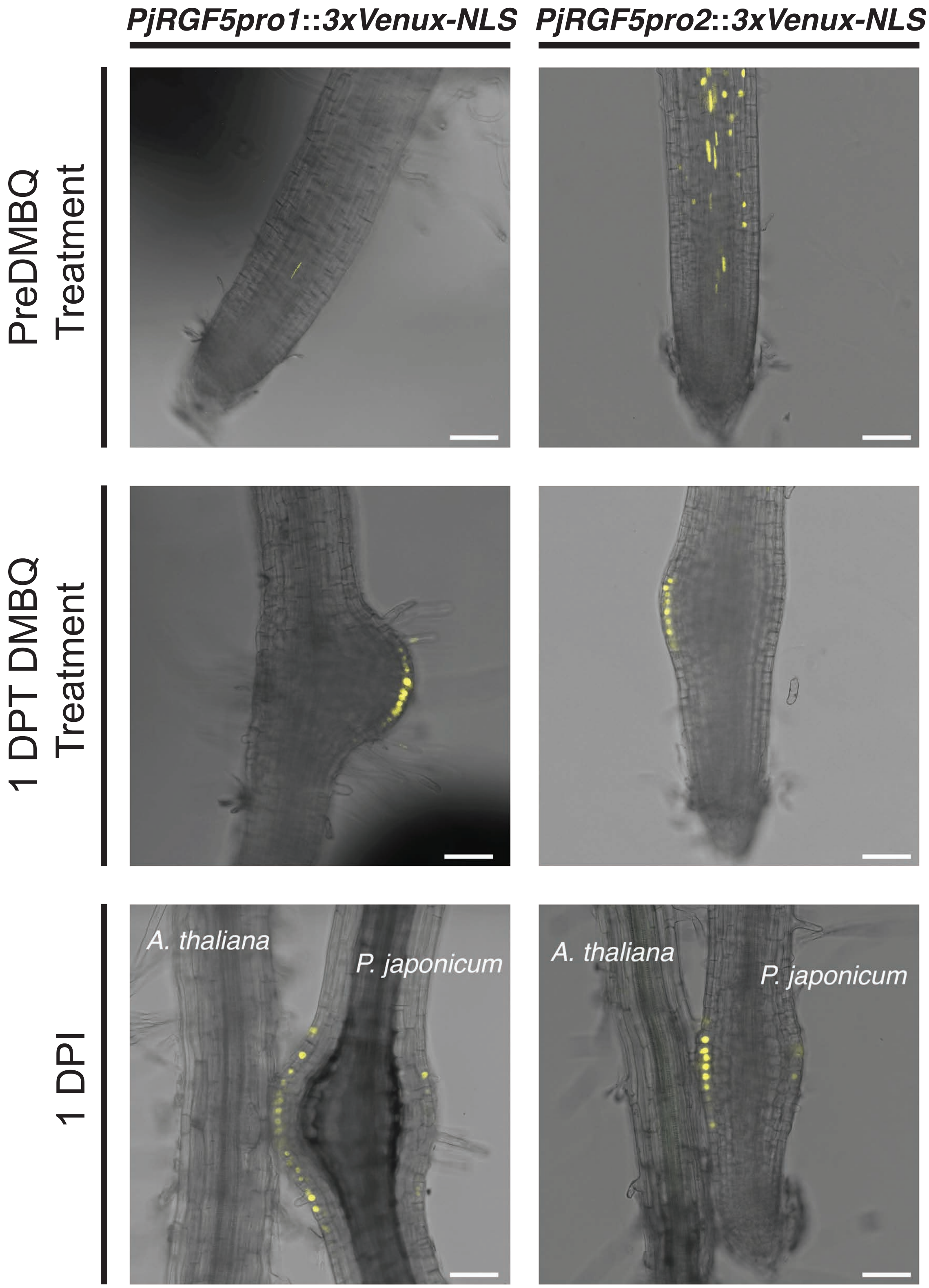


**Fig. S13. Spatial expression pattern of *PjRGF5* promoters.** Representative confocal images of *P. japonicum* hairy roots expressing transcriptional reporters for *PjRGF5-1* promoter (*PjRGF5-1pro*) and *PjRGF5-2* promoter (*PjRGF5-2pro*). Both PjRGF5 peptides are expressed in similar locations preDMBQ treatment, at 1 DPT 10 μM DMBQ treatment, and at 1 DPI. As seen with the spatial expression pattern of *PjRGF5-5pro* (Fig 2B), both *PjRGF5-1pro* and *PjRGF5-2pro* are expressed in the epidermal cells of the prehaustorium of *P. japonicum*. Scale bars are 75 μm.

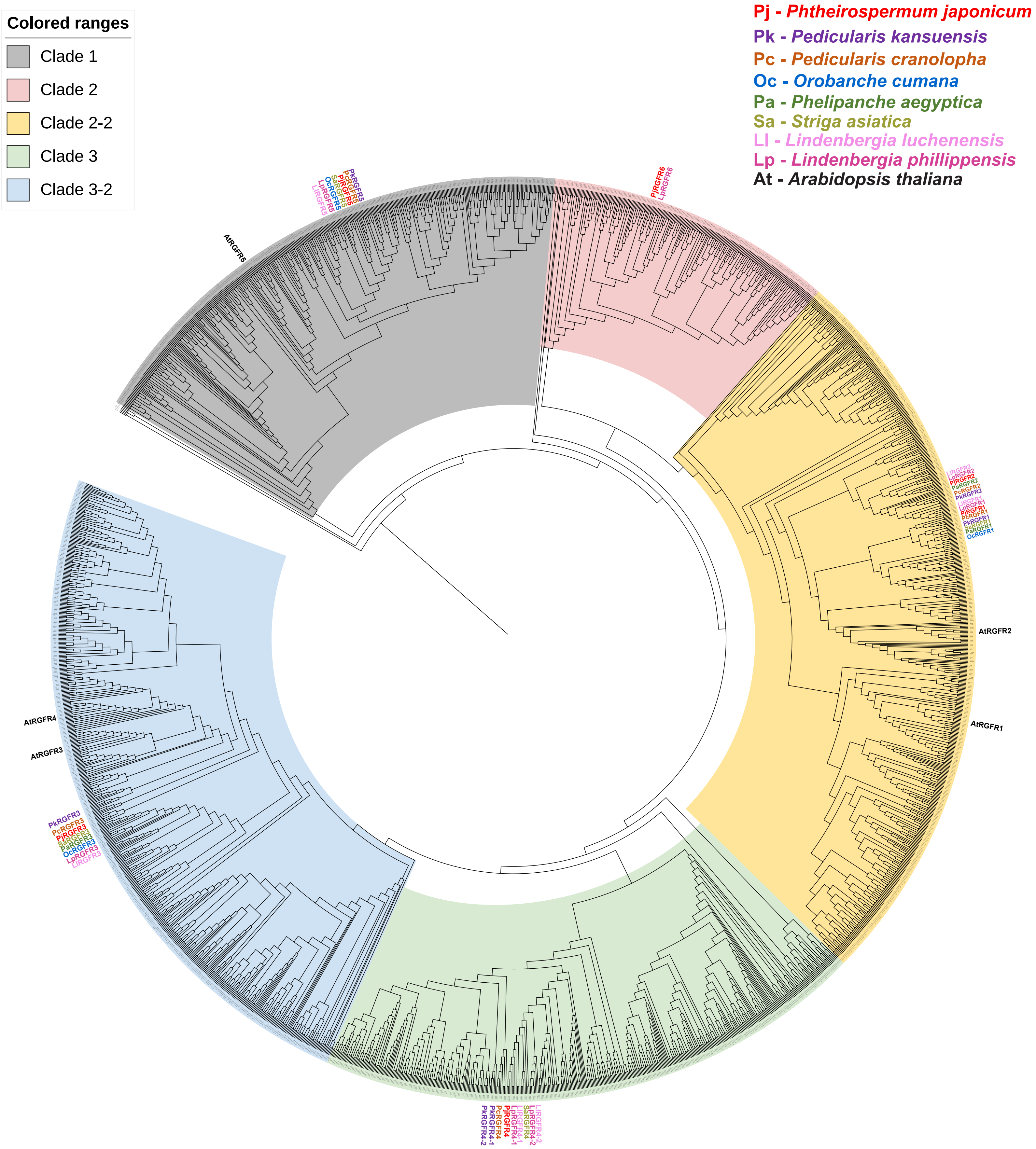


**Fig. S14. Phylogenetic tree of the full sequence of RGFRs found in 360 species.** RGFRs group into three clades, with clade 2 and clade 3 forming two subclades. RGFRs from eight species in the Orobanchaceae family, including *P. japonicum*, and the RGFRs from Arabidopsis are labelled. Phylogenetic distances between RGFR sequences have been ignored in this tree.

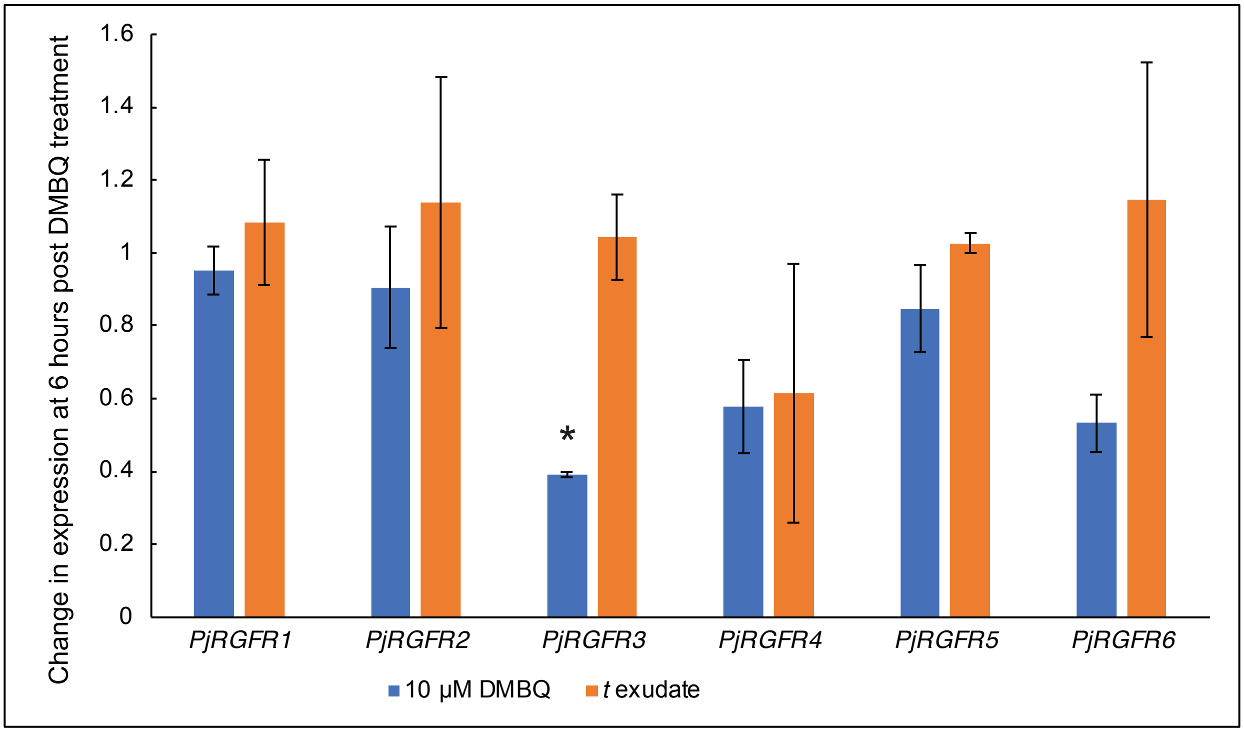


**Fig. S15. Change in expression of PjRGFRs following HIF treatment.** qRT-PCR data showing the change in expression of all the PjRGFRs encoded in the P. japonicum genome at 6 hours post DMBQ or N. benthamiana exudate treatmtment. PjUBC was used as a housekeeping gene. An * denotes changes in gene expression with a p-value < 0.05 as compared to a DMSO control.

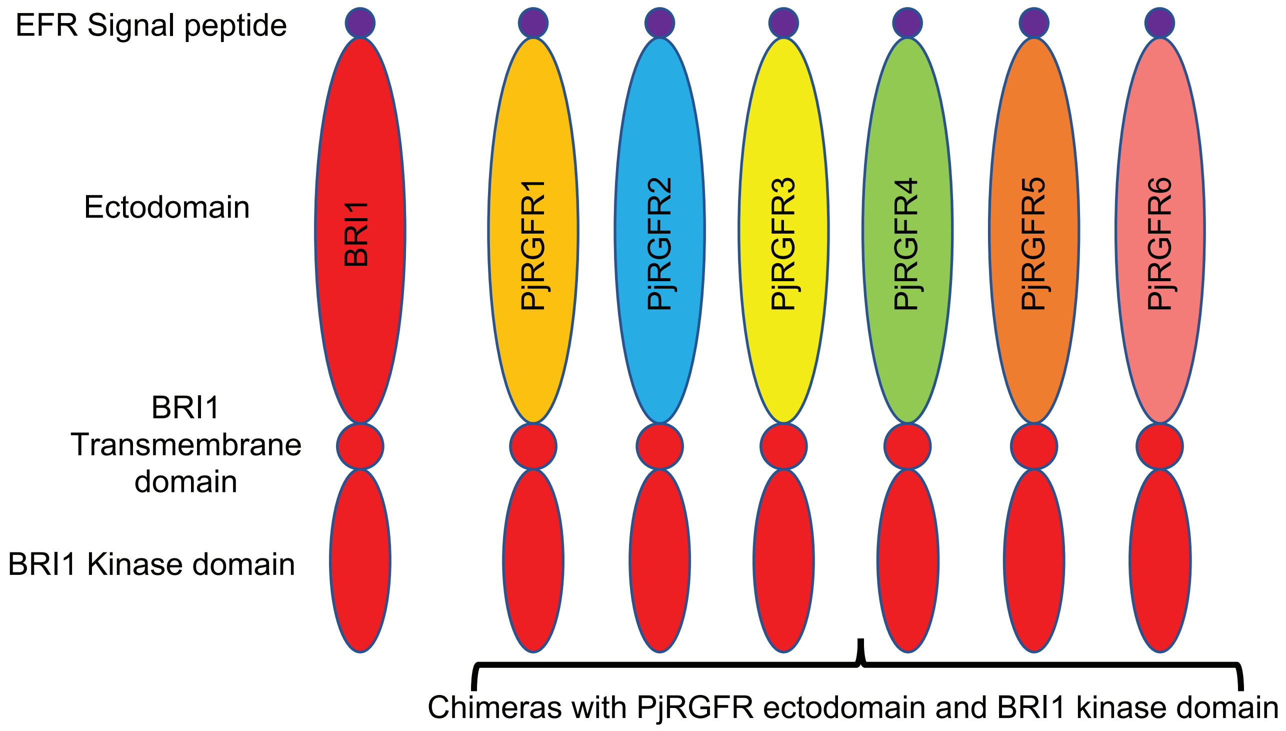


**Fig. S16. Construction of PjRGFR-BRI1 chimeras.** A) Diagram showing how the PjRGFR-BRI1 chimeras were constructed. All the chimeras consist of the designated PjRGFR ectodomain, the EFR signal peptide, BRI1 transmembrane domain, and BRI1 kinase domain

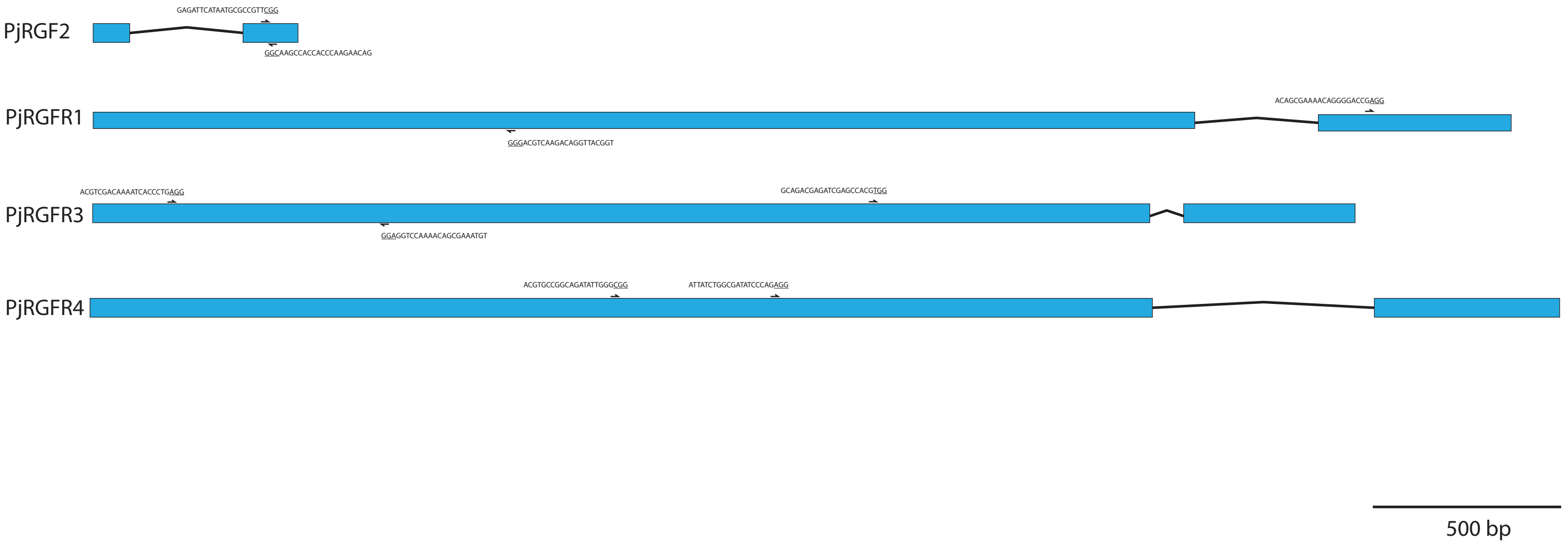


**Fig. S17. Locations of gRNAs used during hairy root CRISPR.** Illustrationns of the genes *PjRGF2*, *PjRGFR1*, *PjRGFR3*, and *PjRGFR4* with the locations of the gRNAs used during CRISPR gene editing.

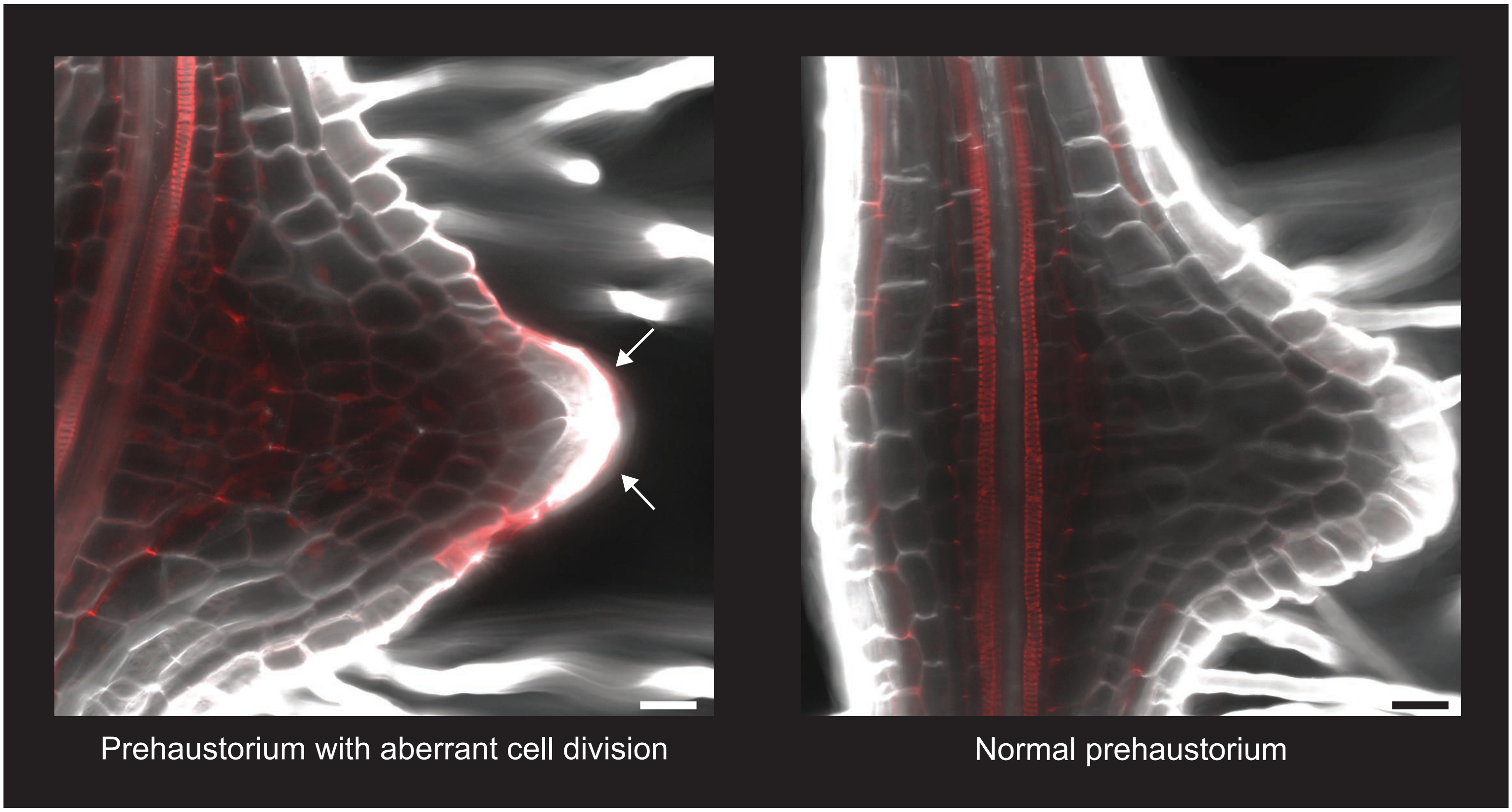


**Fig. S18. Images of *P. japonicum* prehaustoria**. A prehaustorium with aberrant cell division on the left and normal cell division on the right. The prehaustorium with aberrant cell division has a more pointed structure than the normal prehaustorium. The apex of the prehaustorium is covered by two large cells that are divided at the tip of the prehaustorium. This is not observed in a normal prehaustorium. White arrows point to large cells at the tip of the prehaustoriums. Basic fuschin - Red, Calcofluor white - white. Scale bars are 25 μm.

Supplemental Table S3. Orobanchaceae RGFR gene ID and gene names

| ID # | Gene name |
|---|---|
| *Phtheirospermum japonicum* | |
| Pjv1_00003866 | PjRGFR1 |
| Pjv1_00010527 | PjRGFR2 |
| Pjv1_00011762 | PjRGFR3 |
| Pjv1_00001310 | PjRGFR4 |
| Pjv1_00019566 | PjRGFR5 |
| Pjv1_00020079 | PjRGFR6 |
| *Pedicularis cranolopha* | |
| T326817N0C002G04530 | PcRGFR1 |
| T326817N0C000G03413 | PcRGFR2 |
| T326817N0C006G02525 | PcRGFR3 |
| T326817N0C008G00686 | PcRGFR4 |
| T326817N0C001G02733 | PcRGFR5 |
| *Pedicularis kansuensis* | |
| evm.model.CTG_208.3 | PkRGFR1 |
| evm.model.CTG_888.24 | PkRGFR2 |
| evm.model.CTG_3012.4 | PkRGFR3 |
| evm.model.CTG_218.93 | PkRGFR4-1 |
| evm.model.CTG_1877.6 | PkRGFR4-2 |
| evm.model.CTG_299.53 | PkRGFR5 |
| *Phelipanche aegyptica* | |
| GWHPBHPU009855 | PaRGFR1 |
| GWHPBHPU011410 | PaRGFR2 |
| GWHPBHPU017060 | PaRGFR3 |
| *Orobanche cumana* | |
| GWHPBHPV022007 | OcRGFR1 |
| GWHPBHPV038702 | OcRGFR3 |
| GWHPBHPV048321 | OcRGFR5 |
| *Lindenbergia luchenensis* | |
| GWHPBHPT016274 | LlRGFR1 |
| GWHPBHPT036181 | LlRGFR2 |
| GWHPBHPT017559 | LlRGFR3 |
| GWHPBHPT017039 | LlRGFR4-1 |
| GWHPBHPT010114 | LlRGFR4-2 |
| GWHPBHPT022036 | LlRGFR5 |
| *Lindenbergia phillippensis* | |
| Lphilippensis_11928 | LpRGFR1 |
| Lphilippensis_24740 | LpRGFR2 |
| Lphilippensis_13055 | LpRGFR3 |
| Lphilippensis_12391 | LpRGFR4-1 |
| Lphilippensis_20014 | LpRGFR4-2 |
| Lphilippensis_5496 | LpRGFR5 |

| | |
|---|---|
| Lphilippensis_23448 | LpRGFR6 |
| *Striga hermonthica* | |
| CAA0823899 | ShRGFR1 |
| CAA0832479 | ShRGFR3 |
| CAA0836134 | ShRGFR4 |
| CAA0830122 | ShRGFR5 |
| *Striga asiatica* | |
| Sasiatica_22940 | SaRGFR1 |
| Sasiatica_19278 | SaRGFR3 |
| Sasiatica_3184 | SaRGFR4 |
| Sasiatica_27030 | SaRGFR5 |

Table S4. Percentage of edited hairy roots per construct

| **Target** | **Knockout** | **Unedited** | **Total sequenced** | **Efficiency** |
|---|---|---|---|---|
| *PjRGF2* | 23 | 85 | 108 | 20% |
| *PjRGFR1* | 21 | 87 | 108 | 19% |
| *PjRGFR3* | 20 | 44 | 62 | 32% |
| *PjRGFR4* | 23 | 28 | 51 | 45% |

Supplemental Table S5. Edits and phenotypes for *P. japonicum* hairy roots

| **Target Gene** | | | |
|---|---|---|---|
| PjRGF2 | Sequence | Edits | Phenotype |
| WT | 5'-AAGAGAGATTCATAATGCGCCGTTCGGTGGTGGGTTCTTGTCTGGGGATTATTCACTACC-3' | N/A | |
| | 5'-AAGAGAGATTCATAATGCGCCGTTCGGTGGTGGGTTCTTGTCTGGGGATTATTCACTACC-3' | N/A | |
| #1 | 5'-AAGAGAGATTCATAATGCGCCTGTTCGGGTGGTGGGTTCTTGTCTGGGGATTATTCACTACC-3' | +2 bp | Prehaustorium |
| | 5'-AAGAGAGATTCATAATGCGCCGTTCGGGTGGTGGGTTCTTGTCTGGGGATTATTCACTACC-3' | +1 bp | |
| #2 | 5'-AAGAGAGATTCATAATGCGCCGTT-------GGGTTCTTGTCTGGGGATTATTCACTACC-3' | -7 bp | Prehaustorium |
|  | 5'-AAGAGAGATTCATAATGCGCCGTT-------GGGTTCTTGTCTGGGGATTATTCACTACC-3' | -7 bp |  |
| #3 | 5'-AAGAGAGATTCATAATGCGC-GTTCGGTGGTGGGTTCTTGTCTGGGGATTATTCACTACC-3' | -1 bp | Prehaustorium |
|  | 5'-AAGAGAGATTCATAATGCGC-GTTCGG------GTTCTTGTCTGGGGATTATTCACTACC-3' | -7 bp |  |
| #4 | 5'-AAGAGAGATTCATAATGCGC-GTTCGGTGGTGGGTTCTTGTCTGGGGATTATTCACTACC-3' | -1 bp | No prehaustorium |
| | 5'-AAGAGAGATTCATAATGCGC-GTTCGGTGGTGGGTTCTTGTCTGGGGATTATTCACTACC-3' | -1 bp | |
| #5 | 5'-AAGAGAGATTCATAATGCGC-GTTCGGTGGTGGGTTCTTGTCTGGGGATTATTCACTACC-3' | - 1 bp | No prehaustorium |
| | 5'-AAGAGAGATTCATAATGCGC-GTTCGGTGGTGGGTTCTTGTCTGGGGATTATTCACTACC-3' | -1 bp | |
| #6 | 5'-AAGAGAGATTCATAATGCGA-GTTCGGTGGTGGGTTCTTGTCTGGGGATTATTCACTACC-3' | C -> A, -1 bp | Prehaustorium |
| | 5'-AAGAGAGATTCATAATGCGC-GTTCGGTGGTGGGTTCTTGTCTGGGGATTATTCACTACC-3' | -1 bp | |
| #7 | 5'-AAGAGAGATTCATAATGCGC-GTTCGGTGGTGGGTTCTTGTCTGGGGATTATTCACTACC-3' | -1 bp | No prehaustorium |
|  | 5'-AAGAGAGATTCATAATGCGCCCGTTCGG------GTTCTTGTCTGGGGATTATTCACTACC-3' | +1 bp, -6 bp |  |
| #8 | 5'-AAGAGAGATTCATAATGCGC-GTTCGGTGGTGGGTTCTTGTCTGGGGATTATTCACTACC-3' | -1 bp | Prehaustorium |
| | 5'-AAGAGAGATTCATAATGCGC-GTTCGGTGGTGGGTTCTTGTCTGGGGATTATTCACTACC-3' | -1 bp | |
| #9 | 5'-AAGAGAGATTCATAATGGGCTCATATGAAAAGACGAT----------------------C-3' | G->C, +17 bp, -39 bp | Prehaustorium |
|  | 5'-AAGAGAGATTCATAATGCGCCGTTCGGT-------TCTTGTCTGGGGATTATTCACTACC-3' | -7 bp |  |
| #10 | 5'-AAGAGAGATTCATAATGCGCCAGTTCGGTGGTGGGTTCTTGTCTGGGGATTATTCACTACC-3' | +1 bp | No prehaustorium |
| | 5'-AAGAGAGATTCATAATGCGCCAGTTCGGTGGTGGGTTCTTGTCTGGGGATTATTCACTACC-3' | +1 bp | |
| #11 | 5'-AAGAGAGATTC-------------------------CTTGTCTGGGGATTATTCACTACC-3' | -25 bp | Some cell division |
|  | 5'-AAGAGAGATTC-------------------------CTTGTCTGGGGATTATTCACTACC-3' | -25 bp |  |
| #12 | 5'-AAGAGAGATTCATAATGCGCCGTTCG----TGGGTTCTTGTCTGGGGATTATTCACTACC-3' | -4 bp | Aborted prehaustorium |
|  | 5'-AAGAGAGATTCATAATGCGCCGTTCG----TGGGTTCTTGTCTGGGGATTATTCACTACC-3' | -4 bp |  |
| #13 | 5'-AAGAGAGATTCATAATGCGCCGTTCG-TGGTGGGTTCTTGTCTGGGGATTATTCACTACC-3' | -1 bp | No prehaustorium |
|  | 3'-AGA---------- ------------CACCACCCAAGAACAGACCCCTAATAAGTGATGG-5' | -130 bp |  |
| #14 | 5'-AAGAGAGATTCATAATGCGCCGTTCG-TGGTGGGTTCTTGTCTGGGGATTATTCACTACC-3' | -1 bp | Prehaustorium with aberrant cell divisiion |
| | 5'-AAGAGAGATTCATAATGCGCCGTTCGGGTGGTGGGTTCTTGTCTGGGGATTATTCACTACC-3' | +1 bp | |
| #15 | 5'-AAGAGAGATTCATAATGCG-------GTGGTGGGTTCTTGTCTGGGGATTATTCACTACC-3' | -7 bp | Prehaustorium with aberrant cell divisiion |
| | 5'-AAGAGAGATTCATAATGCGCCGTTCG-TGGTGGGTTCTTGTCTGGGGATTATTCACTACC-3' | -1 bp | |
| #16 | 5'-AAGAGAGATTCATAATGCGCCGTTCGATTCAAGAAAAGACGATCAGTTCACAATGACGATCAGTTCACAATGCGCCG TTTCTTGAATCGACGGCTGCAAA----TGGGTTCTTGTCTGGGGATTATTCACTACC-3 | -4 bp, +74 bp | Prehaustorium |
| | 5'-AAGAGAGATTCATAATGCGCCGTTCG-TGGTGGGTTCTTGTCTGGGGATTATTCACTACC-3' | -1 bp | |
| #17 | 5'-AAGAGAGATTCATAATGCGCCGTTCGGGTGGTGGGTTCTTGTCTGGGGATTATTCACTACC-3' | +1 bp | Prehaustorium with aberrant cell divisiion |
| | 5'-AAGAGAGATTCATAATGCGCCGTTCG-TGGTGGGTTCTTGTCTGGGGATTATTCACTACC-3' | -1 bp | |
|  | 5'-AAGAGAGATTCATAATGCGCCGTT-------------TTGTCTGGGGATTATTCACTACC-3' | -13 bp |  |
| #18 | 5'-AAGAGAGATTCATAATGCGCCGTTCG-TGGTGGGTTCTTGTCTGGGGATTATTCACTACC-3' | -1 bp | No prehaustorium |
|  | 5'-AAGAGAGATTCATAATGCGCCGTTC-----------------------------ACTACC-3' | -29 bp |  |
| | 5'-AAGAGAGATTCATAATGCGCCTGTTCGGGTGGTGGGTTCTTGTCTGGGGATTATTCACTACC-3' | +2 bp | |
| #19 | 5'-AAGAGAGATTCATAATGCGC-----------------TTGTCTGGGGATTATTCACTACC-3' | -17 bp | Preahaustorium |
|  | 5'-AAGAGAGATTCATAATGCGC-----------------TTGTCTGGGGATTATTCACTACC-3' | -17 bp |  |
| #20 | 5'-AAGAGAGATTCATAATGCGCCGTTCG-TGGTGGGTTCTTGTCTGGGGATTATTCACTACC-3' | -1 bp | No prehaustorium |
| | 5'-AAGAGAGATTCATAATGCGCCGTTCG-TGGTGGGTTCTTGTCTGGGGATTATTCACTACC-3' | -1 bp | |
| #21 | 5'-AAGAGAGATTCATAATGCGCCTGTTCGGTGGTGGGTTCTTGTCTGGGGATTATTCACTACC-3' | +1 bp | Prehaustorium |
| | 5'-AAGAGAGATTCATAATGCGCCGTTCG-TGGTGGGTTCTTGTCTGGGGATTATTCACTACC-3' | -1 bp | |
| #22 | 5'-AAGAGAGATTCATAATGCGC-GTTCGGTGGTGGGTTCTTGTCTGGGGATTATTCACTACC-3'(-1 bp) | -1 bp | No prehaustorium |
|  | 5'-AAGAGAG--------------------------GTTCTTGTCTGGGGATTATTCACTACC-3' | -26 bp |  |
| #23 | 5'-AAGAGAGATTCATAATGCGCCGTTCG-TGGTGGGTTCTTGTCTGGGGATTATTCACTACC-3' | -1 bp | Prehaustorium with aberrant cell divisiion |
| | 5'-AAGAGAGATTCATAATGCGCCAGTTCGGTGGTGGGTTCTTGTCTGGGGATTATTCACTACC-3' | +1 bp | |
| PjRGFR3 | | | |
| WT | 5'- CGCCACGTCGACAAAATCACCCTGAGGT…TAACTCCGGCAGACGAGATCGAGCCACGTGGCGGCCGCGCGAAGG-3' | N/A | |
| | 5'- CGCCACGTCGACAAAATCACCCTGAGGT…TAACTCCGGCAGACGAGATCGAGCCACGTGGCGGCCGCGCGAAGG-3' | N/A | |
| #1 | 5'- CGCCACGTCGACAAAATCA-CCTGAGGT…TAACT-----------------------------CCGCGCGAAGG-3 | -29 bp, -1 bp | Some cell division |
|  | 5'- CGCCACGTCGACAAAATCA-CCTGAGGT…TAACTCCGGCAGACGAGATCGA--------------GCGCGAAGG-3' | -14 bp, -1 bp |  |
|  | 5'- -------------------CCCTGAGGT…TAACT-----------------------------CCGCGCGAAGG-3' | -29 bp, +1 bp, -533 bp, -1 bp |  |
| #2 | 5'- CGCCACGTCGACAAAATCA-CCTGAGGT…TAACTCCGGCAGACGAGATCGAGCCACGTGGCGGCCGCGCGAAGG-3' | -1 bp | prehaustorium |
| | 5'- CGCCACGTCGACAAAATCA-CCTGAGGT…TAACTCCGGCAGACGAGATCGAGCCACGTGGCGGCCGCGCGAAGG-3' | -1 bp | |
| #3 | 5'- CGCCACGTCGACAAAATCA-CCTGAGGT…TAACTCCGGCAGACGAGATCGAGCCACGTGGCGGCCGCGCGAAGG-3' | -1 bp | Some cell division |
| | 5'- CGCCACGTCGACAAAATCA-CCTGAGGT…TAACTCCGGCAGACGAGATCGAGCCACGTGGCGGCCGCGCGAAGG-3' | -1 bp | |
| #4 | 5'- --------------------CCTGAGGT…TAACTCCGGCAGACGAGATCGAGCCACGTGGCGGCCGCGCGAAGG-3 | -34 bp | No prehaustorium |
|  | 5'- --------------------CCTGAGGT…TAACTCCGGCAGACGAGATCGAGCCACGTGGCGGCCGCGCGAAGG-3 | -34 bp |  |
| #5 | 5'- CGCCACGTCGACAAAATCA-CCTGAGGT…TAACTCCGGCAGACGAGATCGAGCCACGTGGCGGCCGCGCGAAGG-3' | -1 bp | prehaustorium |
| | 5'- CGCCACGTCGACAAAATCACCCTGAGGT…TAACTCCGGCAGACGAGATCGAG-CACGTGGCGGCCGCGCGAAGG-3' | -1 bp | |
| #6 | 5'- CGCCACGTCGACAAAATCA-CCTGAGGT…TAACTCCGGCAGACGAGATCGAGCCACGTGGCGGCCGCGCGAAGG-3' | -1 bp | prehaustorium |
|  | 5'- CGCCACGTCGACAAAA----CCTGAGGT…TAACTCCGGCAGACGAGATCGAGCCACGTGGCGGCCGCGCGAAGG-3' | -4 bp |  |
| #7 | 5'- CGCCACGTCGACAAAATCA-CCTGAGGT…TAACTCCGGCAGACGAGATCGAGCCACGTGGCGGCCGCGCGAAGG-3' | -1 bp | prehaustorium |
| | 5'- CGCCACGTCGACAAAATCA-CCTGAGGT…TAACTCCGGCAGACGAGATCGAGCCACGTGGCGGCCGCGCGAAGG-3' | -1 bp | |
| #8 | 5'- CGCCACGTCGACAAAATCA-CCTGAGGT…TAACTCCGGCAGACGAGATCGAGCCACGTGGCGGCCGCGCGAAGG-3' | -1 bp | prehaustorium |
|  | 5'- CGCCACGTCGACAAAATCACCCTGAGGT…TAACTCCGGCAGACGAGATCGA----------------GCGAAGG-3' | -16 bp |  |
| #9 | 5'- CGCCACGTCGACAAAATCACCCTGAGGT…TAACTCCGGCAGACGAGATCGAG-CACGTGGCGGCCGCGCGAAGG-3' | -1 bp | prehaustorium |
| | 5'- CGCCACGTCGACAAAATCA-CCTGAGGT…TAACTCCGGCAGACGAGATCGAGCCACGTGGCGGCCGCGCGAAGG-3' | -1 bp | |
| #10 | 5'- CGCCACGTCGACAAAATCA--CTGAGGT…TAACTCCGGCAGACGAGATCGAGCCCACGTGGCGGCCGCGCGAAGG-3 | +1 bp, -2 bp | prehaustorium |
| | 5'- CGCCACGTCGACAAAATCACCCCTGAGGT…TAACTCCGGCAGACGAGATCGAG--ACGTGGCGGCCGCGCGAAGG-3 | -2 bp, +1 bp | |
|  | 5'- CGCCACGTCGACAAAATCA--CTGAGGT…TAACTCCGGCAGACGAGATCG-----CGTGGCGGCCGCGCGAAGG-3 | -5 bp, -2 bp |  |
| #11 | 5'- CGCCACGTCGACAAAATCACCCTGAGGT…TAACTCCGGCAGACGAG--------ACGTGGCGGCCGCGCGAAGG-3' | -8 bp | No prehaustorium |
| | 5'- CGCCACGTCGACAAAATCA-CCTGAGGT…TAACTCCGGCAGACGAGATCGAGCCACGTGGCGGCCGCGCGAAGG-3' | -1 bp | |
| #12 | 5'- CGCCACGTCGACAAAATCA-CCTGAGGT…TAACTCCGGCAGACGAGATCGAGCCACGTGGCGGCCGCGCGAAGG-3' | -1 bp | No prehaustorium |
| | 5'- CGCCACGTCGACAAAATCA-CCTGAGGT…TAACTCCGGCAGACGAGATCGAGCCACGTGGCGGCCGCGCGAAGG-3' | -1 bp | |
| #13 | 5'- CGCCACGTCGACAAAATCA-CCTGAGGT…TAACTCCGGCAGACGAGATCGAGCCACGTGGCGGCCGCGCGAAGG-3' | -1 bp | No prehaustorium |
| | 5'- CGCCACGTCGACAAAATCA-CCTGAGGT…TAACTCCGGCAGACGAGATCGAGCCACGTGGCGGCCGCGCGAAGG-3' | -1 bp | |
| #14 | 5'- -----------------------GAGGT…TAACTCCGGCAGA-GAGATCGAGCCACGTGGCGGCCGCGCGAAGG-3' | -1 bp, -52 bp | No prehaustorium |
|  | 5'- -----------------------GAGGT…TAACTCCGGCAGA-GAGATCGAGCCACGTGGCGGCCGCGCGAAGG-3' | -1 bp, -52 bp |  |
| #15 | 5'- -----------------------GAGGT…TAACTCCGGCAGACGAGATCGAGCCACGTGGCGGCCGCGCGAAGG-3' | -55 bp | Some cell division |
| | 5'- CGCCACGTCGACAAAATCA-CCTGAGGT…TAACTCCGGCAGACGAGATCGAGCCACGTGGCGGCCGCGCGAAGG-3' | -1 bp | |
|  | 5'- -----------------------GAGGT…----------------------------------CCGCGCGAAGG-3' | -985 bp, -55 bp |  |
| #16 | 5'- CGCCACGTCGACAAAATCACCCTGAGGT…TAACTCCGGCAGACGAGA-------ACGTGGCGGCCGCGCGAAGG-3' | -7 bp | No prehaustorium |
|  | 5'- CGCCAC--------------------GT…TAACTCCGGCAGACGAGA-------ACGTGGCGGCCGCGCGAAGG-3' | -7 bp, -20 bp |  |
| #17 | 5'- CGCCACGTCGACAAAATCACC-TGAGGT…TAACTCCGGCAGACGAGATCGAGCCACGTGGCGGCCGCGCGAAGG-3' | -1 bp | No prehaustorium |
|  | 5'- CGCCACGTCGACATAATCT---------…TAACTCCGGCAGACGAGATCGAGCCACGTGGCGGCCGCGCGAAGG-3' | A->T, +1 bp, -18 bp |  |
| #18 | 5'- CGCCACGTCGACAAAATCACC-------…TAACTCCGGCAGACGAGATCGAGCCACGTGGCGGCCGCGCGAAGG-3' | -29 bp | prehaustorium |
|  | 5'- CGCCACGTCGACAAAATCACC-------…TAACTCCGGCAGACGAGATCGAGCCACGTGGCGGCCGCGCGAAGG-3' | -29 bp |  |
| #19 | 5'- CGCCACGTCGACAAAATCACC-TGAGGT…TAACTCCGGCAGACGAGATCGAGCCACGTGGCGGCCGCGCGAAGG-3' | -1 bp | No prehaustorium |
|  | 5'- CGCCACGTCGAC----------TGAGGT…TAACTCCGGCAGACGAGATCGAGCCACGTGGCGGCCGCGCGAAGG-3' | -10 bp |  |
| #20 | 5'- CGCCACGTCGACAAAATC----TGAGGT…TAACTCCGGCAGACGAGATCGAGCCAACGTGGCGGCCGCGCGAAGG-3' | -4 bp, +1 bp | Some cell division |
| | 5'- CGCCACGTCGACAAAATCACC-TGAGGT…TAACTCCGGCAGACGAGATCGAGC-ACGTGGCGGCCGCGCGAAGG-3' | -1 bp, -1 bp | |
| | 5'- CGCCACGTCGACAAAATCACC-TGAGGT…TAACTCCGGCAGACGAGATCGAGCCACGTGGCGGCCGCGCGAAGG-3' | -1 bp | |
| PjRGFR1 | | | |
| WT #1 | 5'-CGGTTCCATCCCTGCAGTTCTGTCCAATGCCA…ACAGCGAAAACAGGGGACCGAGGTTTTGGGCAGGA-3' | N/A | |
| | 5'-CGGTTCCATCCCTGCAGTTCTGTCCAATGCCA…ACAGCGAAAACAGGGGACCGAGGTTTTGGGCAGGA-3' | N/A | |
| WT #2 | 5'- ACCTCCAGGTTTTGTCGCTTTACACGA…CTCCGGTTCCATCCCTGCAGTTCTGTCCAATGCCACGAG-3' | N/A | |
| | 5'- ACCTCCAGGTTTTGTCGCTTTACACGA…CTCCGGTTCCATCCCTGCAGTTCTGTCCAATGCCACGAG-3' | N/A | |
| #1 | 5'--------------------------------…ACAGCGAAAACAGGGGAACCGAGGTTTTGGGCAGGA-3' | -1035 bp, +1 bp | Prehaustorium |
|  | 5'-CGGTTCCATCCCTGC-GTTCTGTCCAATGCCA…----------------------GTTTTGGGCAGGA-3' | -1 bp, -1371 bp |  |
|  | 5'--------------------------------…----------------------GTTTTGGGCAGGA-3' | -35 bp, -1371 bp |  |
| #2 | 5'-CGGTTCC-------------TGTCCAATGCCA…ACAGCGAAAACAGGGGAACCGAGGTTTTGGGCAGGA-3' | -13 bp, +1 bp | Some cell division |
|  | 5'-CGGTTCC-------------TGTCCAATGCCA…ACAGCGAAAACAGGGGAACCGAGGTTTTGGGCAGGA-3' | -13 bp, +1 bp |  |
| #3 | 5'-CGGTTCCATCCCTGC-----------------…ACAGCGAAC---------------------------3' | -125 bp, +1 bp, -73 bp | no prehaustorium |
| | 5'-CGGTTCCATCCCTGCAAGTTCTGTCCAATGCCA…ACAGCGAAAACAGGGGAACCGAGGTTTTGGGCAGGA-3' | +1 bp, +1 bp | |
|  | 5'-CGGTTCCATCCCTGC-----------------…ACAGCGAAAACAGGGGAACCGAGGTTTTGGGCAGGA-3' | -125 bp, +1 bp |  |
| #4 | 5'-CGGTTCCATCCCTGCAAGTTCTGTCCAATGCCA…ACAGCGAAAACAGGGGACCGAGGTTTTGGGCAGGA-3'(+1 bp) | +1 bp | Some cell division |
|  | 5'-CGGTTCCATCCCTGC-----TGTCCAATGCCA…ACAGCGAAAACAGGGGACCGAGGTTTTGGGCAGGA-3'(-5 bp) | -5 bp |  |
| | 5'-CGGTTCCATCCCTGCAGTTCTGTCCAATGCCA…ACAGCGAAAACAGGGGAACCGAGGTTTTGGGCAGGA-3'(+1 bp) | +1 bp | |
| #5 | 5'-CGGTTCCATCCCTGCA-TTCTGTCCAATGCCA…ACAGCGAAAACAGGG-ACCGAGGTTTTGGGCAGGA-3' | -1 bp, -1 bp | Prehaustorium |
|  | 5'-CGGTTCCATCCCTGC----CTGTCCAATGCCA…ACAGCGAAAACAGGGGACCCCACAATACCAGA-------TGGGCAGGA-3' | -4 bp, +13 bp, -7 bp |  |
| #6 | 5'-GATTTTCAAAGCTCC-----------------…ACAGCGAAAACA----CCGAGGTTTTGGGCAGGA-3' | +15 bp, -1164 bp, -5 bp | no prehaustorium |
| | 5'-CGGTTCCATCCCTGCAAGTTCTGTCCAATGCCA…ACAGCGAAAACAGGGGAGCTGCCTGACTGTACTATAGTATTATTAGCCACACCGAGGTTTTGGGCAGGA-3' | +1 bp, +34 bp | |
| | 5'-CGGTTCCATCCCTGCAGTTCTGTCCAATGCCA…ACAGCGAAAACAGGGGAGCTGCCTGACTGTACTATAGTATTATTAGCCACACCGAGGTTTTGGGCAGGA-3' | +34 bp | |
| #7 | 5'-CGGTTCCATCCCTGC---------CAATGCCA…ACAGCGAAAACAGGGGAACCGAGGTTTTGGGCAGGA-3' | -9 bp, +1 bp | no prehaustorium |
| | 5'-CGGTTCCATCCCTGCA---CTGTCCAATGCCA…ACAGCGAAAACAGGGGAACCGAGGTTTTGGGCAGGA-3' | -3 bp, +1 bp | |
| #8 | 5'-CGGTTCCATCC-----GT---GTCCAATGCCA…ACAGCGAAAACAGGGGAACCGAGGTTTTGGGCAGGA-3' | -8 bp, +1 bp | no prehaustorium |
| | 5'-CGGTTCCATCCCTGCA-TTCTGTCCAATGCCA…ACAGCGAAAACAGGGGAACCGAGGTTTTGGGCAGGA-3' | -8 bp, +1 bp | |
| #9 | 5'-CGGTTCCATCCCTGCAAGTTCTGTCCAATGCCA…ACAGCGAAAACAGGGGAACCGAGGTTTTGGGCAGGA-3' | +1 bp, +1 bp | Some cell division |
|  | 5'-CGGTTCCATCCCTGC-----TGTCCAATGCCA…ACAGCGAAAACAGGGGAACCGAGGTTTTGGGCAGGA-3' | -5 bp, +1 bp |  |
|  | 5'-CGGTTCCATCCCTGCT----------------…ACAGCGAAAACAGGGGACCGAGGTTTTGGGCAGGA-3' | +1 bp, -1308 bp |  |
|  | 5'-CGGTTCCATCCCTGC-GTTCTGTCCAATGCCA…---------------------GGTTTTGGGCAGGA-3' | -1 bp, -1372 bp |  |
| #10 | 5'-CGGTTC--------------TGTCCAATGCCA…ACAGCGAAAACAGGGGACCGAGGTTTTGGGCAGGA-3' | -14 bp | Prehaustorium |
|  | 5'-CGGTTCCATCCCTGC---------CAATGCCA…ACAGCGAAAACAGGGGACCGAAGGTTTTGGGCAGGA-3' | -9 bp, +1 bp |  |
| | 5'-CGGTTCCATCCCTGCAGTTCTGTCCAATGCCA…ACAGCGAAAACAGGGGACCGAAGGTTTTGGGCAGGA-3' | +1 bp | |
| #11 | 5'-CGGTTCCATCCCTGCAGTTCTGTCCAATGCCA…ACAGCGAAAACAGGGGACCGAAGGTTTTGGGCAGGA-3' | +1 bp | Some cell division |
|  | 5'-CGGTTCCATCCCTGC-----TGTCCAATGCCA…ACAGCGAAAACAGGGGACCGAAGGTTTTGGGCAGGA-3' | -5 bp, +1 bp |  |
| #12 | 5'-CGGTTCCATCCCTGCAGTTCTGTCCAATGCCA…ACAGCGAAAACAGGGGAACCGAGGTTTTGGGCAGGA-3' | +1 bp | Prehaustorium |
|  | 5'-CGGTTCCATCCCTGCAGT-----CCAATGCCA…ACAGCGAAAACAGGGGATCCGAGGTTTTGGGCAGGA-3' | -5 bp, +1 bp |  |
|  | 5'-CGGTTCCATCCCTG--------TCCAATGCCA…ACAGCGAAAACAGGGGTCATCT-GGTATTGGGTAGGA-3' | -8 bp, +1 bp, -1 bp |  |
| #13 | 5'-CGGTTCCATCCCTGC-GTTCTGTCCAATGCCA…ACAGCGAAAACAGGGGACCGAGGTTTTGGGCAGGA-3' | -1 bp | Some cell division |

| | | | |
|---|---|---|---|
| | 5'-CGGTTCCATCCCTGC-GTTCTGTCCAATGCCA…ACAGCGAAAACAGGGGAC-GAGGTTTTGGGCAGGA-3' | -1 bp, -1 bp | |
| #14 | 5'-CGGTTCCATCCCTGCAGTTCTGTCCAATGCCA…ACAGCGAAAACAGGGGAACCGAGGTTTTGGGCAGGA-3' | +1 bp | Prehaustorium |
| | 5'-CGGTTCCATCCCTGCAGTTCTGTCCAATGCCA…ACAGCGAAAACAGGGGA--GAGGTTTTGGGCAGGA-3' | -2 bp | |
| #15 | 5'-CGGT-------------------ACAATGCCA…ACAGCGAAAACAGGGGAACCGAGGTTTTGGGCAGGA-3' | -19 bp, +1 bp, +1 bp | Prehaustorium |
| | 5'-CGGTTCCATCCCTGCAGTTCTGTCCAATGCCA…ACAGCGAAAACAGGGGAACCGAGGTTTTGGGCAGGA-3' | +1 bp | |
| #16 | 5'-CGGTTCCATCCCTGCAGT——-CCAATGCCA…ACAGCGAAAACAGGGGAACCGAGGTTTTGGGCAGGA-3' | -5 bp, +1 bp | no prehaustorium |
|  | 5'-CGGTTCCATCCCTGCAGT-----CCAATGCCA…ACAGCGAAAACAGGGGAACCGAGGTTTTGGGCAGGA-3' | -5 bp, +1 bp |  |
| #17 | 5'-CGGTTCCATCCCTGCAGTTCTGTCCAATGCCA…ACAGCGAAAACAGGGGAACCGAGGTTTTGGGCAGGA-3' | +1 bp | Prehaustorium |
| | 5'-CGGTTCCATCCCTGCAGTTCTGTCCAATGCCA…ACAGCGAAAACAGGGGAACCGAGGTTTTGGGCAGGA-3' | +1 bp | |
| #18 | 5'-CGGTTCCATCCCTGCAAGTTCTGTCCAATGCCA…----------------ACCGAGGTTTTGGGCAGGA-3' | +1 bp, -53 bp | Prehaustorium |
| | 5'-CGGTTCCATCCCTGCA-TTCTGTCCAATGCCA…ACAGCGAAAACAGGGGACCGAGGTTTTGGGCAGGA-3' | -1 bp | |
| #19 | 5'- ACCTCCAGGTTTTGTCGCTTTACACGA…CTCCGGTTCCATCCCTGCAGT-----CCAATGCCACGAG-3' | -5 bp | Prehaustorium |
|  | 5'- ACCTCCAGGTTTTGTCGCTTTACACGA…CTCCGGTTC--------------TGTCCAATGCCACGAG-3' | -14 bp |  |
| #20 | 5'- ACCTCCAGGTTTTGTCGCTTTACACGA…CTCCGGTTC--------------TGTCCAATGCCACGAG-3' | -14 bp | Aborted prehaustorium |
|  | 5'- ACCTCCAGGTTTTGTCGCTTTACACGA…CTCCGGTTC--------------TGTCCAATGCCACGAG-3' | -14 bp |  |
| #21 | 5'- ACCTCCAGGTTTTGTCGCTTTACACGA…CTCCGGTTCCATCCCTGCAGTTTCTGTCCAATGCCACGAG-3'(+1 bp) | +1 bp | no prehaustorium |
| | 5'- ACCTCCAGGTTTTGTCGCTTTACACGA…CTCCGGTTCCATCCCTGC--TTCTGTCCAATGCCACGAG-3'(-2 bp) | -2 bp | |
| | 5'- ACCTCCAGGTTTTGTCGCTTTACACGA…CTCCGGTTCCATCCCTGCA-TTCTGTCCAATGCCACGAG-3'(-1 bp) | -1 bp | |
| PjRGFR4 | | | |
| WT | 5'- AACGTGCCGGCAGATATTGGGCGGTTGAAGAAT… TGGAATAAATTATCTGGCGATATCCCAGAGGAGC-3' | N/A | |
| | 5'- AACGTGCCGGCAGATATTGGGCGGTTGAAGAAT… TGGAATAAATTATCTGGCGATATCCCAGAGGAGC-3' | N/A | |
| #1 | 5'- AACGTGCCGGCAGATATTTGGGCGGTTGAAGAAT… TGGAATAAATTATCTGGCGATATCC-AGAGGAGC-3' | +1 bp, -1 bp | Prehaustorium |
| | 5'- AACGTGCCGGCAGATATTTGGGCGGTTGAAGAAT… TGGAATAAATTATCTGGCGATATCCCCAGAGGAGC-3' | +1 bp, +1 bp | |
| #2 | 5'- AACGTGCCGGCAGATATTTGGGCGGTTGAAGAAT… TGGAATAAATTATCTGGCGATATCCTCAGAGGAGC-3' | +1 bp, +1 bp | Prehaustorium |
| | 5'- AACGTGCCGGCAGATATTTGGGCGGTTGAAGAAT… TGGAATAAATTATCTGGCGATATCC-AGAGGAGC-3' | +1 bp, -1 bp | |
| #3 | 5'- AACGTGCCGGCAGATAT-GGGCGGTTGAAGAAT… --------------------------AGAGGAGC-3' | -1 bp, -31 bp | aborted prehaustorium |
|  | 5'- AACGTGCCGGCAGATAT-GGGCGGTTGAAGAAT… --------------------------AGAGGAGC-3' | -1 bp, -31 bp |  |
| #4 | 5'- AACGTGCCGGCAGATATTTGGGCGGTTGAAGAAT… TGGAATAAATTATCTGGCGATATCCACAGAGGAGC-3' | +1 bp, +1 bp | Prehaustorium |
| | 5'- AACGTGCCGGCAGATATTTGGGCGGTTGAAGAAT… TGGAATAAATTATCTGGCGATATCCACAGAGGAGC-3' | +1 bp, +1 bp | |
| #5 | 5'- AACGTGCCGGCAGATA--GGGCGGTTGAAGAAT… TGGAATAAATTATCTGGCGATATCCCAGAGGAGC-3' | -2 bp | Prehaustorium |
| | 5'- AACGTGCCGGCAGATA--GGGCGGTTGAAGAAT… TGGAATAAATTATCTGGCGATATCCCAGAGGAGC-3' | -2 bp | |
| #6 | 5'- AACGTGCCGGCAGATAT-GGGCGGTTGAAGAAT… TGGAATAAATTATCTGGCGATATCCCAGAGGAGC-3' | -1 bp | Prehaustorium |
|  | 5'- AACGTGCCGGCAGAT---------TTGAAGAAT… TGGAATAAATTATCTGGCGATATCCCAGAGGAGC-3' | -11 bp |  |
| | 5'- AACGTGCCGGCAGATAA---GGCGGTTGAAGAAT… TGGAATAAATTATCTGGCGATATCCCAGAGGAGC-3' | +1 bp, -3 bp | |
| #7 | 5'- AACGTGCCGGCAGATAT-GGGCGGTTGAAGAAT… TTCAGTT--------------------------------GC-3' | -1 bp, +6bp, -32 bp | Prehaustorium |
|  | 5'- AACGTGCCGGCAGATAT-GGGCGGTTGAAGAAT… TTCAGTT--------------------------------GC-3' | -1 bp, +6bp, -32 bp |  |
| #8 | 5'- AACGTGCCGGCAGATAT-GGGCGGTTGAAGAAT… TGGAATAAATTATCTGGCGATATCCCAGAGGAGC-3' | -1 bp | Prehaustorium |
|  | 5'- AACGTGCCGGCAG-----------TTGAAGAAT… TGGAATAAATTATCTGGCGATATCCCAGAGGAGC-3' | -11 bp |  |
| #9 | 5'- AACGTGCCGGCAGATA--GGGCGGTTGAAGAAT… TGGAATAAATTATCTGGCGATATCCCAGAGGAGC-3' | -2 bp | Prehaustorium |
|  | 5'- AACGTGCCGGCAGATAT----------------… ------------------------CCAGAGGAGC-3' | -424 bp |  |
| #10 | 5'- AACGTGCCGGCAGATA----GCGGTTGAAGAAT… TGGAATAAATTATCTGGCGATATCCCAGAGGAGC-3' | -4 bp | Prehaustorium |
|  | 5'- AACGTGCCGGCAGATA----GCGGTTGAAGAAT… TGGAATAAATTATCTGGCGATATC--AGAGGAGC-3' | -4 bp, -2 bp |  |
| #11 | 5'- AACGTGCCGGCAGATA----GCGGTTGAAGAAT… TGGAATAAATTATCTGGCGATATCC-AGAGGAGC-3' | -4 bp, -1 bp | Prehaustorium |
|  | 5'- AACGTGCCGGCAGAT------CGGTTGAAGAAT… TGGAATAAATTATCTGGCGATATCC-AGAGGAGC-3' | -6 bp, -1 bp |  |
| #12 | 5'- AACGTGCCGGCAGATATTG--------AAGAAT… TGGAATAAATT CCGGGCCTACATTAGTGAGCTCCTCTGATTCCAGCTCGCTGGAATAAT----------------------CTCTA-3' | -8 bp, +48 bp, -22 bp | Prehaustorium |
|  | 5'- AACGTGCCGGCAGAT----GGCGGTTGAAGAAT… TGGAATAAATTATCTGGCGATATCCCAGAGGAGCTCTG-3' | -4 bp, A->G |  |
| #13 | 5'- AACGTGCCGGCAGATATTTGGGCGGTTGAAGAAT… TGGAATAAATTATCTGGCGATATCC-AGAGGAGC-3' | +1 bp, -1 bp | Prehaustorium |
| | 5'- AACGTGCCGGCAGATATTTGGGCGGTTGAAGAAT… TGGAATAAATTATCTGGCGATATCCCAGAGGAGC-3' | +1 bp, -1 bp | |
| #14 | 5'- AACGTGCCGGCAGATA----GCGGTTGAAGAAT… TGGAATAAATTATCTGGCGATATCCCAGAGGAGC-3' | -4 bp | Prehaustorium |
|  | 5'- AACGTGCCGGCAGA----GGGCGGTTGAAGAAT… TGGAATAAATTATCTGGCGATATCCCAGAGGAGC-3' | -4 bp |  |
| #15 | 5'- AACGTGCCGGCAGATATTTGGGCGGTTGAAGAAT… TGGAATAAATT---TGGCGATATCCCAGAGGAGC-3' | +1 bp, -3 bp | Prehaustorium |
|  | 5'- AACGTGCCGGCAGATATT---------------… ----------------------------------3' | -16 bp, -387 bp |  |
| #16 | 5'- AACGTGCCGGCAGATATTAGGGCGGTTGAAGAAT… TGGAATAAATTATCTGGCGATATCCCAGAGGAGC-3' | +1 bp, -3 bp | Some cell division |
|  | 5'- AACGTGCCGGC---------GCGGTTGAAGAAT… TGGAATAAATTATCTGGCGATATCC-AGAGGAGC-3' | -9 bp, -1 bp |  |
| | 5'- AACGTGCCGGCAGATATTAGGGCGGTTGAAGAAT… TGGAATAAATTATCTGGCGATATCC-AGAGGAGC-3' | +1 bp, -1 bp | |
| #17 | 5'- AACGTGCCGGCAGATA--GGGCGGTTGAAGAAT… TGGAATAAATTATCTGGCGATATCCTCAGAGGAGC-3' | -2 bp, +1 bp | Prehaustorium |
| | 5'- AACGTGCCGGCAGATATTTGGGCGGTTGAAGAAT… TGGAATAAATTATCTGGCGATATCCCAGAGGAGC-3' | +1 bp | |
| #18 | 5'- AACGTGCCGGCAGATATT---------------… -------------------------CAGAGGAGC-3' | -424 bp | Prehaustorium |
|  | 5'- AACGTGCCGGCAGATATT---------------… -------------------------CAGAGGAGC-3' | -424 bp |  |
| #19 | 5'- AACGTGCCGGCAGATATTTGGGCGGTTGAAGAAT… TGGAATAAATTATCTGGCGATATCCCAGAGGAGC-3' | +1 bp | Prehaustorium |
| | 5'- AACGTGCCGGCAGATATTTGGGCGGTTGAAGAAT… TGGAATAAATTATCTGGCGATATCCCAGAGGAGC-3' | +1 bp | |
| #20 | 5'- AACGTGCCGGCAGAT-----GCGGTTGAAGAAT… --------------------------AGAGGAGC-3' | -5 bp, -31 bp | No prehaustorium |
|  | 5'- AACGTGCCGGCAGAT-----GCGGTTGAAGAAT… --------------------------AGAGGAGC-3' | -5 bp, -31 bp |  |
| #21 | 5'- AACGTGCCGGCAGAATA------GGTTGAAGAAT… TGGAATAAATTATCTGGCGATATC-----------3' | +1 bp, -6 bp, -15 bp | Prehaustorium |
|  | 5'- AACGTGCCGGCAGAATA------GGTTGAAGAAT… TGGAATAAATTATCTGGCGATATC-----------3' | +1 bp, -6 bp, -15 bp |  |
| #22 | 5'- AACGTGCCGGCAGATAT-GGGCGGTTGAAGAAT… TGGAATAAATTATCTGGCGATATCCCCAGAGGAGC-3' | -1 bp, +1 bp | Prehaustorium |
|  | 5'- AACGTGCCGGCAGATAT-GGGCGGTTGAAGAAT… ----------------------------------3' | -1 bp, -169 bp |  |
| #23 | 5'- AACGTGCCGGCAGATA--GGGCGGTTGAAGAAT… TGGAATAAATTATCTGGCGATATCCCAGAGGAGC-3'(-2 bp) | -2 bp | Prehaustorium |
| | 5'- AACGTGCCGGCAGATA--GGGCGGTTGAAGAAT… TGGAATAAATTATCTGGCGATATCCCAGAGGAGC-3'(-2 bp) | -2 bp | |

Supplemental Table S6. Primers used in this study

| Primer name | Sequence | Description | Reference |
|---|---|---|---|
| oMRF698 | attcaagcttggaggagctctagtgcaaaatttgcacagcaac | PjRGF3 promoter fwd hifi | |
| oMRF699 | tgctcaccattgagctagcttaatgttattgtaatgaggggtcaagtt | PjRGFR3 promoter rev hifi | |
| oMRF815 | TTGATGAAGATGGCGAATACGA | PjRGF3 qPCR primer sense | |
| oMRF816 | AATCTGCATTGAAGGCAACAAA | PjRGF3 qPCR primer antisense | |
| oMRF817 | ACTCATCCTCCGGTCCATAA | PjRGF4 qPCR primer sense | |
| oMRF818 | TTCCCAAAGTACCCTCGAAATC | PjRGF4 qPCR primer antisense | |
| oMRF819 | GGCTCTTGAAATCGCGTTAAAT | PjRGFR1 qPCR primer sense | |
| oMRF820 | GTGAAAGGTCCAATACGGAGAG | PjRGFR1 qPCR primer antisense | |
| oMRF821 | GGGAGTTACGGCGAATTGT | PjRGFR2 qPCR primer sense | |
| oMRF822 | CCCGAGTGTTGATGGGATTAG | PjRGFR2 qPCR primer antisense | |
| oMRF823 | AACAGGCTTGGTGGTACTATTC | PjRGFR3 qPCR primer sense | |
| oMRF824 | CCCGAAATCGAAGGAGGTATG | PjRGFR3 qPCR primer antisense | |
| oMRF825 | GGGCGAGTAGTAATAAGCTGAC | PjRGFR4 qPCR primer sense | |
| oMRF826 | GGAAATCTCGGGCGGTATAAA | PjRGFR4 qPCR primer antisense | |
| oMRF827 | GATTTGCAGTTGCTCGATGTG | PjRGFR6 qPCR primer sense | |
| oMRF828 | GCACTCAGGTTCAAGAGTATGT | PjRGFR6 qPCR primer antisense | |
| oMRF829 | GAACCAATTATCGGGCCAAATC | PjRGFR5 qPCR primer sense | |
| oMRF830 | CCCAAATGCGGGAGGTATAG | PjRGFR5 qPCR primer antisense | |
| oMRF831 | CAGAGTGGTGGCAGTCATATTCCTTC | PjPLT promoter fwd | |
| oMRF832 | GGGTTTCAATGTTGTCATTGACTAACCC | PjPLT promoter rev | |
| oMRF833 | ATGAATTCCAACAACTGGCTTTCTTTTC | PjPLT gene fwd | |
| oMRF834 | TCATTCATTCCACATGCTGAAAACAC | PjPLT gene rev | |
| oMRF835 | aattcaagcttggaggagctgaacctgggcgtaaaacc | PjPLT promoter fwd hifi for cel5 sacI | |
| oMRF836 | ttgctcaccattttcttttttttttggtgccc | PjPLT promoter rev hifi for mNG | |
| oMRF837 | aaaaaagaaaatggtgagcaagggagag | mNG forward for 21464 hifi | |
| oMRF838 | ttggaattcatacctgaagatccCTTGTAAAGCTCGTCCATTCC | mNG rev for 21464 hifi | |
| oMRF839 | gatcttcaggtatgaattccaacaactggCTT | PjPLT gene fwd hifi for mNG | |
| oMRF840 | catcttcatataagcgagcttcattcattccacatgctgAAAAC | PjPLT gene rev hifi for cel5 sacI | |
| oMRF849 | CTCTTCATCAACCGGTCCTAAA | SH14Contig_4244 (ShRGF1) sense | |
| oMRF850 | TTCGACTTGTTTCGATCTCCTC | SH14Contig_4244 (ShRGF1) antisense | |
| oMRF869 | CAGGATGTTAAGGAGATGGACAA | PjRGF1 qPCR primer sense | |
| oMRF870 | GGGATTGGCTTCAGTAGGTTTA | PjRGF1 qPCR primer antisense | |
| oMRF884 | GCTATTCCACGTGGTTCCACCT | PjRGFR1 pro fwd primer | |
| oMRF885 | GGCATCGGCATTTGGACAATGT | PJRGFR1 pro rev primer | |
| oMRF886 | aattcaagcttggaggagcttccacgtggttccaccttaaaatc | PjRGFR1 pro fwd primer for hifi | |
| oMRF887 | cccttgctcaccattgagctcatcggcatttggacaatgt | PjRGFR1 pro rev primer for hifi | |
| oMRF888 | gaaGGCAAAGCAATCCTCCGGAa | PjRGFR2 pro fwd primer | |
| oMRF889 | GTGGCGTTCAAGAATTGCCTCG | PjRGFR2 pro rev primer | |
| oMRF890 | aattcaagcttggaggagctggggttgtccaccatgaag | PjRGFR2 pro fwd primer for hifi | |
| oMRF891 | cccttgctcaccattgagctttggggttttgcaaaaattgtt | PjRGFR2 pro rev primter for hifi | |
| oMRF892 | CTGGTCACCGGAGTTTCTTACT | PjRGFR6 pro fwd primer | |
| oMRF893 | GATGAAGTTTGTGCTTGCCTTGT | PjRGFR6 pro rev primer | |
| oMRF894 | aattcaagcttggaggagctctggtcaccggagtttcttac | PjRGFR6 pro fwd primer for hifi | |
| oMRF895 | cccttgctcaccattgagctgatgaagtttgtgcttgccttgt | PjRGFR6 pro rev primer for hifi | |
| oMRF896 | TGAATATGAGTAGGCTACATGACATGACTT | PjRGFR5 pro fwd primer | |
| oMRF897 | caTCTTGAAAATGGATGTCAACGAAAGTC | PjRGFR5 pro rev primer | |
| oMRF898 | aattcaagcttggaggagctggctacatgacatgacttatggtg | PjRGFR5 pro fwd primer for hifi | |
| oMRF899 | cccttgctcaccattgagctcttgaaaatggatgtcaacgaaagt | PjRGFR5 pro fwd primer for hifi | |
| oMRF950 | ccacccccacacactcaggTgtcagtaaaggtgaagctgTGAT | mSCARLET fwd for hifi Pjv1_00019181 signal peptide | |
| oMRF951 | ttaaatctatccgcgccgtttttatacaactcatccattccacctg | mSCARLET rev for hifi Pjv1_00019181 signal peptide | |
| oMRF952 | acctgagtgtgtgggggtg | Pjv1_00019181 GPI anchor peptide rev | |
| oMRF953 | aacggcgcggatagatttaaggg | Pjv1_00019181 GPI anchor peptide fwd | |
| oMRF955 | CAGCAAATCTCTCTAATCGACAGC | PjRGF2 promoter fwd | |
| oMRF956 | CCTGAATGATGTATTTCAGCTGT | PjRGF2 promoter rev | |
| oMRF957 | attcaagcttggaggagctccagcaaatctctctaatcgaca | PjRGF2 promoter fwd for hifi | |
| oMRF958 | cccttgctcaccattgagctcacatactctcgatatatataaacaaagg | PjRGF2 promoter rev for hifi | |
| oMRF959 | TTCCATGCAAAGGAGTTTAACAG | PjRGF2 qPCR sense | |
| oMRF960 | ACCGAACGGCGCATTAT | PjRGF2 qPCR antisense | |
| oMRF965 | ACCATAATTGGACCGCCTGA | qPCR for PjUBC sense | 13 |
| oMRF966 | ACAGTTGGGGGACTGTTTGG | qPCR for PjUBC antisense | |
| oMRF1079 | ATCTTTGCTCAAGCCGTCAATCCACAGGGCG | PjRGFR4 ectodomain fwd primer in-fusion | |
| oMRF1080 | CACACTACCAGCAAGTCTAGCAGCCCTTCTTCTGCTGGCAGCTGCGCTGCCAC | PjRGFR4 ectodomain rev primer in-fusion | |
| oMRF1081 | ATCTTTGCTCAAGCCCTCTCTCTAGACGGACAAGCCCT | PjRGFR5 ectodomain fwd primer in-fusion | |
| oMRF1082 | CACACTACCAGCAAGGTTTTTGGCCGACTTAAACCCGT | PjRGFR5 ectodomain rev primer in-fusion | |
| oMRF1083 | ATCTTTGCTCAAGCCATTAACGACCAGGGCCAAGCTC | PjRGFR3 ectodomain fwd primer in-fusion | |
| oMRF1084 | CACACTACCAGCAAGCATGGCGGCCTTCGCGC | PjRGFR3 ectodomain rev primer in-fusion | |
| oMRF1085 | ATCTTTGCTCAAGCCGCAAACACCGAAGCTATTGTTTTGTTTTCATGG | PjRGFR2 ectodomain fwd primer in-fusion | |
| oMRF1086 | CACACTACCAGCAAGTAGCCTCCACTTCCGCCCG | PjRGFR2 ectodomain rev primer in-fusion | |
| oMRF1087 | ATCTTTGCTCAAGCCGCAAACACCAACCCAGAAGTCGAC | PjRGFR1 ectodomain fwd primer in-fusion | |
| oMRF1088 | CACACTACCAGCAAGCATTTTTAGCCTCCACGACTGTTTAAAATTACG | PjRGFR1 ectodomain rev primer in-fusion | |
| oMRF1089 | ATCTTTGCTCAAGCCTTGAACCATGAAGGCCTTTCTCTTCTG | PjRGFR6 ectodomain fwd primer in-fusion | |
| oMRF1090 | CACACTACCAGCAAGGATATATTCCCATTTCCTTCTGTGAATTTCCTGA | PjRGFR6 ectodomain rev primer in-fusion | |
| oMRF1110 | ATACAGTGGTCGAGCCTTAC | Sh TUB1 sense | 67 |
| oMRF1111 | TAGGTGTCGTGAGCTTAAGC | Sh TUB1 antisense | |
| oMRF1112 | tgtgccgaattcaagcttgggaacctgggcgtaaaacc | PjPLT promoter forward for BsmbI hifi | |
| oMRF1113 | cctcgcccttgctcacCATTtttcttttttttttggtgcccTAA | PjPLT promoter forward for BsmbI hifi | |
| oMRF1195 | CTTGCAGAGACCGACTTGTT | Sh14Contig_42422 (ShRGF2) sense | |
| oMRF1196 | TCCGTGTCTGCATGATTTAGG | Sh14Contig_42422 (ShRGF2) antisense | |
| oMRF1197 | CTCAACGAGCAAGAACAATGAC | Sh14Contig_2998 (ShRGF4) sense | |
| oMRF1198 | GGCGTTTGCTATCGAGGAT | Sh14Contig_2998 (ShRGF4) antisense | |
| oMRF1199 | CTGCAGCACTTGCAACATT | Sh14Contig_71709 (ShRGF3) sense | |
| oMRF1200 | GAAAGAAAGATTGGCCAAAGGG | Sh14Contig_71709 (ShRGF3) antisense | |
| oMRF1201 | GGAAGATCGGTTGTGTTCAGTT | ShRGF5 sense | |
| oMRF1202 | CCTTGCCATTGGTGGTTACA | ShRGF5 antisense | |
| oMRF1203 | TCAAGAAGGCGAACGAGATAAA | ShRGF6 sense | |
| oMRF1204 | TGATGAATCCAGTGCTGACC | ShRGF6 antisense | |
| oMRF1438 | GCTTGTAATCGTTTCCCATATATAGTTCAGTTTGC | PjRGF5 pro 1/2 fwd | |
| oMRF1439 | TTTAAGCAACCCAGCTAATATAGTGCTATAAATCTTGC | PjRGF5 pro 1/2 rev | |
| oMRF1440 | ccgaattcggatccggGCTTGTAATCGTTTCCCATATATAGTTCAG | PjRGF5 pro fwd for infusion | |
| oMRF1441 | gcccttgctcacCATTTTTAAGCAACCCAGCTAATATAGTGC | PjRGF5 pro rev for infusion | |
| oMRF1442 | ATGTCGACAATCTTATCGTACTTTCTTGTGG | PjRGF5 CDS fwd | |
| oMRF1443 | TCAATTGTGGATCGGAGGTCTCCC | PjRGF5 CDS rev | |
| oMRF1444 | CCCAAACCAGACACCAGAAA | PjRGF5 qPCR sense | |
| oMRF1445 | CCAAATCGGTCGGGATATGTT | PjRGF5 qPCR antisense | |
| oMRF1463 | CTCACGCCTAGGAGACC | PjRGF5 for colony PCR | |
| oMRF1464 | CTTGTTTAACACTCGTGTATTACACGAGATTTATC | PjRGF5 pro 5 fwd | |
| oMRF1465 | TTTAAGCAACCCAACTAATATAGTGCTATAAATCTTG | PjRGF5 pro 5 rev | |
| oMRF1466 | ccgaattcggatccggCTTGTTTAACACTCGTGTATTACACGAGAtttATC | PjRGF5 pro 5 fwd infusion | |
| oMRF1467 | gcccttgctcacCATTTTTAAGCAACCCAACTAATATAGTGCTATAAATCTTG | PjRGF5 pro 5 rev infusion | |
| oMRF1471 | CACCATGTCGACAATCTTATCGTACTTTCTTGTGG | PjRGF5 CDS fwd for pENTR | |
| oMRF1490 | ATGGCTGTTTTGCCAACCAAGC | PjRGF6 CDS fwd | |
| oMRF1491 | CTAATTATGAATCGGAGTTTTCTTTCGTGCAGG | PjRGF6 CDS rev | |
| oMRF1492 | ATGAGGTCTTGGATGTTGATATCTCTGGTAC | PjRGF7 CDS fwd | |
| oMRF1493 | TTAGTTGTGTATAGGAGGTTTTCTTCTTGCCT | PjRGF7 CDS rev | |
| oMRF1494 | ATGTCGTCAATCTTGTCGTCTCTTGTG | PjRGF8 CDS fwd | |
| oMRF1495 | TCAATTGTGAACTGGTGGGTGC | PjRGF8 CDS rev | |
| oMRF1496 | ATGGTGAGGTTTACTATTTTCATTTTCATGATTTTGC | PjRGF10 CDS fwd | |

| | | |
|---|---|---|
| oMRF1497 | TCAGTTATTCTTCGGAGGATGCGACT | PjRGF10 CDS rev |
| oMRF1498 | ATGGTGAAAATTGTATCTGATAAGAAAGAATTTGCAG | PjRGF10-2 CDS fwd |
| oMRF1499 | TCAGTTATTCTTCGGAGGATGCGAC | PjRGF10-2 CDS rev |
| oMRF1500 | ATGGAGTTTAAAAGTTTACTACTAAATGTTCTAATCATTTTGCTTG | PjRGF11 CDS fwd |
| oMRF1501 | TTAATTATTTGGCTGGTGATTGTGGATTGG | PjRGF11 CDS rev |
| oMRF1502 | ATGGGCAAAAATCTCTCAAACAATATGGGT | PjRGF12 CDS fwd |
| oMRF1503 | TTAAGGGGCAGAATTAACTGGGATATGCT | PjRGF12 CDS rev |
| oMRF1504 | ATGGAACTATCAAAAACCCTAATTATAGCTACATCCT | PjRGF17 CDS fwd |
| oMRF1505 | TTATGGGACGAATGGGATTTGCTTG | PjRGF17 CDS rev |
| oMRF1506 | CCGCTCTCAAGGAAGAAGTTTA | PjRGF6 sense #1 |
| oMRF1507 | TAATCCATGGCACCCAAGTC | PjRGF6 antisense #1 |
| oMRF1508 | CATCTCAACCCGCTCTCAA | PjRGF6 sense #2 |
| oMRF1509 | CTCCTTCCTTGCAACCATTTC | PjRGF6 antisense #2 |
| oMRF1510 | CGGAAATGGCAAGGAAAGATG | PjRGF7 sense #1 |
| oMRF1511 | ATGTCCAATGTGTCTGGATAGG | PjRGF7 antisense #1 |
| oMRF1512 | CTTGTCGTCTCTTGTGGTAGTG | PjRGF8 sense #1 |
| oMRF1513 | CTGCCATGTCTTTGTGGAGTA | PjRGF8 antisense #1 |
| oMRF1514 | CAGTCGACGTCTAAATCGGATAC | PjRGF8 sense #2 |
| oMRF1515 | CTTGCTCTCCTCTCTTCTTGTG | PjRGF8 antisense #2 |
| oMRF1516 | CGCGGTTAAGAAGACGAGTAA | PjRGF10 sense #1 |
| oMRF1517 | GCGCTGAAAGCCATGAAATAG | PjRGF10 antisense #1 |
| oMRF1518 | CGTATGCTCGTCAAGTAAAGGA | PjRGF10 sense #2 |
| oMRF1519 | TCGAACAGCATCAGGTTGAG | PjRGF10 antisense #2 |
| oMRF1520 | GATCCACAAGCTACCCTTACAA | PjRGF11 sense #1 |
| oMRF1521 | TCAGGTTTGCCACCTCATC | PjRGF11 antisense #1 |
| oMRF1522 | AAATGTTGCAAGTGGTTGATGA | PjRGF11 sense #2 |
| oMRF1523 | GAGCTTGTAAGGGTAGCTTGT | PjRGF11 antisense #2 |
| oMRF1524 | GCTGCCAAGGAAACTTAAACTC | PjRGF12 sense #1 |
| oMRF1525 | CCATTCTCTCCATGTCCCTTTAT | PjRGF12 antisense #1 |
| oMRF1526 | GCAAAGGTCACTCTGCCTAAT | PjRGF12 sense #2 |
| oMRF1527 | CGCGCGGGACAATATCTTT | PjRGF12 antisense #2 |
| oMRF1528 | ACCCGGTACTGCTTCAAATC | PjRGF17 sense #1 |
| oMRF1529 | TTCCTTGGCAGTGAAGTTAGAG | PjRGF17 antisense #1 |
| oMRF1530 | CTTCACTGCCAAGGAAACTTAAAC | PjRGF17 sense #2 |
| oMRF1531 | TGAGGGTTTGATCAGCTTCTTC | PjRGF17 antisense #2 |
| oMRF1539 | TCGAAGTAGTGATTGATTATCTGGCGATATCCCAGGTTTTAGAGCTAGAAATAGCAAGTTAAAATAAGG | PjRGFR4 gRNA4 fwd for pGTR w/ pMRF345 U6 pro overlap |
| oMRF1540 | AACTTGCTATTTCTAGCTCTAAAACCTGGGATATCGCCAGATAATTGCACCAGCCGGGAATCgaa | PjRGFR4 gRNA4 rev for pGTR w/ pMRF345 gRNA scaffold overlap |
| oMRF1541 | AACTTGCTATTTCTAGCTCTAAAACCCCAATATCTGCCGGCACGTTGCACCAGCCGGGAATCgaa | PjRGFR4 gRNA18 rev for pGTR w/ pMRF345 gRNA scaffold overlap |
| oMRF1545 | TCGAAGTAGTGATTGACAGCGAAAACAGGGGACCGGTTTTAGAGCTAGAAATAGCAAGTTAAAATAAGG | PjRGFR1 gRNA6 fwd for pGTR w/ pMRF345 U6 pro overlap |
| oMRF1546 | AACTTGCTATTTCTAGCTCTAAAACCGGTCCCCTGTTTTCGCTGTTGCACCAGCCGGGAATCgaa | PjRGFR1 gRNA6 rev for pGTR w/ pMRF345 gRNA scaffold overlap |
| oMRF1547 | AACTTGCTATTTCTAGCTCTAAAACTGCAGTTCTGTCCAATGCCATGCACCAGCCGGGAATCgaa | PjRGFR1 gRNA7 rev for pGTR w/ pMRF345 gRNA scaffold overlap |
| oMRF1551 | TCGAAGTAGTGATTGGACAAGAACCCACCACCGAAGTTTTAGAGCTAGAAATAGCAAGTTAAAATAAGG | PjRGF2 gRNA1 fwd for pGTR w/ pMRF345 U6 pro overlap |
| oMRF1552 | AACTTGCTATTTCTAGCTCTAAAACAACGGCGCATTATGAATCTCTGCACCAGCCGGGAATCgaa | PjRGF2 gRNA6 rev for pGTR w/ pMRF345 gRNA scaffold overlap |
| oMRF1584 | tgtgccgaattcggatccggtggtattgctacagttgccgg | PjRGF1 promoter fwd for infusion of pMRF295 |
| oMRF1585 | cctcgcccttgctcacCATTgatgaaccaccacactaattcttttag | PjRGF1 promoter rev for infusion of pMRF295 |
| oMRF1620 | ATGGCCTCACTTGTTTATCTTGTTTTATATCTC | PjRGF13 fwd |
| oMRF1621 | ATTACATCTAGGTCCCCCTGTTATGG | PjRGF13 rev |
| oMRF1624 | CAGAAAGGTCGACACTGGAATC | PjRGF13 qPCR sense |
| oMRF1625 | GAGGCAGCGCATAATCCATAA | PjRGF13 qPCR antisense |
| oMRF1654 | TAGAGTCGAAGTAGTGATTGACGTCGACAAAATCACCCTGGTTTTAGAGCTAGAAATAGCAAGTTAAAAT | PjRGF3 gRNA1 fwd for pGTR w/ pMRF345 U6 pro overlap |
| oMRF1655 | AACTTGCTATTTCTAGCTCTAAAACCGTGGCTCGATCTCGTCTGCTGCACCAGCCGGGAATC | PjRGFR3 gRNA3 rev for pGTR w/ pMRF345 gRNA scaffold overlap |
| oMRF1656 | TAGAGTCGAAGTAGTGATTGTGTAAAGCGACAAAACCTGGTTTTAGAGCTAGAAATAGCAAGTTAAAAT | PjRGFR1 gRNA2 fwd for pGTR w/ pMRF345 U6 pro overlap |
| oMRF1724 | TCTGTTGTTGATGTGATTACAGaaatgaaacggacagccgacg | pFASTRK bp1 fwd for LjUBQ |
| oMRF1725 | TTGCTAaagatctcttggaggtagcagatcctg | pFASTRK bp1 reverse for zCas9i |
| oMRF1726 | ctccaagagatcttTAGCAACGAGATGG | zCas9i fwd for bp |
| oMRF1727 | CTCTGTGCTTGTTGTACGCGGAAAG | zCas9i rev for bp |
| oMRF1728 | CGCGTACAACAAGCACAGAGataagcctatcagg | pFASTRK bp2 fwd for Zcas9i |
| oMRF1729 | GTTTGAACGATCTGCTTGACAAGCctaggatccacctgaaaccttcctc | pFASTRK bp2 reverse for nosT |

Supplemental Table S7. Plasmids and strains used in this study

| Name | Strain | Plasmid description | Antibiotic Resistance | Reference |
|---|---|---|---|---|
| pMRF121 | DH5a | pAGM4723::LjUBQpro_3xmCherry-syp122-Pjv1_00001310pro_3xVenus-N7 | Km | This study |
| pMRF122 | DH5a | pAGM4723::LjUBQpro_3xmCherry-syp122-Pjv1_00011762pro_3xVenus-N7 | Km | This study |
| pMRF139 | E. coli DH5a | pAGM4723::LjUBQ_mCherry-syp122_PjPLTpro_mNG-PjPLT | Km | This study |
| pMRF143 | E. coli DH5a | pAGM4723::LjUBQpro_3xmCherry-syp122-PjRGFR5pro_3xVenus-N7 | Km | This study |
| pMRF144 | E. coli DH5a | pAGM4723::LjUBQpro_3xmCherry-syp122-PjRGFR2pro_3xVenus-N7 | Km | This study |
| pMRF145 | E. coli DH5a | pAGM4723::LjUBQpro_3xmCherry-syp122-PjRGFR6pro_3xVenus-N7 | Km | This study |
| pMRF148 | E. coli TOP10 | pAGM4723::LjUBQpro_3xmCherry-syp122-PjRGFR1pro_3xVenus-N7 | Km | This study |
| pMRF149 | E. coli TOP10 | pAGM4723::LjUBQpro_3xmCherry-syp122-PjRGF2pro_3xVenus-N7 | Km | This study |
| pMRF163 | E. coli DH5a | pICH41308::RUBY | Sp | This study |
| pMRF217 | E. coli DH5a | ePiGreen:Pjv1_00019566ecto-BRI1kinase-3xHA | Km | This study |
| pMRF218 | E. coli DH5a | ePiGreen:Pjv1_00010527ecto-BRI1kinase-3xHA | Km | This study |
| pMRF219 | E. coli DH5a | ePiGreen:Pjv1_00020079ecto-BRI1kinase-3xHA | Km | This study |
| pMRF220 | E. coli DH5a | ePiGreen:Pjv1_00003866ecto-BRI1kinase-3xHA | Km | This study |
| pMRF221 | E. coli DH5a | ePiGreen:Pjv1_00011762ecto-BRI1kinase-3xHA | Km | This study |
| pMRF222 | E. coli DH5a | ePiGreen:Pjv1_00001310ecto-BRI1kinase-3xHA | Km | This study |
| pMRF223 | E. coli DH5a | ePiGreen:Pjv1_00007151ecto-BRI1kinase-3xHA | Km | This study |
| pMRF224 | E. coli DH5a | pAGM4723::PjPLT-3xVenus_LjUBQpro-19181sp-mScarlet-19181GPI | Km | This study |
| pMRF295 | E. coli DH5a | pAGM4723::bsmbI-3xVenusNLS-hspT_35S-19181sp-mScarlet-19181GPI-nosT | Km | This study |
| pMRF296 | E. coli DH5a | pAGM4723::bsmBI-bsmBI-hspT_35S-3xVenusNLS-nosT | Km | This study |
| pMRF354 | E. coli DH5a | pAGM4723::U6-26pro-BsmBIcel5-gRNAscaffold_LjUBQ-Zcas9i-nosT_35S-RUBY-nosT | Km | This study |
| pMRF357 | E. coli DH5a | pMRF354::PjRGFR4gRNAs | Km | This study |
| pMRF358 | E. coli DH5a | pMRF354::PjRGFR1gRNAs | Km | This study |
| pMRF363 | DH5a | pMRF295::PjRGF1pro | Km | This study |
| pMRF365 | E. coli DH5a | pMRF354::PjRGF2gRNAs | Km | This study |
| pMRF366 | E. coli DH5a | pMRF354::PjRGFR3gRNAs | Km | This study |
| pMRF374 | E. coli DH5a | pAGM4723::U6-26pro-BsmBIcel5-gRNAscaffold_LjUBQ-bp-Zcas9i-bp-nosT_35S-RUBY-NI | Km | This study |
| pMRF379 | E. coli DH5a | pMRF374::PjRGFR1gRNAs | Km | This study |
| | E. coli DH5a | pGWB4::35S-RUBY-nosT | Km | This study |
| aMRF066 | A. rhizogenes AR1193 | pMRF121 | Strep, Cb, Rif, Km | This study |
| aMRF067 | A. rhizogenes AR1193 | pMRF122 | Strep, Cb, Rif, Km | This study |
| aMRF084 | A. rhizogenes AR1193 | pMRF139 | Strep, Cb, Rif, Km | This study |
| aMRF088 | A. rhizogenes AR1193 | pMRF143 | Strep, Cb, Rif, Km | This study |
| aMRF089 | A. rhizogenes AR1193 | pMRF144 | Strep, Cb, Rif, Km | This study |
| aMRF090 | A. rhizogenes AR1193 | pMRF145 | Strep, Cb, Rif, Km | This study |
| aMRF095 | A. rhizogenes AR1193 | pMRF149 | Strep, Cb, Rif, Km | This study |
| aMRF096 | A. rhizogenes AR1193 | pGWB4::35S-RUBY-nosT | Strep, Cb, Rif, Km | This study |
| aMRF115 | A. rhizogenes AR1193 | pMRF179 | Km, Sp, Rif | This study |
| aMRF146 | A. tumafaciens AGL1 | pMRF217 | Km, Cb | This study |
| aMRF147 | A. tumafaciens AGL1 | pMRF218 | Km, Cb | This study |
| aMRF148 | A. tumafaciens AGL1 | pMRF219 | Km, Cb | This study |
| aMRF149 | A. tumafaciens AGL1 | pMRF220 | Km, Cb, Rif | This study |
| aMRF150 | A. tumafaciens AGL1 | pMRF221 | Km, Cb | This study |
| aMRF151 | A. tumafaciens AGL1 | pMRF222 | Km, Cb | This study |
| aMRF160 | A. rhizogenes AR1193 | pMRF224 | Km, Sp, Rif | This study |
| aMRF241 | A. rhizogenes AR1193 | pMRF357 | Km, Rif | This study |
| aMRF242 | A. rhizogenes AR1193 | pMRF358 | Km, Rif | This study |
| aMRF250 | A. rhizogenes AR1193 | pMRF365 | Km, Rif | This study |
| aMRF251 | A. rhizogenes AR1193 | pMRF366 | Km, Rif | This study |
| aMRF257 | A. rhizogenes AR1193 | pMRF379 | Km, Rif | This study |
| pMLBART | A. rhizogenes AR1193 | DR5::3xVenus-N7 | Sp, Rif | 68 |
| | A. tumafaciens AGL1 | ePiGreen::EFRecto-BRI1kinase-3xHA | Km | 36 |
| | A. tumafaciens AGL1 | ePiGreen::BES1-3xHA | Km | 36 |
